# Integrated Framework for Probing Multimodal Protein Foundation Models with Structure-Functional Interpretability Analysis in Detection of Allosteric Binding Sites

**DOI:** 10.64898/2026.07.02.736203

**Authors:** Alina Bazarova, Gennady M. Verkhivker

## Abstract

Allosteric regulation represents a fundamental mechanism of protein function, yet distinguishing allosteric from orthosteric protein binding sites remains a persistent computational challenge. While multimodal protein foundation models offer the potential to integrate complementary biological signals including sequence, structure, functional annotations, and conformational dynamics, their performance determinants in allosteric binding site detection remain poorly understood. We introduce a unified computational framework for profiling multimodal protein foundation models across distinct binding-site separability regimes. Rather than evaluating models solely by predictive accuracy, the framework combines systematic modality embedding ablations, encoder architecture comparisons, and variance decomposition to characterize how evolutionary, structural, functional, and dynamical information contribute to allosteric site discrimination. Using the OneProt multimodal model, we evaluate two complementary levels of multimodal integration: (a) encoder architectures that differ in the modalities incorporated during pretraining, and (b) downstream combinations of pocket, sequence, and text embeddings used for classification. To systematically probe the determinants of model performance, we benchmark these configurations across four assembled datasets of protein complexes representing a spectrum of biological complexity and a range of structural, dynamic, and evolutionary context for orthosteric and allosteric binding sites. Through comprehensive embedding ablations, encoder architecture comparisons, and variance decomposition, we demonstrate that model performance is governed primarily by intrinsic dataset properties rather than architectural complexity, with dataset identity accounting for 63.7% of explainable variance. Across all examined datasets, we identify three distinct separability regimes: a low-separability regime where current representations fail to reliably distinguish the two classes; an intermediate regime where multimodal integration substantially improves performance; and a high-separability regime where most architectures converge to near-ceiling performance. Critically, embedding contributions are regime-dependent: pocket geometry dominates when regulatory classes share structural contexts, while text and sequence embeddings become essential when evolutionary constraint distinguishes them. At the encoder level, structural and molecular dynamics encoders provide the greatest benefit in intermediate- and high-separability settings. Structure-functional analysis of correctly classified binding sites reveals that prediction success reflects the underlying biological organization of each regime. These findings establish that the success of multimodal foundation models depends critically on alignment between available modalities and the biological signatures that distinguish regulatory classes in each dataset.

## Introduction

The discovery of druggable binding sites in proteins—particularly those that are cryptic, transient, or allosteric—remains one of the most persistent bottlenecks in structure-based drug design and chemical biology.^1–4^ Allosteric sites, in particular, offer tremendous therapeutic potential due to their capacity for exquisite selectivity, tunable activity control, and resistance evasion, yet they remain notoriously difficult to identify computationally. Unlike orthosteric ligands, which directly compete with endogenous substrates at catalytic or active sites, allosteric modulators influence protein behavior indirectly through long-range conformational coupling and regulatory communication within the protein structure.^5–8^ Because allosteric modulation can provide increased selectivity, tunable activity control, and reduced toxicity, allosteric drug discovery has emerged as an important strategy in modern pharmacology and chemical biology.^9^ Orthosteric pockets are often evolutionarily conserved and structurally well defined, whereas allosteric sites tend to exhibit greater heterogeneity in their geometry, physicochemical composition, and spatial relationship to functional regions.^10–13^ In many proteins, allosteric regulation is strongly coupled to conformational dynamics and transient structural states that are not fully captured by static structural descriptions alone.^14^ Consequently, distinguishing allosteric binding sites from orthosteric binding requires integrating information across multiple biological scales, including sequence conservation, local pocket geometry, functional annotations, and conformational variability. Computational strategies for binding site prediction have evolved from geometric and energetic approaches such as Fpocket,^15^ SiteHound,^16^ and MetaPocket 2.0^17^ to machine learning (ML) methods such as P2Rank^18,19^ which predicts binding site residues by scoring local chemical neighborhoods using a random forest classifier trained on known protein–ligand complexes. Deep learning (DL) has further advanced the field, with voxel-based approaches such as DeepSite^20^ object detection architectures exemplified by FRSite^21^ segmentation networks,^22^ and dynamics-aware models BiteNet^23^ achieving high accuracy on benchmarks. However, these structure-based methods exhibit performance drops when predicting binding sites in the ligand-bound holo complexes using apo structures and identifying cryptic sites that emerge during dynamic changes revealing their limitations in allosteric site detection, where conformational heterogeneity and transient pocket formation are important features underpinning allosteric regulatory events.^24–26^

Recent advances in protein foundation models provide a promising framework for integrating complementary signals across biological scales. Protein language models (PLMs) have demonstrated that evolutionary-scale representations encode substantial information about protein structure and function.^27,28^ Trained on hundreds of millions of sequences, PLMs learn evolutionary constraints and structural propensities directly from primary sequence, enabling scalable proteome-wide predictions of binding sites and ligand binding without explicit structural input.^29–31^ PLM fine-tuning strategies can adapt model parameters during task-specific training which can be achieved through full fine-tuning, in which all transformer layers are updated^32^ or via parameter-efficient fine-tuning (PEFT) methods that introduce a small set of trainable parameters while freezing the original PLM weights.^33,34^ A range of fine-tuning protocols for the prediction of cryptic binding sites was evaluated using benchmarking against frozen-embedding baselines revealing that sequence-based models could be enhanced by combining domain-informed fine-tuning with multi-task learning and high-quality curated datasets.^35^ However, the reliance on evolutionary conservation can often render PLMs to become less robust in detection of allosteric and cryptic sites, which are often evolutionarily labile and lack deep sequence conservation.^31,35^ Recent approaches have fused PLM embeddings with structural graphs (EquiBind,^36^ DiffSBDD,^37^ dynamics simulations (PocketMiner^38^), and physics-informed strategies (AlloPred,^39^ Allosite,^40^ DeepAllo^41,42^).

More recently, multimodal protein foundation models have extended this paradigm by jointly incorporating sequence, structural, biochemical, textual, and dynamical information into shared latent embedding spaces learned through self-supervised or contrastive pretraining.^43–47^ By aligning multiple biological modalities, these models aim to capture complementary aspects of protein organization not accessible from sequence or structure alone. A prominent example is OneProt model^48^ which integrates (a) sequence by using ESM2 model, (b) structural information through Foldseek^49^ tokens and ProNet graphs,^50^ (c) binding pockets via ProNet, and (d) text annotations with BiomedBERT^51^ through contrastive alignment to a sequence anchor. An extended version further incorporates molecular dynamics (MD) trajectories via a transformer-based MDGen encoder,^52^ capturing protein flexibility showing that MD inclusion alters latent-space organization and produces distinct downstream transfer characteristics.

Despite this progress, robust detection of functional allosteric and cryptic binding sites remains elusive as most methods are optimized for stable, evolutionarily conserved orthosteric pockets, whereas allosteric sites are sparsely populated in structural databases.^53^ A recent investigation integrated local binding geometry, coevolution, and dynamic allostery into a three-parameter model to identify potentially hidden allosteric sites in ensembles of protein structures with orthosteric ligands.^54^ The performance dichotomy raises a fundamental question: what physical principles govern the evolutionary design of allosteric sites, and how can we leverage these principles to build more interpretable predictive models? To systematically interrogate the physical determinants of binding site predictability, we have recently developed a dual analytical framework that integrates a fine-tuned PLM with a physics-based energy landscape-centric interpretability layer derived from energy landscape theory and frustration analysis approach.^55–62^ Applying this framework to human kinases, we demonstrated that orthosteric and allosteric sites occupy fundamentally different energetic regimes where orthosteric ATP-binding sites can be reliably identified with high precision, whereas allosteric sites are detected with substantially lower confidence across kinase families and conformational states.^31^ This analysis revealed that predictive success is governed by the local energetic embedding of binding sites: orthosteric pockets are located in minimally frustrated basins that generate strong evolutionary and structural signatures, whereas allosteric pockets occupy predominantly neutrally frustrated zones associated with conformational plasticity and reduced evolutionary constraint.^31^ Most existing studies focus on maximizing predictive performance on individual benchmarks, while comparatively little attention has been given to how intrinsic dataset properties influence the learnability of allosteric regulation. In particular, it remains unclear how structural diversity, conformational heterogeneity, functional annotation coverage, and class imbalance affect the relative contribution of different biological modalities. To address this gap, we develop a unified analytical framework for systematically profiling multimodal protein foundation models across distinct binding-site separability regimes. The framework enables systematic dissection of how different biological modalities contribute to allosteric-site discrimination across diverse biological contexts. Using the OneProt multimodal framework, we systematically evaluate both encoder architectures, which differ in the biological modalities incorporated during multimodal pretraining, and downstream combinations of pocket, sequence, and text embeddings extracted from the corresponding pretrained OneProt encoders across multiple datasets containing experimentally annotated allosteric and orthosteric binding sites. Rather than treating allostery prediction as a single benchmark problem, we interpret model behavior across four designed datasets (PPI-Site, KinSite, Dual-Site and AlloDiverse) reflecting distinct biological regimes of separability. PPI-Site dataset comprises binding sites annotated by their spatial relationship to protein–protein interaction interfaces, where allosteric and orthosteric sites often share the same structural context. KinSite dataset contains kinase inhibitor complexes, where orthosteric ATP-binding sites are evolutionarily conserved while allosteric sites are structurally heterogeneous. Dual-Site dataset combines PPI-Site and KinSite to evaluate for synergistic effects. AlloDiverse dataset incorporates allosteric sites from diverse protein families to assess performance under high-signal, high-diversity conditions. We investigate how modality composition, structural diversity, conformational variability, and class imbalance shape predictive performance and representation quality across different allosteric learning regimes. Our results reveal that model performance is governed primarily by intrinsic dataset properties rather than architectural complexity alone. Across datasets, we identify three distinct regimes: (a) low-separability settings, where current representations underperform to reliably distinguish allosteric and orthosteric pockets; (b) intermediate regimes, where multimodal integration substantially improves ranking and sensitivity; and (c) highly separable regimes, where most architectures converge to near-ceiling performance. We further show that the contributions of pocket, sequence, and text embedding combinations, as well as those of structural and MD encoder architectures, depend on dataset composition and structural diversity, highlighting the context-dependent role of multimodal protein representations in modeling allosteric regulation. Structure-functional interpretability analysis of recovered true positive (allosteric) and true negative (orthosteric) predictions reveals that prediction success reflects the underlying structural and functional organization of binding sites. We find that in PPI-Site dataset, allosteric and orthosteric sites often share identical residue compositions, differing only in conformational arrangement relative to protein–protein interfaces, explaining why pocket geometry alone provides the only discriminative signal. In KinSite, orthosteric ATP-binding clefts show strong sequence conservation while allosteric regulatory loops exhibit weak conservation, creating differential evolutionary signatures that sequence and text embeddings capture most effectively. In the combined Dual-Site setting, structural diversity from PPI-Site regularizes the model, improving recognition of rare allosteric kinase sites, while evolutionary signals from KinSite resolve ambiguous PPI-Site cases. These findings establish that the success of multimodal foundation models depends critically on alignment between available modalities and the biological signatures that distinguish regulatory classes in each dataset. Through this framework, we provide a systematic characterization of when and why multimodal integration benefits allostery prediction. By linking predictive performance to underlying biological and statistical properties and grounding our analysis in physical principles, this study establishes a foundation for developing more robust, context-aware, and physically informed models for allosteric site discovery.

## Materials and Methods

### General Formulation and Multimodality Downstream Prediction Pipeline

In this study, we investigate the discrimination of allosteric and orthosteric competitive binding sites using multimodal protein foundation model representations. Unlike conventional ligand-binding prediction tasks, our objective is not to identify whether a protein region binds ligands, but rather to distinguish between distinct functional and regulatory binding regimes assuming that the binding site location is known a priori.

Formally, let a protein *P* be represented by its amino acid sequence *S* ∈ *A^L^*, where *A* denotes the alphabet of 20 standard amino acids and *L* is the sequence length. Associated with *P* is a three-dimensional structure *X ∈ R^Nx3^* comprising *N* atoms, and a set of functional annotations *T* derived from curated biological databases. Where available, conformational dynamics are represented by an MD trajectory *M ∈ R^Nx3xτ^*, where τ denotes the number of trajectory frames. For a given binding site *B*, we define its spatial extent as a set of *K* = 100 residues *R_B_ = {r_1_, r_2_…,r_K_}* proximal to the bound ligand. During multimodal pretraining, the structural modality *X* may be encoded either through graph-based structural representations *X_SG_* or through structure-token representations *X_ST_*, depending on the selected OneProt encoder architecture.^48^

The downstream task is formulated as binary classification, where experimentally annotated allosteric pockets are treated as positive instances *y = 1* and orthosteric competitive pockets as negative instances *y = 0*. Let the encoder input space be defined as

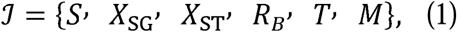

For a given OneProt encoder architecture, only a subset 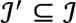 is used during multimodal pretraining. The pretrained is therefore represented as

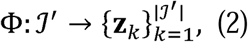

where each ***z****_k_ ∊ R^d^* is a modality-specific embedding vector. For downstream allostery prediction, only the pocket ***z****_pocket_*, sequence ***z****_seq_*, and text ***z****_text_* embeddings are considered.

Depending on the embedding ablation setting, these embeddings are concatenated to form 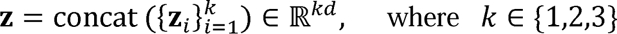 depending on the selected embeddingss combination. The downstream classifier *f:R^kd^ →[0,1]*; predicts the probability *ŷ = P(y = 1 | **z**)*.

Rather than using OneProt solely as a predictive classifier, we interpret multimodal representation behavior across datasets as a means of probing which biological modalities contribute to allosteric discrimination under different structural and statistical conditions.

The full multimodal representation pipeline, illustrated in Figure 1, proceeds through four integrated phases. First, we combine three complementary datasets. The PPI-Site dataset was derived from a large-scale pocket-centric collection of protein–ligand binding sites and protein–protein interaction (PPI) interfaces^63^ derived from experimentally resolved Protein Data Bank^64^ structures. The KinSite dataset was assembled from multiple structural and activity resources, including PDB,^64^ KLIFS database^65^ KinLigDB^66^ (that provides structures of 4219 kinase-ligand complex structures covering 297 human and the Kinase Conformation Resource (KinCoRe).^67,68^ The Allosteric Database (ASD) datset^69^ includes allosteric ligand-protein complexes that includes diverse protein families, including kinases, phosphatases, GPCRs, ion channels, and transcription factors.

**Figure 1.**
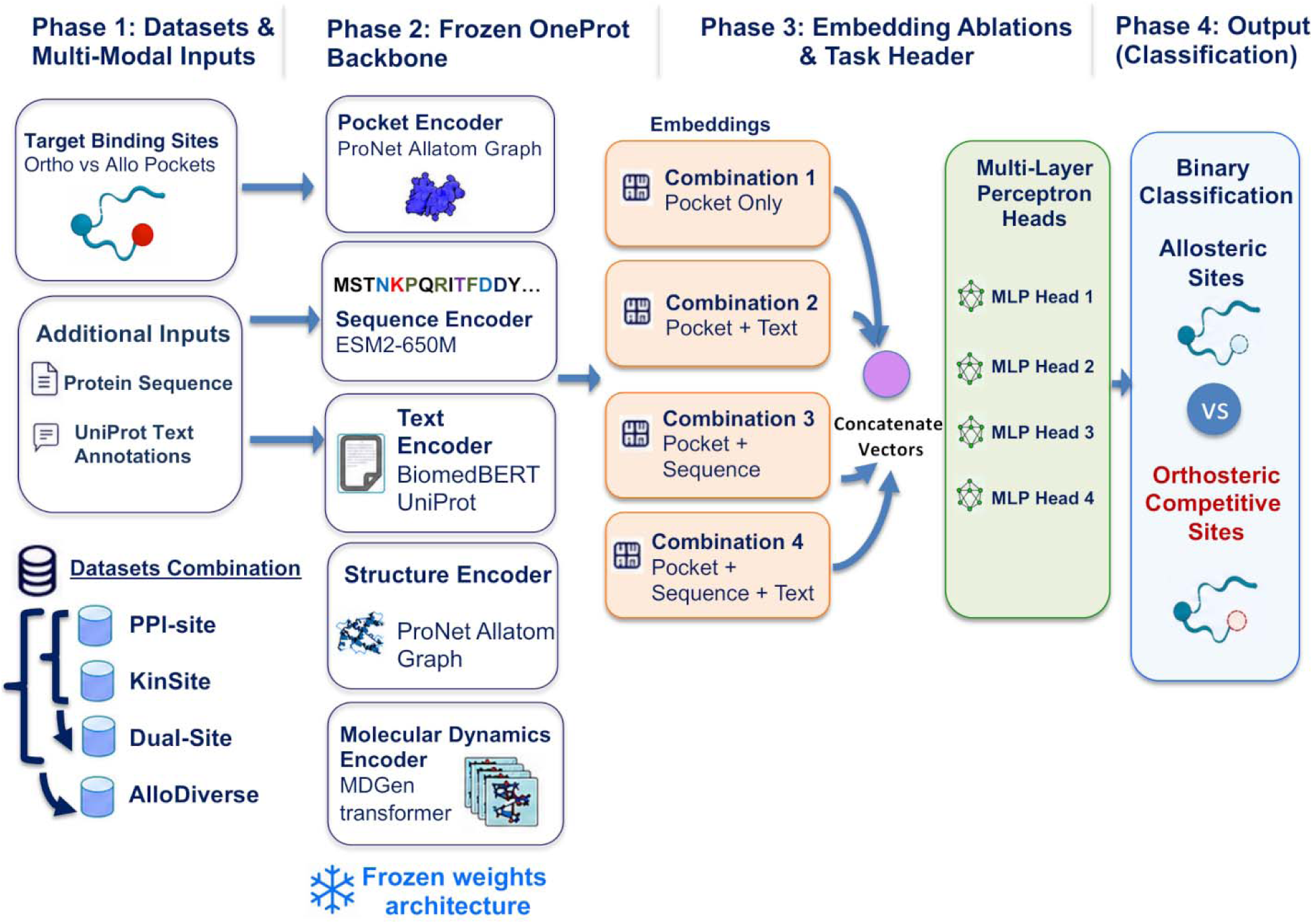
Analytical framework for investigating multimodal representations for binding site detection. The study is organized into four interconnected stages. Phase 1 defines the allosteric-site prediction task and introduces four benchmark datasets—PPI-Site, KinSite, Dual-Site, and AlloDiverse—that span distinct biological regimes and collectively define a progression from low to high separability. Phase 2 employs a frozen OneProt multimodal backbone comprising pocket, sequence, text, structure, and molecular-dynamics encoders to generate protein representations. Phase 3 systematically evaluates two levels of multimodal integration: encoder architectures that differ in the modalities incorporated during pretraining (ST, SG, MD variants) and downstream embedding combinations (pocket-only, pocket+text, pocket+sequence, pocket+sequence+text). Phase 4 performs binary classification of allosteric versus orthosteric competitive sites and enables quantitative comparison of predictive performance across datasets, architectures, and embedding combinations, providing a systematic assessment of how dataset composition, modality selection, and architectural design influence allosteric-site prediction.

For each binding pocket, we assemble the input modalities required by the frozen pretrained OneProt encoders to generate modality-specific embeddings for downstream fine-tuning: (a) the protein sequence *S* (amino acid string), (b) the three-dimensional pocket structure *X* (atomic coordinates of the *z = 100* residues surrounding the binding site), and (c) UniProt^70^ text annotations *T* (including protein names, protein family assignments, organism information, functional descriptions, pathway annotations, subcellular localization). This multi-modal input design is critical because allostery is encoded not only in sequence conservation and geometric pocket shape, but also in the functional context and evolutionary history of the protein. We use pretrained OneProt encoder architectures^48^ to generate modality-specific embeddings for downstream allostery classification. Throughout this study, the pretrained encoders remain frozen, while only a lightweight multilayer perceptron (MLP), taking the selected embedding combinations as input, is trained on the allosteric versus orthosteric classification task using a binary cross-entropy loss. The downstream classifier receives only combinations of the pocket, sequence, and text embeddings extracted from the pretrained encoders. However, the pretrained OneProt architectures differ in the encoder modalities incorporated during multimodal pretraining. In particular, some architectures additionally include structure-token, structure-graph, and MD encoders. Although embeddings from these additional encoders are not used directly for downstream classification, they influence the shared latent representation learned during pretraining and thereby shape the resulting pocket, sequence, and text embeddings. This design enables us to distinguish the contribution of multimodal pretraining from that of the embedding combinations used by the downstream classifier. For each modality, a dedicated encoder extracts task-relevant features. Pocket Encoder (ProNet Atom Graph) is a graph neural network operating on the atomic graph of the binding pocket, capturing local geometric and chemical features including atom types, bond distances, and residue contacts. Sequence Encoder (ESM2 Transformer) is a transformer-based encoder that processes the amino acid sequence, capturing evolutionary constraints, co-evolutionary signals, and context-dependent residue relationships;. Text Encoder (BiomedBERT UniProt) is a biomedical language model fine-tuned on UniProt text annotations, extracting semantic information from functional descriptions, domain annotations, and pathway information. Optional encoders include Structural Encoders (ProNet Atom Graph + ESM2 Transformer). The ProNet encoder^50^ processes the protein structure as an atom graph, whereas the ESM2 transformer operates on Foldseek-derived structure tokens. Together, these complementary structural encoders provide alternative representations of the same three-dimensional protein structure, whose outputs are integrated during multimodal pretraining. MD Encoder (MDGen transformer encoder) is an encoder that processes trajectory data, capturing dynamical features such as residue fluctuations and conformational ensembles. The outputs of the modality-specific encoders are projected into a shared latent space during multimodal pretraining. The pretrained OneProt backbone is kept frozen throughout this study, allowing us to evaluate how different encoder architectures shape the learned latent representation without modifying it during downstream training. Consequently, structural, and MD encoders influence downstream performance indirectly by affecting the latent space from which the pocket, sequence, and text embeddings are extracted.

To dissect the contribution of each biological modality to allosteric discrimination, we perform a systematic embedding ablation analysis. We train separate classifiers on four distinct embedding combinations: Combination 1: Pocket only (local structural features alone); Combination 2: Pocket + Text (local structure + functional annotations) Combination 3: Pocket + Sequence (local structure + evolutionary sequence patterns) Combination 4: Pocket + Sequence + Text (trimodal integration). This ablative design allows us to quantify the marginal contribution of each modality to classification performance. We interpret multimodal representation behavior across datasets as a means of probing which biological modalities contribute to allosteric discrimination under different structural and statistical conditions. By systematically ablating input modalities and comparing classification performance across datasets, we can ask: Which signals—local structural geometry, evolutionary sequence constraints, or functional annotations—are most diagnostic of allosteric sites? This diagnostic framing transforms the classification task from a black-box prediction problem into a mechanistic investigation of allosteric site encoding in biological sequence and structure space.

### Datasets and Biological Regimes

We evaluated the proposed framework on three complementary datasets containing experimentally annotated allosteric and orthosteric binding sites. Rather than serving solely as independent benchmarks, the selected datasets were deliberately chosen to represent distinct structural, functional, and statistical regimes, enabling systematic investigation of how dataset composition influences allosteric discriminability and modality-dependent representation behavior. Where predefined train/validation/test partitions were not available, proteins were clustered using MMseqs2^59^ at 30% sequence identity prior to dataset splitting to ensure that homologous proteins did not appear across splits. Where training/validation splits were available (ASD dataset), we inherited those while randomly splitting the original validation set into two equally sized validation and test sets.

### PPI-Site Dataset

The PPI-Site dataset^63^ was constructed through a multi-stage filtering pipeline incorporating experimental quality constraints, including X-ray structures with resolution ≤3.5OÅ, ligand filtering, and interface-aware classification of binding pockets. Binding pockets were identified using VolSite.^71^ A defining feature of the PPI-Site dataset is that binding pockets are classified according to their spatial relationship to PPI interfaces, yielding three distinct types: PLOC (Protein-Ligand Orthosteric Competitive) where the ligand pocket directly overlaps with the PPI epitope; PLONC (Protein-Ligand Orthosteric Non-Competitive) where the ligand pocket is near but non-overlapping with the PPI interface; and PLA (Protein-Ligand Allosteric) where the ligand pocket is spatially distinct from the PPI interface. In this study, PLOC pockets were treated as negative instances (orthosteric) and PLA pockets as positive instances (allosteric); PLONC pockets were excluded to maintain a clear binary distinction between direct competitive binding and allosteric modulation. Because PPI-Site class labels arise from spatial relationships rather than independent regulatory mechanisms, allosteric and orthosteric pockets often share closely related structural and functional contexts. This makes PPI-Site a low-separability regime, where discrimination depends primarily on subtle conformational differences rather than distinct structural architectures. PPI-Site contains structurally consistent pocket annotations across a broad range of protein families with moderate class imbalance, comprising 1,845 PLOC and 1,689 PLA sites across 1,278 unique protein chains spanning 847 CATH families.

### KinSite Dataset

In the KinSite dataset, KinLigDB^66^ contains 4,219 curated kinase-ligand complex structures of 297 human kinases from 106 kinase families with ∼ 75% of these kinases having at least two structures with different ligands, and 92 kinases having more than 10 complex structures. For this database, there are reported type I inhibitors (3167 entries), type II inhibitors (283 entries), type III inhibitors (23 entries), type IV inhibitors (9 entries), competitive inhibitors (554 entries), covalent inhibitors (31 entries), activators (49 entries), and other allosteric ligands (103 entries).^66^ KinLigDB entries are combined with KinCoRe^67,68^ which classifies inhibitors into five mechanistically distinct classes. Type I inhibitors are ATP-competitive and orthosteric, binding to the DFG-in, αC-in conformation. Type I.5 inhibitors are ATP-competitive and orthosteric, binding to the DFG-in, αC-out conformation, extending into the back pocket. Type II inhibitors are ATP-competitive and orthosteric, binding to the DFG-out conformation with back pocket extension. Type III inhibitors are allosteric and proximal, binding >6.0OÅ from the hinge region. Type IV inhibitors are allosteric and distal, binding >6.5OÅ from both the hinge region, Types I, I.5, and II were treated as negative instances (orthosteric), while Types III and IV were treated as positive instances (allosteric). Kinase ATP-binding sites are highly conserved generating strong evolutionary signals whereas allosteric kinase sites are structurally heterogeneous. This creates an intermediate-separability regime where sequence information becomes essential for discrimination. After rigorous filtering (temporal: structures deposited after 2020 only; sequence: ≥30% sequence identity removed; pocket-level: ≥40% residue identity to training pockets removed; quality: resolution ≤3.0OÅ; occupancy: ligand atoms with occupancy <0.5 excluded), the KinSite dataset comprises 3,363 orthosteric and 172 allosteric high-confidence complexes spanning 246 kinases.

### ASD Dataset

Unlike PPI-Site and KinSite, ASD primarily provides residue-level annotations rather than predefined pocket structures. To obtain representations compatible with OneProt, residue-level annotations were converted into fixed-size structural pocket representations through spatial clustering and geometric reconstruction. ASD introduces substantially greater allosteric diversity, spanning diverse protein families, regulatory mechanisms (including both orthosteric and allosteric modulation), and evolutionary contexts. This creates a high-separability regime where the broad structural and functional heterogeneity of allosteric sites provides a strong discriminative signal. The ASD-derived dataset comprises 14,043 allosteric pocket instances across 1,842 unique proteins (spanning 247 CATH families and 531 Pfam families), substantially increasing the diversity of the positive class and enabling investigation of multimodal representation behavior under conditions of increased allosteric heterogeneity.

### Dataset Combinations and Regime Progression

To systematically investigate how dataset composition influences learnability, we constructed four evaluation configurations that systematically vary total sample size, allosteric diversity, and class balance, enabling controlled investigation of how dataset composition influences discriminability and modality-dependent behavior. The first configuration, uses the PPI-Site dataset alone, comprising 1,845 PLOC (negative) and 1,689 PLA (positive) sites, with a class imbalance ratio ρ = 1.09. This represents a low-separability regime where class discrimination depends on subtle conformational differences rather than distinct structural architectures.

The second configuration uses the KinSite dataset alone, comprising 3,363 orthosteric (negative) and 174 allosteric (positive) high-confidence complexes, with ρ = 51.1. This represents an intermediate-separability regime where sequence information becomes essential for discrimination, reflecting the conserved nature of kinase ATP sites contrasted with heterogeneous allosteric pockets. The third configuration Dual-Site combines PPI-Site and KinSite, yielding 5,203 orthosteric (negative) and 1,861 allosteric (positive) sites, with ρ = 2.79. This creates an intermediate-synergistic regime that balances the low-separability structural context of PPI-Site with the sequence-driven signal of KinSite. The fourth configuration AlloDiverse combines ASD with PPI-Site and KinSite, yielding 5,203 orthosteric (negative) and 15,904 allosteric (positive) sites, with ρ = 0.33. This represents a high-separability regime with extensive allosteric diversity and a positive-class majority, where the broad structural and functional heterogeneity of allosteric sites provides a strong discriminative signal. This progression from low to high separability enables controlled investigation of how dataset composition influences discriminability and modality-dependent behavior, with each configuration strategically positioned to probe a distinct aspect of the allosteric discrimination problem.

### Multimodal Backbone Architecture

We adopted the OneProt multimodal protein foundation model as the representation-learning framework for all downstream experiments.^48^ The conceptual role of OneProt in our downstream pipeline is described in the previous section outlining multimodality downstream prediction pipeline. Here, we briefly summarize the architectural components relevant to the present study. Complete details of the model architecture, multimodal pretraining objectives, and optimization procedure are available in the original OneProt model.^48^ Protein sequence representations are generated using the OneProt sequence encoder based on the ESM2-650M transformer architecture [43]. The encoder comprises 33 transformer layers with 20 attention heads per layer and a hidden dimension of 1280. Given an input protein sequence *S = (s_1_, s_2_…s_L_)*, the encoder produces contextualized residue-level representations

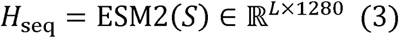

A protein-level sequence representation is obtained by mean pooling across residue positions,

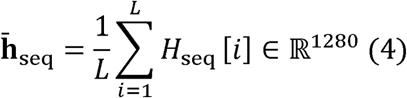

and is subsequently mapped into the shared OneProt latent space using a trainable projection head,

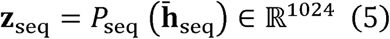

where *P_seq_* denotes the projection network and *dis* the dimension of the shared multimodal embedding space.

Structural representations were generated using two complementary encoders integrated within OneProt. The graph-based structural encoder uses ProNet^50^ which processes protein structures as hierarchical graphs through message passing across amino-acid, backbone, and all-atom levels to capture fine-grained structural details and higher-order geometric organization. The token-based structural encoder uses Foldseek structural tokens^49^ processed with an ESM2 transformer to discretize local three-dimensional environments into sequence-like tokens, capturing broader structural context and fold-level organization. Pocket embeddings were obtained using the binding-site encoder integrated within OneProt. Binding pockets are represented as three-dimensional residue graphs and processed using a graph neural network based on ProNet.^50^

For a pocket comprising *K = 100* residues with centroids 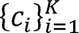, edges are established between residues *i* and *j*, when 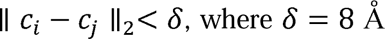. Through hierarchical message passing, the encoder captures localized chemical interactions and higher-order structural organization. The final pocket embedding is obtained by pooling across graph nodes.

Textual representations were derived from UniProt functional annotations [49] and processed using the BiomedBERT encoder,^51^ a domain-adapted BERT model pretrained on biomedical literature. Given a textual annotation *T = (w_1_, w_2_,…, w_M_)*, the encoder produces contextualized token-level representations, which are mean-pooled to obtain a fixed-dimensional text embedding.

The extended OneProt architecture incorporates MD trajectory representations using the latent MD encoder derived from MDGen^52^, a transformer-based generative framework that processes full MD trajectories as temporally ordered structural ensembles. MD trajectories used during pretraining were curated from mdCATH, GPCRmd, and ATLAS databases through MDRepo.^72^ Consistent with the ablation framework, MD embeddings were not directly concatenated into downstream classifier inputs; their contribution arises indirectly through multimodal latent-space shaping during pretraining.

During multimodal pretraining, modality-specific representations are projected into a shared latent space of dimension 1024 and aligned using a multimodal contrastive learning objective based on InfoNCE^73^ with the sequence modality serving as the central alignment anchor. The total alignment loss is computed across all available modality pairs, enabling the model to learn a unified representation spanning sequence, structure, pocket geometry, functional annotations, and conformational dynamics. For complete details of the pretraining loss formulation, architectural hyperparameters, and optimization procedure, we refer the reader to the original OneProt model described in our recent study^48^. More details on encoder architecture is available in Supporting Information Text S1.

### Pocket Construction and Structural Processing

For the PPI-Site dataset, binding pockets were reconstructed from cavity annotations. The geometric centroid of annotated ligand atoms defined the pocket center. Residue centroids were computed as the average coordinates of all atoms in each residue. Residues were ranked by Euclidean distance to the ligand center, and the K — 100 nearest residues were retained to construct fixed-size pocket representations.

For the ASD dataset, residue-level annotations were converted to pockets using DBSCAN clustering.^74^ Given residue coordinates 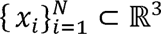, two residues are neighbors if 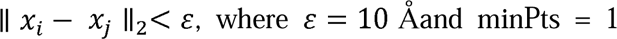. The largest cluster was retained and its centroid defined the pocket center, followed by selection of the 100 nearest residues.

To assess structural overlap between allosteric and orthosteric pockets in PPI-Site, we compared pockets sharing the same protein and chain identifiers at both residue-composition and atom-coordinate levels. Residue-level overlap was quantified using the Jaccard similarity coefficient and the overlap coefficient. Coordinate-level similarity was assessed using KD-tree nearest-neighbor search, measuring the fraction of atoms in one pocket within 4□Å of any atom in the other pocket, and the mean nearest-neighbor distance. We identified 100 pocket pairs (<5% of the dataset) with identical residue composition (Jaccard = 1.0) but substantial coordinate-level divergence (mean distances ranging from <0.1□Å to >18□Å), indicating that allosteric and orthosteric regulation can emerge from distinct conformations within shared local environments. More details on pocket construction and analysis is available in Supporting Text S2.

### Encoder Ablation Studies

To investigate how different biological modalities contribute to allosteric discrimination, we conducted systematic multimodal ablation experiments across all datasets. Rather than treating modality selection solely as an architectural optimization problem, these experiments were designed to probe which biological signals encoded within the pretrained latent space remain informative under different structural and statistical regimes. The ablation strategy operates at two distinct levels. At the encoder level, we control which modalities participate in multimodal pretraining and thus shape the organization of the shared latent space. At the embedding level, we control which modality-specific embeddings are extracted from the pretrained backbone and provided as input features to the downstream classifier. This two-level design distinguishes between two complementary effects: (i) how the inclusion of different modalities during pretraining shapes latent-space organization, and (ii) how different combinations of extracted embeddings contribute to downstream classification. The OneProt framework allows flexible inclusion or exclusion of modality-specific encoders during multimodal pretraining. All configurations retain the sequence encoder as the central anchor for multimodal alignment. The pocket encoder is required to obtain binding-site-specific representations for annotated pockets. The text encoder is retained because previous OneProt experiments demonstrated that textual functional annotations improve multimodal alignment and downstream transfer performance.

Against this common sequence–pocket–text backbone, we systematically ablate three structural and dynamical encoders (Table 1). The first encoder, Structural Tokens (ST), employs token-based structural representations derived from Foldseek structural tokens. This modality captures global fold-level organization and long-range structural context by encoding protein structures as discrete tokens analogous to sequence tokens. The second encoder, Structural Graph (SG), utilizes graph-based structural representations from ProNet. Unlike the global token-based approach, this modality captures fine-grained local geometry and contact relationships by representing proteins as graphs of residues with edges defined by spatial proximity. The MD encoder leverages MD trajectory representations from MDGen.

**Table 1.** Modalities and encoders assessed for their contribution to binding site prediction performance.

| Abbreviation | Modality | Input Type | Information Captured | Reference |
| --- | --- | --- | --- | --- |
| **ST** | Structural Token | Foldseek tokens | Global fold-level organization; long-range structural context | Foldseek<sup>49</sup> |
| **SG** | Structural Graph | Protein graph (nodes: residues; edges: spatial contacts) | Fine-grained local geometry; contact relationships | ProNet<sup>50</sup> |
| **MD** | Molecular Dynamics | MD trajectory frames | Conformational fluctuations; ensemble behavior; temporal dynamics | MDGen<sup>52</sup> |

We evaluate seven encoder configurations representing a progression from the minimal backbone to the full multimodal model. Pocket+Text is a baseline configuration. Only the sequence, pocket, and text encoders participate in pretraining. No structural or MD information is available. This establishes the performance floor against which all structural and dynamical contributions are measured. ST+Pocket+Text encoder combination adds token-based structural representations (ST) to the baseline. This configuration tests whether global structural context provides useful signal for allosteric discrimination beyond sequence, pocket geometry, and functional annotations. SG+Pocket+Text encoder combo adds graph-based structural representations (SG) to the baseline. This evaluates whether fine-grained local geometry and contact relationships provide complementary information to token-based representations. MD+ST+Pocket+Text encoder adds MD trajectories to the token-based structural configuration. This evaluates whether conformational dynamics provide additional signal beyond static structural context. MD+SG+Pocket+Text encoder adds MD trajectories to the graph-based structural configuration. This evaluates whether dynamics complement fine-grained static geometry. ST+SG+Pocket+Text encoder combines both structural encoders without MD. This evaluates whether token-based and graph-based representations encode overlapping or complementary information. MD+ST+SG+Pocket+Text encoder represents full multimodal configuration. All encoders participate in pretraining. This establishes the performance ceiling. For each encoder configuration, we use the corresponding pretrained OneProt model with frozen weights. The configuration determines which modalities contribute to latent-space organization during pretraining. Downstream classifiers then receive embeddings extracted from the available modalities.

### Embedding-Level Ablations: Feature Combinations for Downstream Classification

For each encoder configuration described above, we evaluate four different combinations of extracted embeddings as input features to the downstream MLP classifier (Table 2). These combinations are designed to probe the biological scale at which discriminative allosteric information is encoded—from purely local pocket geometry to broader evolutionary and functional context. We construct the following feature vectors where z_pocket, z_seq, z_text ∈ □¹□²□ denote the normalized embeddings extracted from the frozen OneProt backbone for a given protein sample. For each encoder configuration (determining which modalities are available), we train separate MLP classifiers for each embedding combination. This design enables us to isolate the contribution of each modality at the feature level while controlling for whether that modality participated in latent-space organization during pretraining.

**Table 2.** Feature combinations for downstream MLP classification.

| Combination | Features | Dimension | Modalities Included |
| --- | --- | --- | --- |
| **Pocket** | z_pocket | 1024 | Structural (pocket only) |
| **Pocket + Text** | z_pocket + z_text | 2048 | Structural + Text |
| **Pocket + Sequence** | z_pocket + z_seq | 2048 | Structural + Sequence |
| **Pocket + Sequence + Text** | z_pocket+z_seq+ z_text | 3072 | Structural + Sequence + Text |

The experimental design can be summarized as follows. For encoder-level (what modalities shaped the latent space) we employ seven configurations (Pocket+Text, ST+Pocket+Text, SG+Pocket+Text, MD+ST+Pocket+Text, MD+SG+Pocket+Text, ST+SG+Pocket+Text, MD+ST+SG+Pocket+Text). For embedding-level (what features are given to the classifier) we used four combinations (Pocket, Pocket+Text, Pocket+Sequence, Pocket+Sequence+Text). In total, the full experimental design involves 28 encoder–embedding combinations per dataset with combination evaluated across 10 random seeds, thus amounting to 1,120 independent runs.

### Downstream Allostery Classification

For downstream tasks, MLP classifiers were trained on embeddings extracted from the frozen OneProt backbone. The MLP consisted of an input layer matching the embedding dimension, hidden layers, and a single sigmoid output unit for binary classification. Hyperparameters, including the number and size of hidden layers, activation function, dropout rate, learning rate, batch size, layer normalization were optimized as described in the Supporting Information. For the selected architecture with two hidden layers, the classifier computes

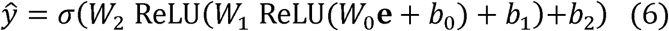

where edenotes the extracted embedding vector and *σ*(·)is the sigmoid activation. Training minimized the binary cross-entropy loss

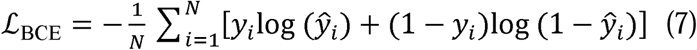

The downstream MLP classifier was optimized using the AdamW optimizer. Models were trained for up to 50 epochs with early stopping based on the validation loss (patience = 20 epochs). For each run, the checkpoint achieving the lowest validation loss was selected and subsequently evaluated on the validation and test sets. Each experimental configuration was evaluated across ten independent random seeds. For each run, the model achieving the highest validation ROC-AUC was selected, and the corresponding test metrics were aggregated across runs. Details of the downstream hyperparameter search are provided in Supporting Information. For AlloDiverse, positive samples (*N*_allo_ — 15,904) substantially outnumbered negative samples (*N*_ortho_ — 5,203). To mitigate training bias, positive instances were randomly subsampled during each training iteration to maintain balanced mini-batches (*B/2* positive and *B/2* negative per batch), while validation and test distributions remained unchanged.

### Variance Decomposition Analysis

To quantify the relative contributions of dataset composition, embedding selection, encoder architecture, and stochastic variation, we performed factorial ANOVA. Each run was represented by ROC-AUC with four categorical factors: Dataset (4 levels), Embedding (4 levels), Architecture (7 levels), and Random Seed (10 levels). A global ANOVA model was first fitted across all experimental runs to estimate the overall contributions of dataset, embedding combination, and encoder architecture together with their pairwise interactions:

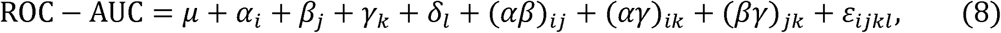

where *α_i_, β_J_, γ_k_, and δ_l_* denote dataset, embedding combination, encoder architecture, and random seed effects respectively, while *ε_ijkl_ ∼ N(0, σ^2^)* is the residual error. To characterize modality contributions within each biological regime, separate ANOVA models were subsequently fitted for each dataset. Since the dataset factor is fixed in these analyses, the model includes only embedding combination, encoder architecture, and their interaction:

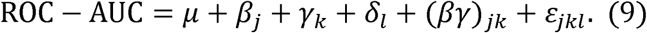

For both the global and dataset-specific analyses, sums of squares were normalized by the total sum of squares to obtain the percentage variance explained by each factor:

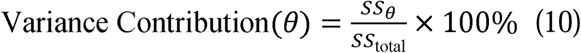

All statistical computations were performed in Python 3.10 using SciPy (v1.10) for hypothesis testing, statsmodels (v0.14)^75^ for multiple comparison correction. Visualization of distributions were generated with Matplotlib (v3.7)^76^ and GraphPad Prism version 11.0.

## Results

### A Multi-Stage Analytical Framework for Investigating Multimodality in Differentiating Orthosteric and Allosteric Binding Sites

To systematically investigate how multimodal protein representations capture allosteric regulation across diverse biological contexts, we designed a staged analytical framework that progresses from dataset characterization through modality ablation to interpretative synthesis (Figure 1). This framework integrates three complementary pillars: (i) a curated set of protein binding site datasets spanning distinct structural and statistical regimes, (ii) the OneProt multimodal foundation model with systematic encoder and embedding ablations, and (iii) a variance decomposition framework to quantify the relative contributions of dataset composition, modality selection, and architectural choice. Our analytical strategy proceeds through four interconnected stages.

We first characterize the intrinsic separability of allosteric and orthosteric binding environments across datasets with varying structural diversity, conformational heterogeneity, and class balance. We then systematically ablate embedding combinations—testing pocket-only, pocket+text, pocket+sequence, and full multimodal configurations—to determine which biological modalities contribute most strongly to allosteric discrimination under different dataset conditions. Next, we evaluate distinct encoder architectures, including structure-token (ST), structure-graph (SG), and MD encoders, to assess whether the specific implementation of structural and dynamical representations influences predictive performance. Finally, we perform variance decomposition to quantify the relative contributions of dataset identity, embedding composition, and architectural choice, establishing a rigorous statistical basis for our conclusions.

This staged design addresses three central questions. First, does the utility of multimodal integration depend on the intrinsic separability of the dataset? Second, which modalities provide complementary information for allosteric discrimination, and does this vary across biological contexts? Third, can we identify distinct learnability regimes that guide the application of multimodal protein foundation models to allostery prediction?

### Overall Performance Reveals a Hierarchy of Dataset Separability

Across all evaluated architectures, embedding combinations, and random seeds, a clear hierarchy of classification performance emerged (Figure 2). The PPI-Site dataset consistently represented the most challenging prediction regime, with ROC curves remaining close to the diagonal and median ROC-AUC values only modestly above random expectation (AUC ≈ 0.55–0.60) (Figure 2A). This near-random performance indicates that allosteric and orthosteric sites in PPI-Site (Figure 2A) share extensive structural and functional similarities that current representations cannot reliably resolve. The challenge arises from the dataset defining characteristic: class labels are assigned based on spatial relationships to protein–protein interaction interfaces rather than intrinsic physicochemical differences between binding sites. Consequently, the same protein, the same ligand-binding pocket, and often the same set of residues can be classified as either allosteric or orthosteric depending on subtle conformational arrangements relative to a protein–protein interface.

**Figure 2.**
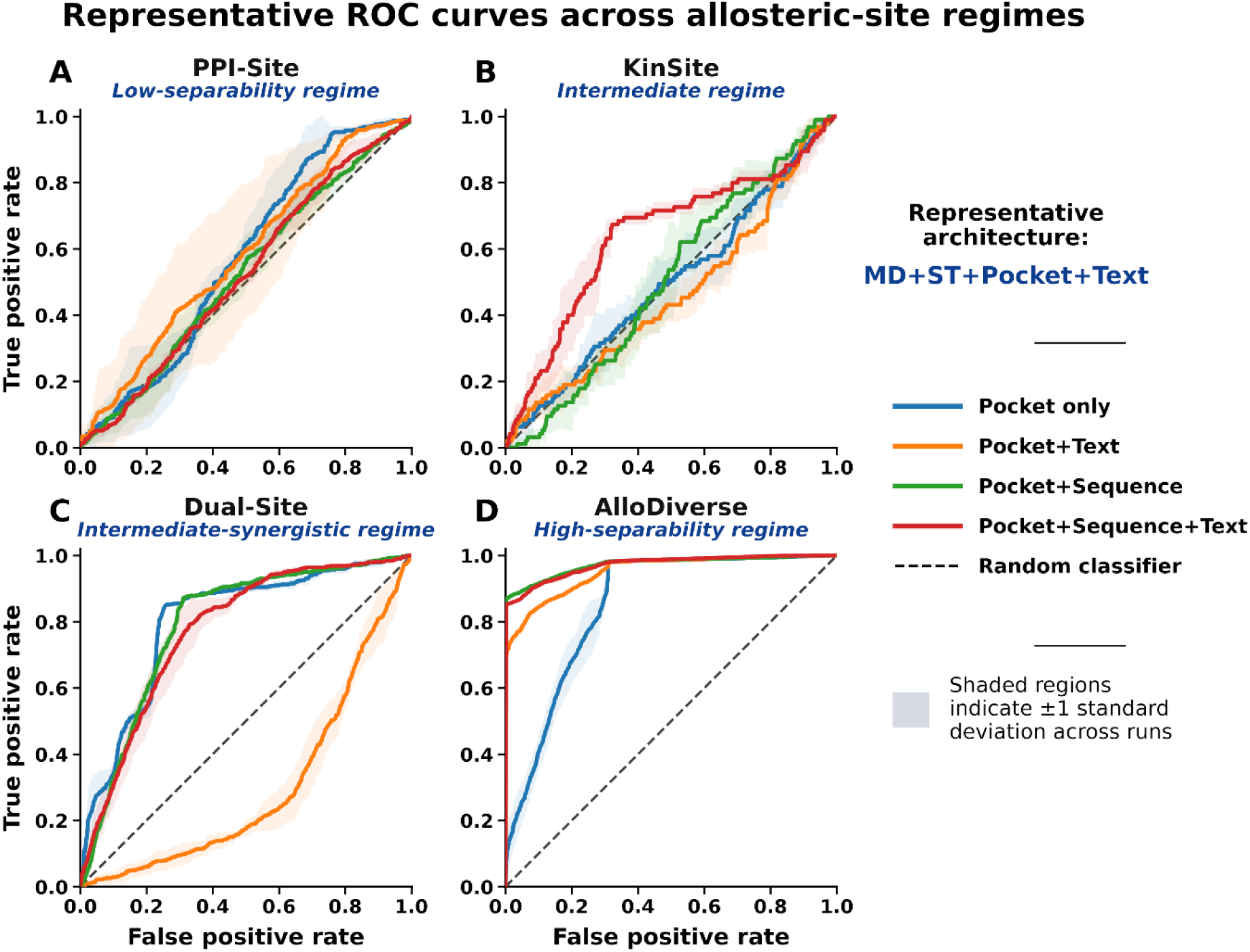
ROC Curves for OneProt Performance Across Datasets and Embedding Combinations. Receiver operating characteristic (ROC) curves illustrate predictive performance of a representative OneProt architecture including sequence, text, structure token, and molecular dynamics encoders across four datasets. Different colors correspond to the different embedding combinations used for predictions: Pocket only (blue), Pocket+Text (orange), Pocket+Sequence (green), and Pocket+Sequence+Text (red). The dashed diagonal line represents random performance (AUC = 0.5). Each curve represents the average across ten independent random seeds; shaded regions indicate ±1 standard deviation. PPI-Site shows the weakest discrimination (curves near diagonal), while AlloDiverse achieves near-ceiling performance across all embedding combinations. KinSite and Dual-Site occupy intermediate regimes with progressive improvement. The vertical dashed lines in each panel indicate the operating points corresponding to the best-performing embedding combination for each dataset, highlighting how the optimal sensitivity-specificity trade-off varies across regimes.

The KinSite (and Dual-Site datasets occupied intermediate positions between these extremes (Figure 2B,C). KinSite yielded AUC values ranging from approximately 0.70–0.75 (Figure 2B), reflecting the challenge of discriminating conserved ATP-binding sites from structurally heterogeneous allosteric pockets in a setting of extreme class imbalance (ρ ≈ 51.1).

The moderate performance improvement over PPI-Site arises from the strong evolutionary signals that distinguish orthosteric ATP-binding clefts—which are under intense purifying selection—from allosteric regulatory sites, which are evolutionarily permissive. Dual-Site exhibited progressively improved discrimination performance (AUC ≈ 0.80–0.85) as dataset diversity increased (Figure 2C), demonstrating that combining structurally overlapping (PPI-Site) and evolutionarily constrained (KinSite) datasets creates a synergistic effect that enables the model to learn more robust representations. In contrast, the AlloDiverse dataset yielded near-ceiling performance across nearly all evaluated configurations (AUC ≈ 0.95–0.98), indicating the presence of a strong and readily separable signal distinguishing allosteric and orthosteric sites (Figure 2D). This dramatic performance improvement reflects the broad structural and functional heterogeneity introduced by incorporating allosteric sites from the Allosteric Database (ASD), which spans diverse protein families. The diversity of allosteric architectures in AlloDiverse creates discriminative signals that multiple modalities can capture, making the regulatory classes readily separable even with relatively simple representations. Full details on the ROC-AUC curves are provided in Supporting Figure S1 and Supporting Table S1. The extreme class imbalance in KinSite makes precision-recall analysis particularly relevant for assessing model performance; we therefore evaluate PR-AUC in addition to ROC-AUC (Supporting Figure S2).

This progression became particularly apparent when performance distributions across all architectures and random seeds were examined collectively (Figure 3). The PPI-Site dataset formed a low-learnability regime characterized by broad overlap with random performance (distributions centered near a median ROC-AUC of 0.55) and substantial sensitivity to model configuration (Figure 3A). Critically, embedding composition had minimal effect within this regime: pocket-only, pocket+text, pocket+sequence, and full multimodal combinations all produced overlapping distributions, indicating that additional modalities could not compensate for the weak underlying discriminative signal. This observation underscores the fundamental limitation of the PPI-Site dataset: when allosteric and orthosteric sites share structural and evolutionary contexts, even sophisticated multimodal representations cannot reliably distinguish them.

**Figure 3.**
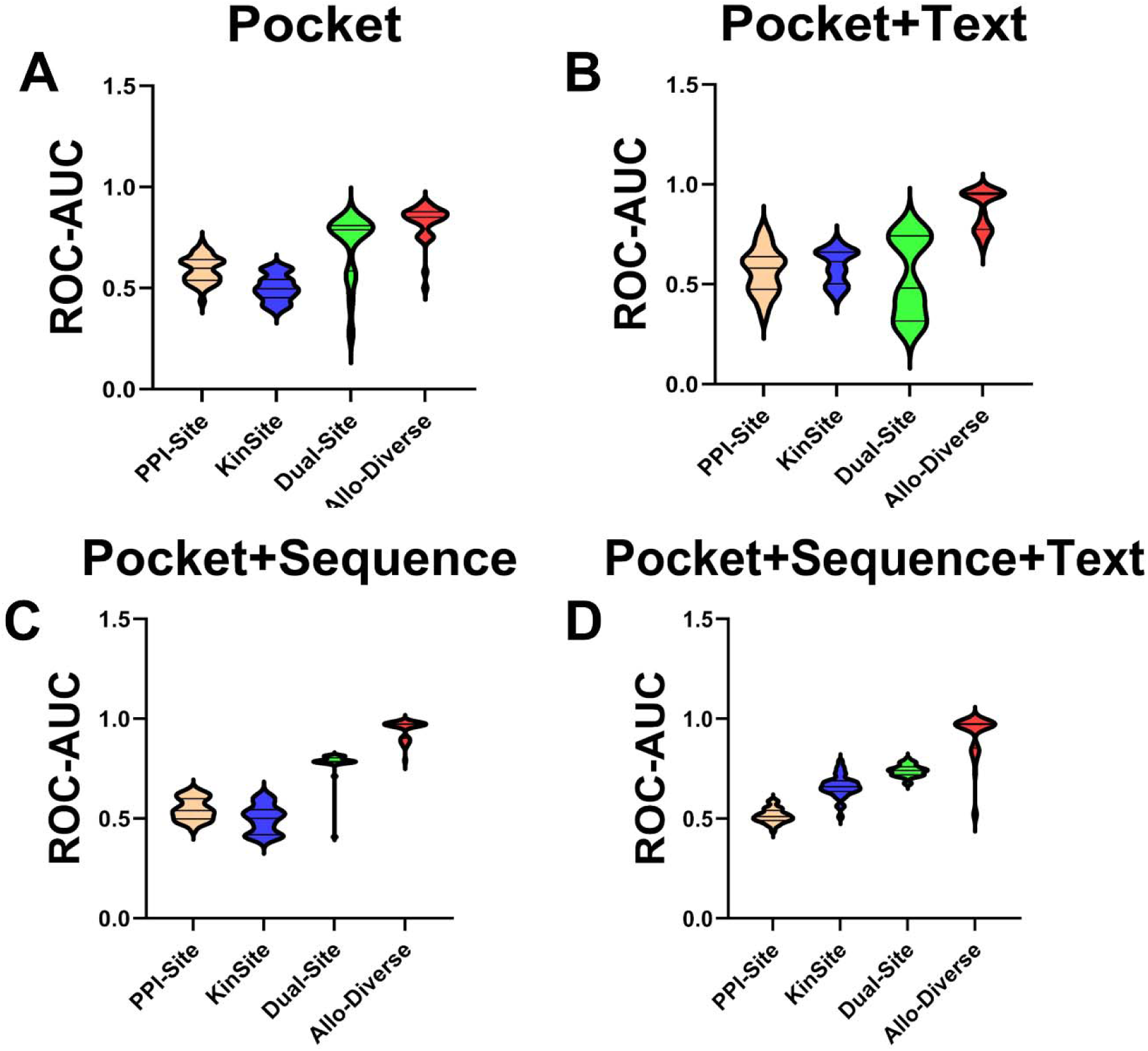
Distribution of ROC-AUC Values Across All Encoder Architectures and Random Seeds. Violin plots showing the distribution of ROC-AUC values across all encoder architectures and random seeds for each dataset and embedding combination. Different colors indicate embedding combinations: Pocket only (blue), Pocket+Text (orange), Pocket+Sequence (green), and Pocket+Sequence+Text (red). Each point represents a single run (one architecture, one random seed). Horizontal dashed line indicates random performance (AUC = 0.5). The figure reveals a systematic progression across datasets: PPI-Site (low-separability), KinSite (intermediate), Dual-Site (intermediate-synergistic), and AlloDiverse (high-separability). Embedding composition shows minimal effect in PPI-Site, whereas sequence-containing embeddings outperform pocket-only in KinSite, Dual-Site, and AlloDiverse. The narrowing of distributions from PPI-Site to AlloDiverse indicates that as the discriminative signal strengthens, performance becomes less sensitive to both architecture and embedding choices.

The KinSite dataset exhibited modest but reproducible improvements over PPI-Site, with AUC distributions centered near 0.70–0.75 and relatively compact interquartile ranges (Figure 3B). Unlike PPI-Site, embedding composition began to differentiate performance here: sequence-containing embeddings (pocket+sequence and pocket+sequence+text) consistently outperformed pocket-only and pocket+text combinations. This suggests that while overall performance is constrained by the extreme class imbalance (ρ ≈ 51.1), the evolutionary signals captured by sequence embeddings provide a consistent discriminative advantage. The compact interquartile ranges further indicate that this signal is robust across architectures, with structural and MD encoders providing modest but reproducible gains.

The combined Dual-Site dataset shifted toward substantially higher ROC-AUC values (AUC ≈ 0.80–0.85), with broader distributions reflecting the increased complexity and diversity of the merged dataset (Figure 3C). The widening of distributions compared to KinSite indicates that combining datasets with complementary biological characteristics—structural diversity from PPI-Site and evolutionary signals from KinSite—creates a more heterogeneous prediction landscape. Within this regime, embedding composition exerted its strongest effect: the gap between pocket-only and sequence-containing embeddings widened considerably, with pocket+sequence+text achieving the highest median performance. Architectural effects also became more apparent, as the addition of structural encoders consistently shifted distributions toward higher AUC values.

Finally, incorporation of ASD-derived allosteric sites produced a highly separable regime in which most evaluated models achieved consistently strong performance (AUC ≈ 0.95–0.98) with remarkably narrow distributions (Figure 3D). The near-ceiling performance across all embedding combinations and architectures indicates that the discriminative signal is so strong that it overwhelms architectural and embedding-specific variations. Notably, even pocket-only embeddings achieved AUC > 0.94, suggesting that the structural and functional diversity of allosteric sites in this dataset creates a signal that is readily separable regardless of representation choice. The narrow distributions further indicate that performance is robust across random seeds and architectural configurations, making this the most reliable prediction regime.

### Variance Decomposition Identifies Dataset Regimes as Primary Performance Determinants

The observed performance hierarchy remained stable despite extensive variation in both embedding composition and encoder architecture. To quantify the relative contributions of these factors, we performed variance decomposition across all evaluated datasets, embedding combinations, architectures, and random seeds (Figure 4). This analysis partitions the total variance in ROC-AUC into components attributable to dataset identity, embedding composition, encoder architecture, their interactions, and stochastic variation, providing a rigorous statistical basis for understanding what drives performance differences. Dataset identity accounted for the dominant fraction of explainable variance of 63.7% substantially exceeding the contributions of embedding composition (1.4%) and architecture choice (3.4%). Random seed effects were negligible (<0.5%). This overwhelming dominance of dataset identity indicates that the primary determinant of predictive performance is not architectural optimization or modality selection, but rather the intrinsic separability of allosteric and orthosteric binding environments within each dataset. In other words, the biological signal available for discrimination is largely fixed by the data itself, and model design choices can only marginally modulate what is already present. Notably, interaction terms between dataset and representation components explained additional variance, with dataset–embedding and dataset–architecture interactions contributing 8.3% and 5.8%, respectively (Figure 4). These interactions are particularly informative: they reveal that neither embedding choice nor architecture exerts a universal effect on performance. Instead, their influence depends strongly on the biological context represented by a particular dataset. For example, sequence embeddings that are essential for discriminating kinase binding sites may be irrelevant or even detrimental when applied to PPI-Site data, where allosteric and orthosteric pockets share evolutionary and functional contexts. This context dependence motivates the dataset-specific analysis that follows.

**Figure 4.**
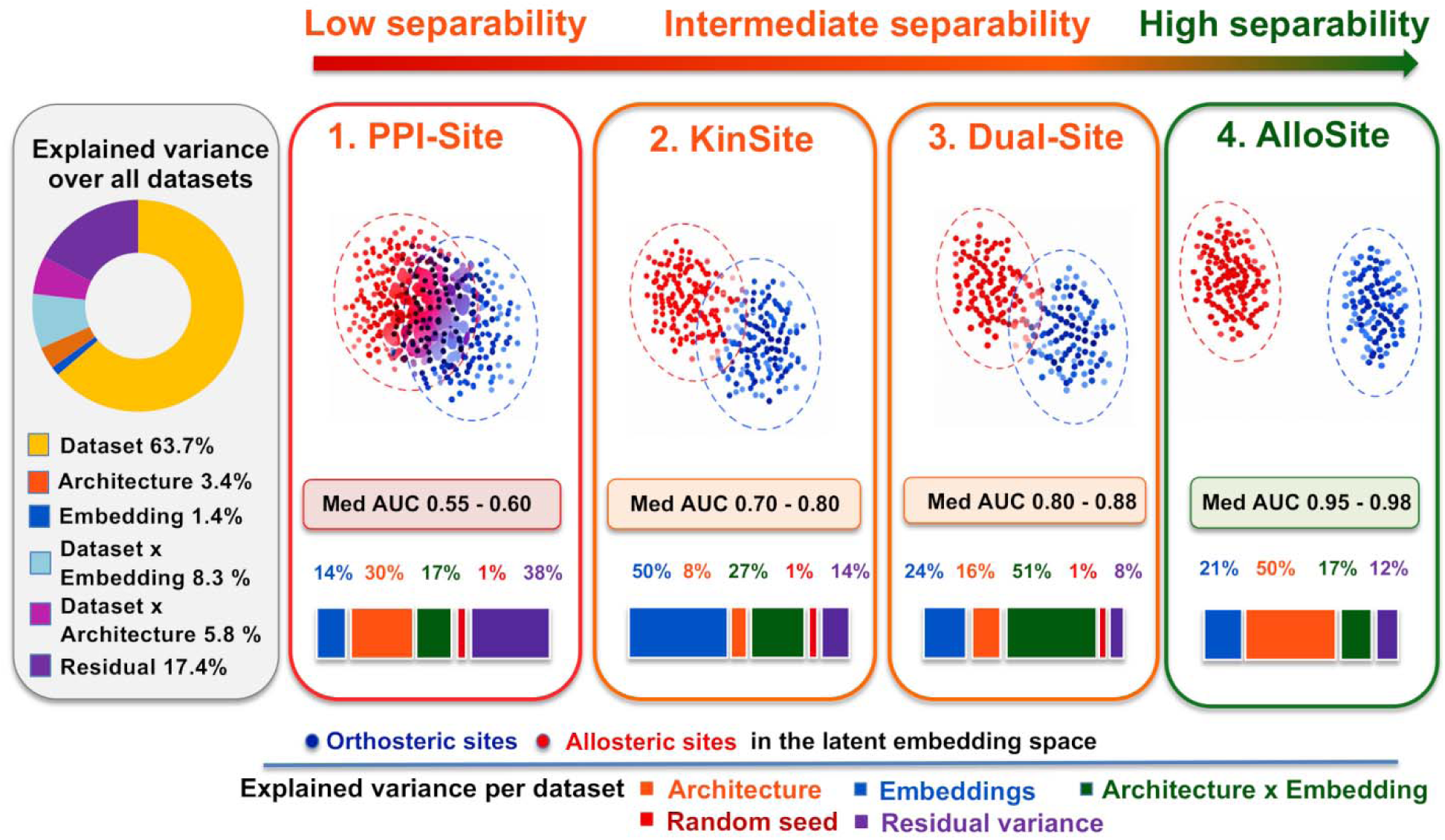
Variance Decomposition of ROC-AUC Across Datasets, Embedding Combinations, Architectures, and Random Seeds. (Main panel) Global variance decomposition across all experimental runs, showing the proportion of total variance attributable to dataset identity, embedding composition, encoder architecture, their pairwise interactions, random seed, and residual error. Dataset identity accounts for the dominant fraction (63.7%), followed by dataset–embedding interaction (8.3%) and dataset–architecture interaction (5.8%). (Panels 1–4) Dataset-specific variance decompositions for PPI-Site, KinSite, Dual-Site, and AlloDiverse, respectively, showing the proportion of variance attributable to embedding composition, encoder architecture, their interaction, random seed, and residual error within each dataset. Each panel is scaled to its own total variance, enabling direct comparison of the relative importance of each factor within each regime. Shaded regions represent the proportion of variance explained by each component.

To further examine these context-dependent effects, we performed variance decomposition separately for each dataset (Figure 4). These analyses reveal how the determinants of performance shift systematically across the separability spectrum, providing insight into the underlying biological signals available in each regime.

In the low-separability PPI-Site regime, architectural variation explained the largest fraction of performance variance (30%), although a substantial residual component (38%) remained unexplained. This pattern reflects the weak discriminative signal and high uncertainty observed across all configurations: when allosteric and orthosteric sites are nearly indistinguishable, architectural refinements become the primary source of performance differences, but they are fundamentally limited by the absence of a strong biological signal. The large residual variance further underscores that performance in this regime is dominated by irreducible uncertainty rather than systematic factors under our control. In KinSite, embedding composition dominated the variance (50%), indicating that modality selection is the primary determinant of performance in this intermediate regime. This finding aligns with the biological dichotomy of kinase binding sites: orthosteric ATP-binding clefts generate strong sequence conservation signatures that protein language models can capture, while allosteric regulatory sites produce weak or absent sequence signals. In this regime, the critical decision is selecting the right biological signal—sequence, text, or structural—rather than choosing a particular encoder to extract it. The dominance of embedding composition over architecture suggests that once the appropriate modality is selected, the specific implementation matters less. The Dual-Site dataset exhibited the strongest architecture × embedding interaction (50%), demonstrating that predictive performance depends not only on the chosen representations but also on how effectively a particular architecture exploits them.

This synergy reflects the increased complexity of the merged dataset, where combining structural diversity (from PPI-Site) and evolutionary signals (from KinSite) creates complementary information sources. The strong interaction indicates that the optimal representation strategy must be matched to the appropriate architectural implementation—a combination that is not obvious from either dataset alone. In the highly separable AlloDiverse dataset, architecture emerged as the dominant contributor (50%), while residual variance remained low. This shift reflects a different regime: once a strong discriminative signal is present—provided by the broad structural and functional heterogeneity of allosteric sites spanning diverse protein families—architectural refinements become the primary source of performance differences. The convergence of multiple structurally informed architectures to near-ceiling performance in this regime further supports the interpretation that the inclusion of structural information rather than the specific encoder implementation is the critical factor once sufficient biological diversity is present.

Collectively, these dataset-specific variance decompositions reveal a systematic progression in the determinants of performance across the separability spectrum. (Table 3). The dominant contribution of dataset identity suggests that the primary challenge in allosteric site prediction is not necessarily architectural optimization but rather the intrinsic separability of allosteric and orthosteric binding environments within a given dataset. In the PPI-Site dataset, all evaluated models remained clustered near random performance regardless of representation choice, suggesting substantial overlap between allosteric and orthosteric competitive sites within the learned latent space. By contrast, the AlloDiverse dataset contained a strong discriminative signal that was readily captured by a broad range of representations. The KinSite and Dual-Site datasets occupied an intermediate regime in which performance differences between architectures and embedding combinations became more apparent (Table 3).

**Table 3.** Summary of separability regimes and key determinants of performance.

| Dataset | Regime | Dominant Modality | Key Architectural Finding |
| --- | --- | --- | --- |
| PPI-Site | Low | Pocket geometry | All architectures perform near-random (AUC ∼0.55–0.58); weak signal dominates. |
| KinSite | Intermediate | Pocket + Text | Structural encoders provide modest gains; modality selection drives performance. |
| Dual-Site | Intermediate-synergistic | Sequence | Architecture–embedding interaction is the primary determinant of performance. |
| AlloDiverse | High | Sequence +Text | Multiple structural architectures achieve near-ceiling performance (∼0.96–0.98 AUC). |

### Contribution of Multimodal Embeddings Across Separability Regimes

Having established that dataset composition is the dominant determinant of performance and that modality selection drives performance in intermediate regimes, we next examined how different embedding combinations contribute within each separability regime.

The average ROC-AUC values across datasets and embedding combinations revealed that no universally optimal representation strategy exists (Figure 5). Instead, the utility of individual modality embeddings depended strongly on dataset composition, reflecting the distinct biological signals available in each regime. In the low-separability PPI-Site regime, pocket-only embeddings achieved the strongest average performance (AUC = 0.59), while the addition of text and sequence representations progressively reduced predictive performance: Pocket+Text (AUC = 0.57), Pocket+Sequence (AUC = 0.55), and Pocket+Sequence+Text (AUC = 0.51) (Figure 5). This striking result—where adding more information consistently degrades performance, with the full multimodal combination performing worst—reflects the biological characteristics of the PPI-Site dataset. As established in our dataset characterization, allosteric and orthosteric sites in PPI-Site share the same protein family context, the same structural architecture, and often the same residue composition, differing only in conformational arrangement relative to protein–protein interaction interfaces. Under these conditions, global evolutionary and functional context—which is largely shared between the two pocket classes within a given protein—introduces variance rather than complementary information. The pocket-overlap analysis supports this interpretation: we identified 100 pocket pairs (<5% of the dataset) with identical residue composition (Jaccard = 1.0) but substantial coordinate-level divergence (mean distances ranging from <0.1 Å to >18 Å), indicating that regulatory discrimination depends primarily on subtle local structural differences. This finding aligns with the variance decomposition analysis, where residual variance dominated PPI-Site (38%), reflecting the weak discriminative signal and irreducible uncertainty that limit performance regardless of embedding strategy.

**Figure 5.**
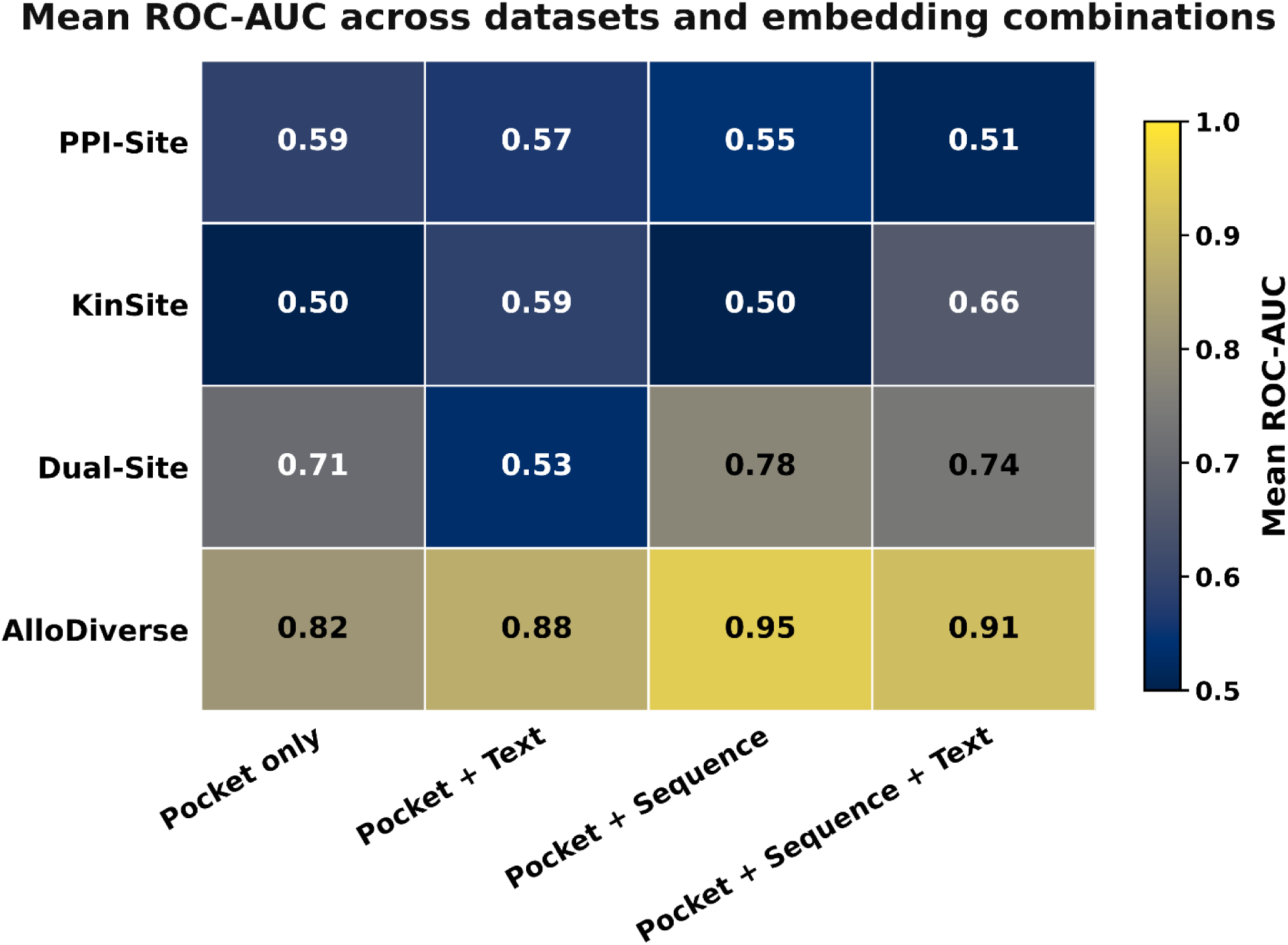
Mean ROC-AUC Heatmap Across Datasets and Embedding Combinations. Rows correspond to the four datasets—PPI-Site (0.59, 0.57, 0.55, 0.51), KinSite (0.50, 0.59, 0.50, 0.66), Dual-Site (0.71, 0.53, 0.78, 0.74), and AlloDiverse (0.82, 0.88, 0.95, 0.91)—ordered from low to high separability. Columns correspond to the four embedding combinations: Pocket only, Pocket+Text, Pocket+Sequence, and Pocket+Sequence+Text. Color intensity ranges from blue (low AUC) to red (high AUC), with numerical values displayed in each cell. The relative ordering of embedding combinations changes systematically across datasets: pocket-only performs best in PPI-Site, while text and sequence embeddings excel in other regimes. This regime-dependent progression reflects the distinct biological signals available in each dataset.

A fundamentally different pattern emerged in the KinSite dataset. Here, embedding combinations produced a clear hierarchy: Pocket-only (AUC = 0.50) < Pocket+Sequence (AUC = 0.50) < Pocket+Text (AUC = 0.59) < Pocket+Sequence+Text (AUC = 0.66) (Figure 5). The striking improvement from pocket-only to full multimodal (+0.16 AUC) indicates that evolutionary and functional context dramatically improves discrimination in this regime. Critically, text embeddings alone (Pocket+Text, AUC = 0.59) provided substantially more discriminative power than sequence embeddings alone (Pocket+Sequence, AUC = 0.50), and the combination of both (AUC = 0.66) yielded the strongest performance. This pattern reflects the biological characteristics of the KinSite dataset: while kinase ATP-binding clefts generate strong sequence conservation signatures, the most informative signal for distinguishing allosteric from orthosteric sites comes from functional annotations that capture protein-specific regulatory mechanisms—mechanistically diverse processes such as myristoyl pocket binding in ABL, αC-helix displacement in BRAF, and activation loop phosphorylation across many kinases. This finding is consistent with the variance decomposition analysis, where embedding composition dominates performance in this intermediate regime (50% of explained variance).

In the Dual-Site dataset, the embedding hierarchy shifted compared to KinSite: Pocket+Text (AUC = 0.53) < Pocket-only (AUC = 0.71) < Pocket+Sequence+Text (AUC = 0.74) < Pocket+Sequence (AUC = 0.78) (Figure 5). The merged dataset exhibited a more complex pattern: Pocket+Sequence emerged as the strongest-performing combination (AUC = 0.78), outperforming the full multimodal configuration (0.74). Notably, Pocket+Text performed poorly (AUC = 0.53), substantially worse than pocket-only (0.71), suggesting that functional annotations alone introduce noise when applied to the structurally diverse PPI-Site component of the merged dataset. The strong performance of Pocket+Sequence reflects the complementary nature of structural diversity (from PPI-Site) and evolutionary signals (from KinSite): sequence embeddings capture the differential evolutionary constraints that distinguish orthosteric from allosteric sites across both regimes. This pattern aligns with the variance decomposition finding that the architecture–embedding interaction is the primary determinant of performance in Dual-Site (50%), reflecting the increased complexity of the merged dataset where different modalities contribute differently depending on the biological context.

In the AlloDiverse dataset, the hierarchy was: Pocket-only (AUC = 0.82) < Pocket+Text (AUC = 0.88) < Pocket+Sequence+Text (AUC = 0.91) < Pocket+Sequence (AUC = 0.95) (Figure 5). The clear progression from pocket-only to sequence-containing embeddings indicates that evolutionary information provides the most substantial boost in this regime, consistent with the strong sequence conservation signatures of orthosteric sites across diverse protein families. Notably, Pocket+Sequence (AUC = 0.95) substantially outperformed the full multimodal combination (AUC = 0.91), suggesting that text annotations may introduce noise when combined with sequence information in this regime. The relatively high performance of pocket-only (AUC = 0.82) compared to lower regimes reflects the broad structural diversity of allosteric sites in AlloDiverse, which creates a discriminative signal that even simple geometric representations can partially capture. The persistence of gains from sequence information indicates that multimodal integration continues to improve discrimination even when a strong allosteric signal is already present. This observation is consistent with the variance decomposition result showing that architecture becomes the dominant factor (50%) in this high-separability regime, while residual variance remains low—indicating that once sufficient biological diversity is present, architectural refinements rather than modality selection become the primary source of performance differences.

Collectively, these results demonstrate that the contribution of multimodal information is highly context dependent and follows a systematic progression across the separability spectrum. In low-separability settings where allosteric and orthosteric sites share structural and evolutionary contexts, additional modalities introduce noise rather than complementary information. In intermediate regimes where differential evolutionary signals distinguish regulatory classes, sequence and text embeddings provide essential discriminative power. In high-separability settings where broad structural and functional diversity creates strong discriminative signals, multimodal integration provides marginal but consistent improvements. This context-dependent behavior, summarized in Table 3, has direct implications for model development: the optimal embedding strategy must be matched to the biological characteristics of the target dataset.

The complete ROC-AUC matrix underlying these findings is provided in Supporting Table S1, which confirms the systematic progression from low to high separability across all architectures and embedding combinations. A progression from low performance in PPI-Site to near-ceiling performance in AlloDiverse, confirms that dataset identity can be the dominant determinant of allosteric discrimination performance. The corresponding PR-AUC values in Supporting Table S2 further quantify the impact of class imbalance, showing that while ROC-AUC in KinSite reaches moderate levels (∼0.70–0.75), PR-AUC remains substantially lower (0.05–0.16), reflecting the extreme difficulty of recovering rare allosteric sites under severe class imbalance. In contrast, the near-ceiling PR-AUC values for AlloDiverse confirm robust positive-class recovery once sufficient allosteric diversity is present. The data demonstrate that balanced sampling during training has minimal impact on AlloDiverse performance, indicating that the discriminative signal in this regime is robust to sampling strategies (Supporting Table S2).

A comprehensive summary of mean ROC-AUC values across all architectures and embedding combinations is presented in Supporting Figure S3, confirming that dataset identity remains the dominant determinant of performance regardless of representation choice. Threshold-dependent sensitivity and specificity across datasets and embedding combinations are shown in Supporting Figures S4 and S5, respectively. Sensitivity for allosteric site recovery (Figure S4) and specificity for orthosteric site rejection (Figure S5) reveal that in PPI-Site, both metrics remain near random performance; in KinSite, specificity substantially exceeds sensitivity, reflecting the greater difficulty of recovering rare allosteric sites; in Dual-Site, both metrics improve with sequence-containing embeddings; and in AlloDiverse, sensitivity approaches ceiling performance while specificity remains more variable, indicating that models readily identify allosteric sites but differ in their ability to reject orthosteric sites. These observations reinforce the conclusion that dataset-dependent separability governs threshold-dependent classification behavior, consistent with the learnability regimes identified throughout the study.

### Structural and Dynamical Encoder Architectures Contribute Differently Across Separability Regimes

While embedding composition influenced performance in a dataset-dependent manner, substantial variability also remained between encoder architectures. We therefore examined whether different structural and dynamical encoders capture complementary aspects of allosteric regulation and how these architectural choices affect classification performance. To isolate architectural effects, we compared OneProt variants incorporating structure-token (ST), structure-graph (SG), and MD encoders while evaluating a fixed downstream embedding combination consisting of pocket, sequence, and text representations. The impact of encoder architecture depends strongly on the separability regime (Figure 6). In the PPI-Site dataset, all architectures produced ROC curves close to the diagonal and ROC-AUC values near random expectation (AUC ≈ 0.55–0.58). Although minor differences between encoder configurations were observed, none consistently outperformed the others. This result confirms that when allosteric and orthosteric sites share structural contexts and are distinguished primarily by conformational differences, architectural modifications alone are insufficient to recover a robust discriminative signal.

**Figure 6.**
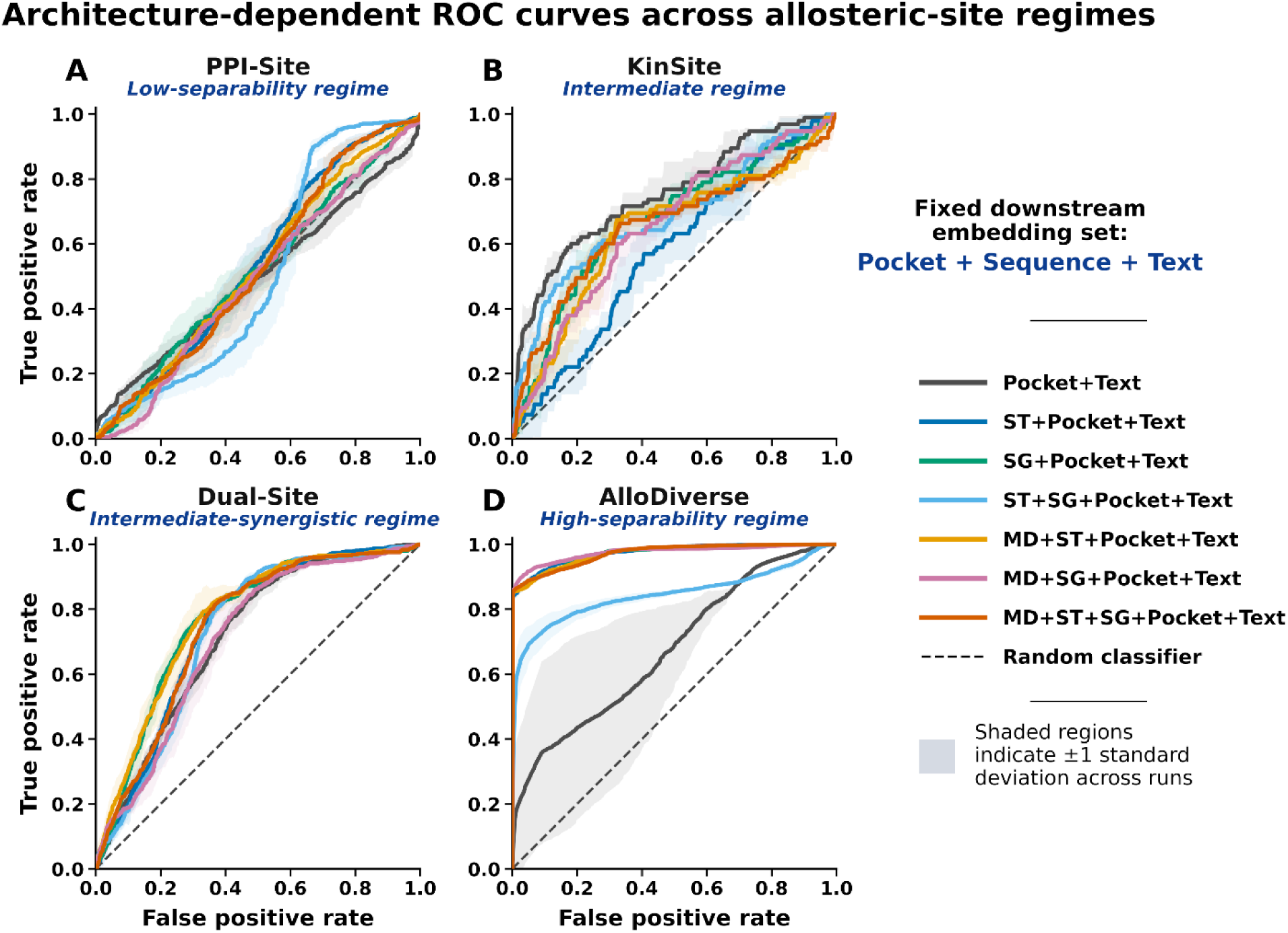
ROC Curves for Different Architectural Encoder Combinations with Fixed Embedding Combination. Different colors indicate encoder configurations: Pocket+Text (baseline, no structural encoders; gray), ST+Pocket+Text (structure-token; blue), SG+Pocket+Text (structure-graph; green), MD+ST+Pocket+Text (MD + structure-token; orange), MD+SG+Pocket+Text (MD + structure-graph; purple), ST+SG+Pocket+Text (both structural encoders; teal), and MD+ST+SG+Pocket+Text (full multimodal; red). The dashed diagonal line represents random performance (AUC = 0.5). Architectural effects vary dramatically across datasets: in PPI-Site, all architectures cluster near random performance; in KinSite, structural encoders provide modest improvements; in Dual-Site, structural encoders substantially outperform the baseline; and in AlloDiverse, multiple structural architectures converge to near-ceiling performance. Full details on the ROC-AUC curves are provided in Supporting Figure S1 and Supporting Table S1.

More substantial architectural differences emerged in the KinSite and Dual-Site datasets. In KinSite, the Pocket+Text baseline remained among the strongest-performing architectures, suggesting that local pocket geometry combined with functional annotations captures a substantial fraction of the discriminative signal. The structure-token-only (ST+Pocket+Text) architecture achieved the highest median AUC (≈0.74) and exhibited the most compact performance distribution, indicating that global fold-level organization captured by Foldseek tokens provides complementary information beyond local pocket features. In contrast, the Pocket+Text architecture became the weakest-performing configuration in the Dual-Site dataset, where the inclusion of structural encoders consistently improved discrimination. This reversal reflects the increased structural and functional diversity introduced by combining kinase and non-kinase proteins. Within the relatively homogeneous kinase family, text-derived functional information provides a strong contextual signal. However, in the more heterogeneous Dual-Site regime, explicit structural representations become essential for capturing differences that generalize across distinct protein folds and regulatory mechanisms. Notably, several structurally informed architectures converged to similar ROC-AUC values (0.84–0.87), indicating that the common factor underlying improved performance was the incorporation of structural information itself, rather than the specific choice of structural encoder.

A similar pattern emerged in the AlloDiverse dataset. Architectures incorporating any structural modality consistently achieved near-ceiling ROC-AUC values (AUC ≈ 0.96–0.98), whereas the Pocket+Text baseline remained substantially lower with considerably greater variability (AUC ≈ 0.93). Structure-token, structure-graph, and MD architectures converged to similar performance levels despite their distinct representational strategies, reinforcing that the critical factor is the presence of structural information rather than the specific encoder. Notably, the MD encoder—which captures conformational fluctuations and ensemble behavior—did not provide additional gains beyond static structural representations in this regime, suggesting that the discriminative signal in AlloDiverse is sufficiently strong that static features already capture the relevant information. The full set of ROC curves across all seven encoder architectures and embedding combinations is provided in Supporting Figure S1.

The architecture-specific performance distributions revealed several notable trends (Figure 7). The progression from PPI-Site to AlloDiverse reveals a systematic shift in the role of architecture. In PPI-Site all architectures cluster near random performance (AUC ≈ 0.55–0.58) regardless of embedding combination (Figure 7A), confirming that when allosteric and orthosteric sites share structural contexts, architectural modifications alone are insufficient to recover a discriminative signal. In PPI-Site, only architectures incorporating structure-token representations (ST+Pocket+Text) or the full multimodal encoder consistently achieved performance above random expectation. This suggests that even in low-separability settings, fold-level structural context provides some weak discriminative signal that local pocket geometry alone cannot capture, although overall performance remains near-random. In KinSite (Figure 7B) the structure-token-only (ST+Pocket+Text) architecture exhibits the most compact distribution and highest median AUC (≈0.74), while architectures incorporating both ST and SG encoders show broader distributions and lower median performance, suggesting redundancy between structural representations. In Dual-Site architectures incorporating structural information (ST, SG, or MD) consistently outperform the Pocket+Text baseline, with convergence across multiple configurations (AUC ≈ 0.84–0.87), indicating that the presence of structural information—rather than the specific encoder—drives improved discrimination (Figure 7C). The ST+SG configuration did not exhibit corresponding improvement beyond individual structural encoders, confirming that these representations encode partially overlapping information. In contrast, architectures incorporating MD information formed part of the highest-performing group, indicating that dynamical information provides genuinely complementary signal in datasets where conformational heterogeneity distinguishes regulatory classes. In AlloDiverse, nearly all architectures containing structural information converge to near-ceiling performance (AUC ≈ 0.96–0.98), with the structure-token architecture remaining among the most stable configurations (Figure 7D). The narrowing of distributions from PPI-Site to AlloDiverse indicates that as the discriminative signal strengthens, performance becomes less sensitive to both architecture and embedding choices, consistent with the variance decomposition analysis (Figure 4) showing that encoder architecture explains substantially less performance variation than dataset composition. The structure-token architecture remained among the most stable configurations, while the full multimodal model achieved comparable performance without providing a substantial advantage.

**Figure 7.**
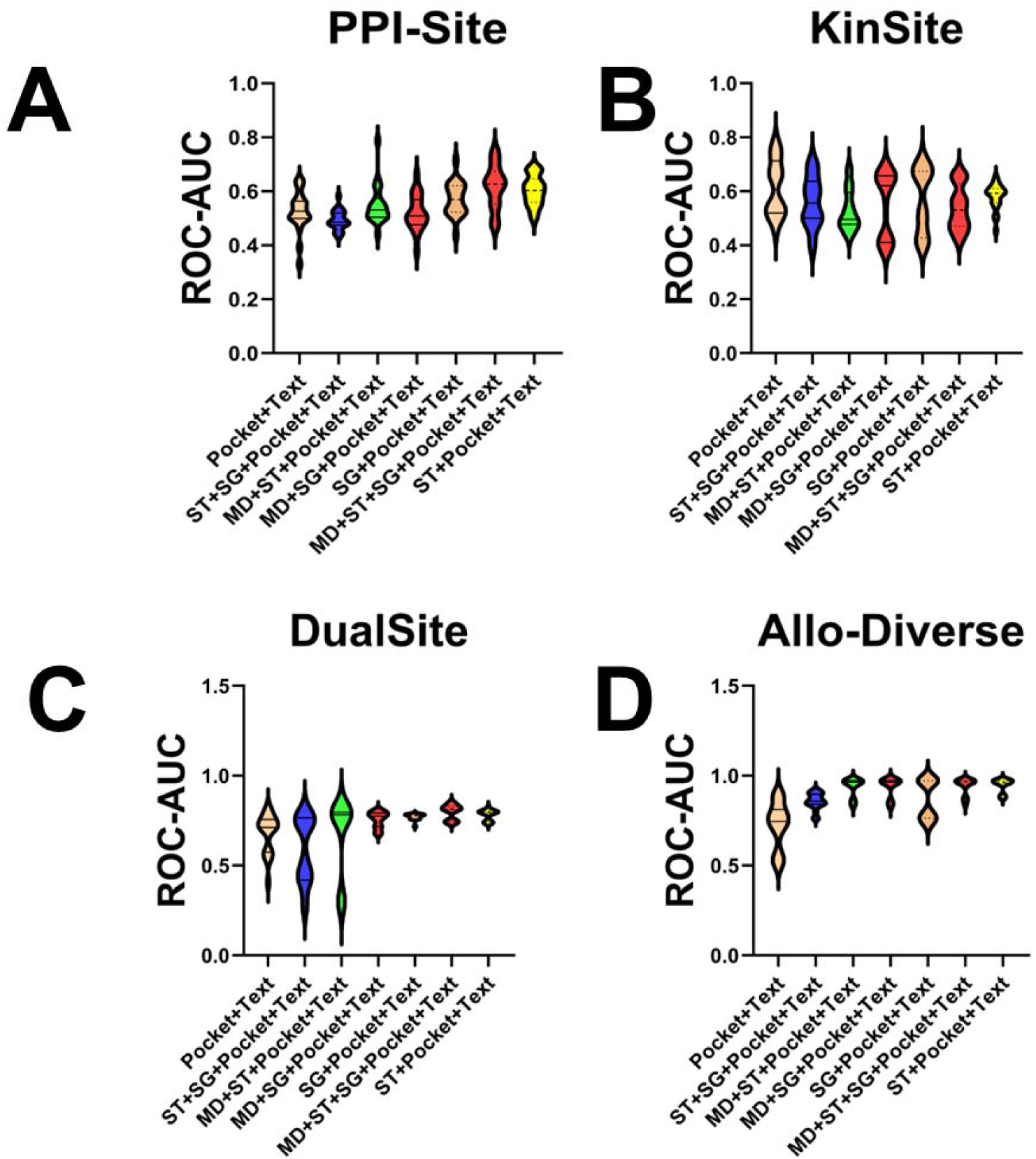
Architecture-Dependent ROC-AUC Distributions Across Separability Regimes. Violin plots showing the distribution of ROC-AUC values across seven OneProt encoder architectures and ten independent random seeds for each of the four datasets. Each panel corresponds to a distinct separability regime: (A) PPI-Site (low-separability), (B) KinSite (intermediate), (C) Dual-Site (intermediate-synergistic), and (D) AlloDiverse (high-separability). The x-axis within each panel denotes the seven encoder architectures evaluated: Pocket+Text (baseline, no structural encoders), ST+Pocket+Text (structure-token), SG+Pocket+Text (structure-graph), MD+ST+Pocket+Text (MD + structure-token), MD+SG+Pocket+Text (MD + structure-graph), ST+SG+Pocket+Text (both structural encoders), and MD+ST+SG+Pocket+Text (full multimodal). Colored points represent the four downstream embedding combinations used for classification: Pocket only (blue), Pocket+Text (orange), Pocket+Sequence (green), and Pocket+Sequence+Text (red). Each point corresponds to a single independent run, with horizontal bars indicating the median and interquartile range. The dashed horizontal line denotes random performance (ROC-AUC = 0.5).

Because several datasets exhibited substantial class imbalance, particularly the KinSite dataset (ρ ≈ 51.1), we additionally evaluated model behavior using precision-recall (PR) curves (Supporting Figure S2). The PR curves reveal that while ROC-AUC values in KinSite are moderate (≈0.70–0.75), the corresponding PR-AUC values are substantially lower, reflecting the difficulty of recovering rare allosteric sites under extreme class imbalance. In contrast, the highly separable AlloDiverse regime achieves near-perfect PR-AUC across most architectures and robust positive-class recovery. The complete PR-AUC values across all configurations are provided in Supporting Table S2, which further quantifies the performance differences between architectures and embedding combinations under imbalanced conditions. The data demonstrate that while ROC-AUC values in KinSite are moderate (≈0.70–0.75), the corresponding PR-AUC values are substantially lower (≈0.05–0.16), reflecting the extreme difficulty of recovering rare allosteric sites under severe class imbalance (Supporting Table S2).

Taken together, these results demonstrate that architectural effects are highly context-dependent. In low-separability regimes, performance is constrained by dataset characteristics and remains largely insensitive to encoder design. In intermediate regimes, structural and dynamical encoders contribute meaningfully, with different encoders providing complementary benefits depending on the biological characteristics of the dataset. In high-separability regimes, multiple architectural solutions achieve near-optimal discrimination, and the specific encoder choice becomes less critical. Consistent with the variance decomposition analysis (Figure 4), encoder architecture explains substantially less performance variation than dataset composition. Nevertheless, architectural effects become increasingly apparent once the underlying biological signal is sufficiently informative, particularly in intermediate regimes where the choice of structural representation can meaningfully impact discrimination.

### Decoding the Structural and Functional Signatures of Allosteric and Orthosteric Binding Sites Across Separability Regimes

To bridge the gap between multimodal prediction performance and the underlying biological organization of binding sites, we performed a systematic structural and functional analysis of true positive (TP) and true negative (TN) predictions recovered by the best-performing model configuration (Pocket+Sequence+Text embeddings with the full multimodal encoder architecture). This analysis connects the observed separability regimes to the physical and evolutionary characteristics that distinguish allosteric from orthosteric regulatory sites across different protein families and structural contexts.

The central premise of this analysis is that prediction success—or failure—reflects the degree to which the available modalities capture biologically meaningful discriminative signals. When allosteric and orthosteric sites share structural and evolutionary contexts, sequence and text modalities provide little additional information beyond what pocket geometry already captures. Conversely, when the two classes are distinguished by fundamentally different evolutionary constraints or structural architectures, complementary modalities become essential. By examining the structural families and functional annotations of correctly classified instances, we identify the biological signatures that drive separability in each regime.

The PPI-Site dataset presents the most challenging prediction regime, with all evaluated models struggling to achieve reliable discrimination (AUC ≈ 0.55–0.58 across all configurations). Analysis of the TP and TN predictions reveals that both classes are dominated by proteins sharing a common structural and functional theme: methyltransferases and SAM-dependent enzymes that utilize S-adenosylmethionine (SAM) or S-adenosylhomocysteine (SAH) as cofactors. These proteins, representing approximately one-third of all correctly classified instances across both classes, share a conserved Rossmann fold architecture—a ubiquitous nucleotide-binding motif consisting of a central β-sheet flanked by α-helices—in which the SAM-binding pocket is spatially adjacent to protein–protein interaction interfaces. Figure 8 illustrates the structural context of this conformational ambiguity through representative structures from the PPI-Site dataset. The analysis reveals that allosteric (PLA) and orthosteric (PLOC) sites in methyltransferases often involve the same protein, the same ligand-binding pocket, and frequently the same set of residues, differing only in the conformational relationship between the pocket and the protein–protein interaction interface. The prevalence of SAM-dependent methyltransferases among correctly classified instances and the near-random performance of all models on the broader PPI-Site dataset underscores a fundamental principle: when regulatory classes share structural and evolutionary contexts, the discriminative signal is primarily conformational rather than sequence-based or functional. Pocket geometry, which captures local structural arrangements, is the only modality that can access this signal, while sequence and text modalities provide no additional discrimination. This explains the key observation from our embedding ablation study: pocket-only embeddings achieved the strongest performance in PPI-Site (AUC ≈ 0.58), while the addition of sequence and text representations provided little benefit and in some cases reduced predictive performance (AUC ≈ 0.55–0.56).

**Figure 8.**
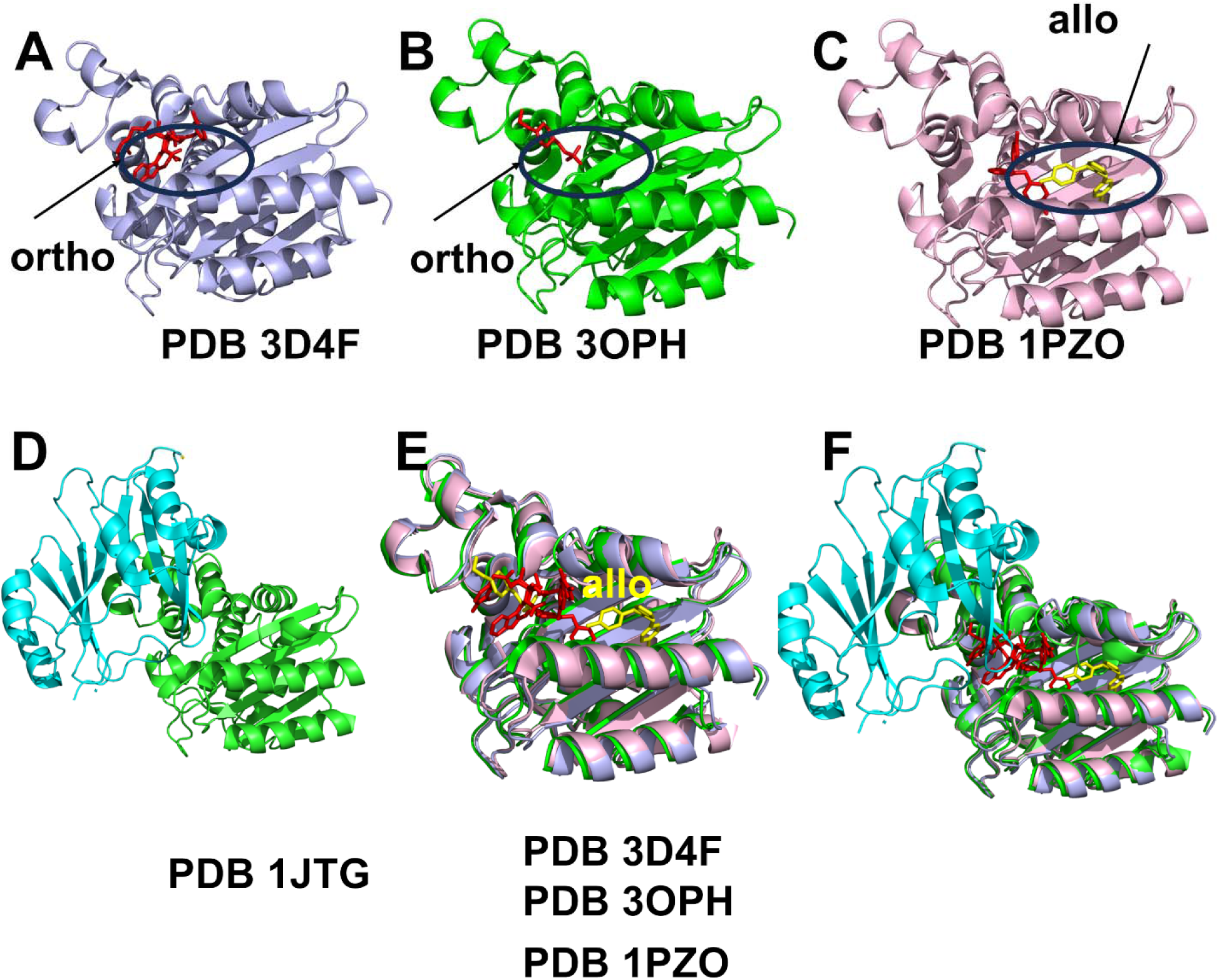
Structural context of allosteric and orthosteric binding sites in PPI-Site methyltransferases. The figure illustrates the conformational ambiguity that defines the low-separability regime of the PPI-Site dataset, where allosteric and orthosteric sites share structural and evolutionary contexts. (A) Rossmann fold architecture of a SAM-dependent methyltransferase (PDB: 3d4f) showing a correctly classified allosteric (PLA) site with the SAM-binding pocket (green) spatially distinct from the protein–protein interaction interface (blue). (B) Orthosteric SAM pocket (PDB: 3oph) showing direct overlap between the SAM-binding pocket (orange) and the protein–protein interaction interface (red). (C) Superposition of allosteric (PDB: 3d4f) and orthosteric (PDB: 1pzo) conformations revealing identical residue composition but distinct spatial arrangements relative to the interface. (D) Orthosteric site from PDB: 1jtg illustrating pocket integration with Rossmann fold architecture. (E) Truly allosteric ligand-binding site from PDB: 1pzo showing spatial separation between pocket and interface. (F) Superposition highlighting subtle conformational shifts between binding modes. This structural overlap illustrates why pocket geometry capturing local structural arrangements is the only modality providing discriminative signal in PPI-Site, while sequence and text provide no additional information.

The structural analysis further reveals that the allosteric classification in PPI-Site arises not from distinct regulatory mechanisms but from the spatial relationship between the ligand-binding pocket and a protein–protein interaction interface. In these methyltransferases, the SAM-binding pocket is proximal to the substrate-binding or protein interaction interface, and its allosteric annotation reflects the fact that ligand binding may influence protein–protein interactions through conformational changes rather than direct competition at an orthosteric site. This creates a regime where allosteric and orthosteric pockets are structurally coupled as they share the same local protein environment and often involve overlapping residue sets. This structural and functional coupling has profound implications for multimodal prediction, explaining the observed performance patterns in the PPI-Site regime. When allosteric and orthosteric sites share identical residue compositions and differ primarily in subtle conformational arrangements, global evolutionary signals and functional annotations introduce variance rather than complementary information.

### Structural and Functional Signatures in the KinSite Regime

The KinSite dataset presents a fundamentally different biological regime characterized by a stark dichotomy in binding site architecture and evolutionary constraint. Analysis of TP and TN predictions from the best-performing model configuration reveals that the standalone KinSite dataset yields only a relatively small number of correctly classified allosteric binding sites that are largely represented by type III allosteric kinase inhibitors. Among correctly predicted allosteric kinase sites correspond to structures of the receptor-interacting serine/threonine protein kinase 1 (RIPK1) bound to distinct necrostatin analogs (necrostatin-3 analog in PDB 4ITI and necrostatin-4 in PDB 4ITJ). The allosteric inhibitors in these structures stabilize RIPK1 in an inactive “DFG-out” conformation. Another correctly predicted allosteric sites correspond to type III kinase inhibitor bound to the catalytic domain of Focal Adhesion Kinase (FAK) in complex with a novel allosteric inhibitor (PDB 4EBW) and type III inhibitor bound to a human LIMK2 kinase domain (PDB 4TPT) (Figure 9A-D, top panel). The structural and mechanistic signature of these inhibitors is defined by several distinct characteristics. They bind to specific hydrophobic clefts (often termed the “B Pocket” or “back pocket”) adjacent to the ATP site. Unlike Type I/II inhibitors that extend into the back pocket from the ATP site, Type III inhibitors bind exclusively to this adjacent pocket. These allosteric binding pockets are predominantly hydrophobic, with well-defined geometries that accommodate the inhibitor through van der Waals interactions rather than hydrogen bonding networks typical of ATP-competitive inhibitors. Type III binding sites involve a relatively localized conformational change centered on the DFG motif flip, creating a pocket adjacent to the ATP site (Figure 9A-D). The correct classification of these Type III sites can be attributed to several key structural and evolutionary features that provide strong discriminative signals. First, these sites occupy well-defined hydrophobic pockets with clear geometric boundaries, making them readily distinguishable from the surrounding protein surface. Second, the DFG-out conformation creates a distinct structural signature that is captured by both structural and sequence-based representations. Third, while Type III sites are allosteric, they are located in regions that exhibit moderate sequence conservation across kinase families—sufficient to generate detectable evolutionary signals, unlike the more heterogeneous allosteric sites found in regulatory loops. Fourth, the localized nature of the conformational change means that these sites are relatively well-represented in crystallographic structures, providing consistent training examples. These features collectively explain why the model can identify these four Type III sites even with the limited allosteric training data available in the standalone KinSite dataset.

**Figure 9.**
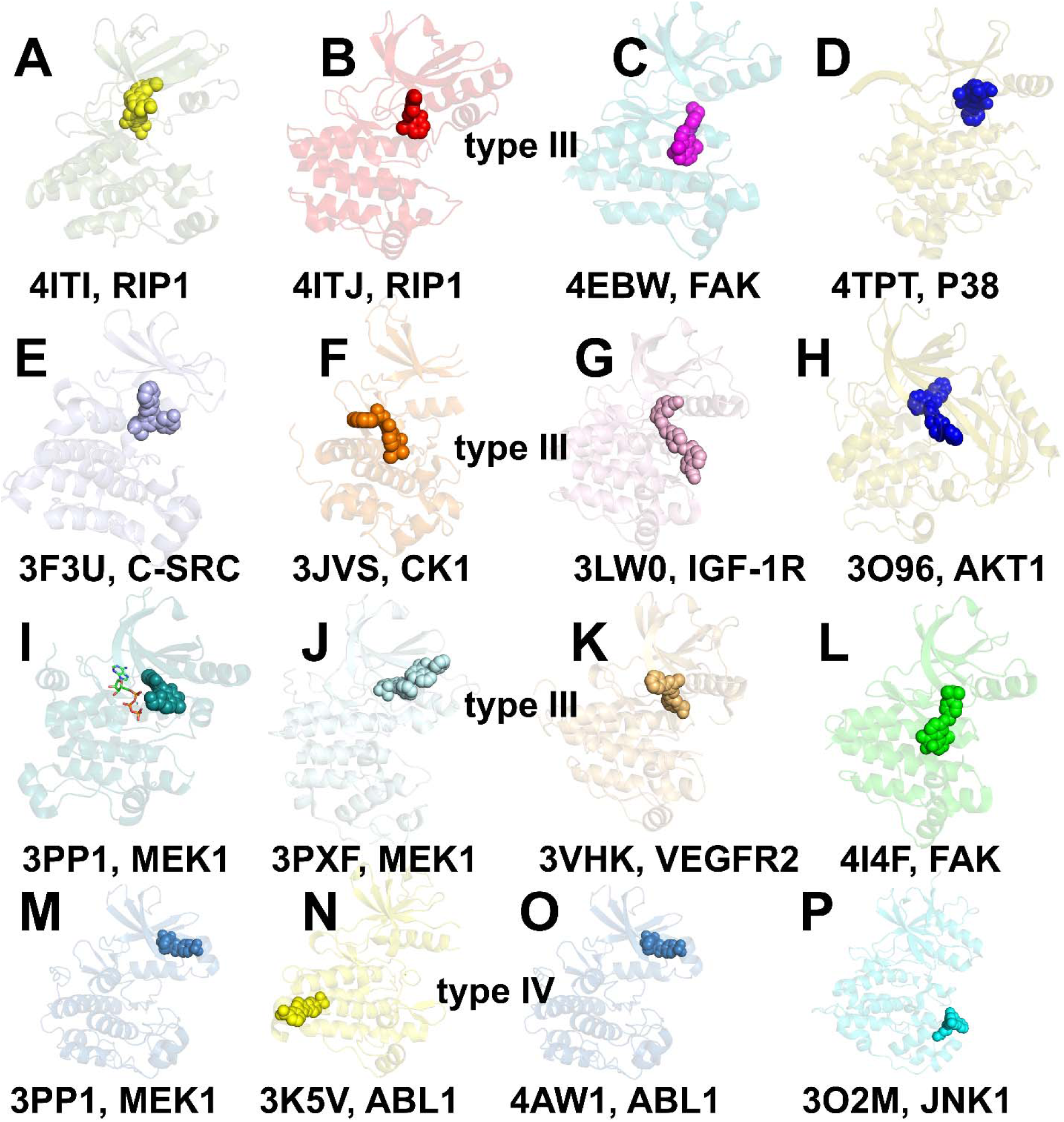
Structural and evolutionary comparison of kinase binding site types. The figure is organized into three panels illustrating the hierarchy of allosteric site detectability. (Top panel) Type III allosteric inhibitors correctly classified in the standalone KinSite dataset (4ITI, 4ITJ, 4EBW, 4TPT). These sites bind to hydrophobic pockets adjacent to the ATP site (B Pocket), with DFG-out conformational state and moderate sequence conservation. (Middle panel) Additional Type III allosteric sites recovered exclusively in the merged Dual-Site dataset, including examples from C-SRC (3F3U), CK1 (3JVS), IGF-1R (3LW0), AKT1 (3O96), MEK1 (3PP1, 3PXF), VEGFR2 (3VHK), and FAK (414F). These sites share structural features with the initial Type III sites and become detectable with structural regularization from PPI-Site. (Bottom panel) Type IV allosteric inhibitors that could not be recovered in any dataset (3K5V, 4AW1, 3O2M). These sites bind to distal regulatory regions (myristoyl pocket, docking interfaces) and exhibit high conformational flexibility, weak sequence conservation, and lack recurrent structural patterns. The hierarchy of detectability—orthosteric > Type III > Type IV—reflects the progressive loss of structural definition and evolutionary constraint as sites move from the catalytic core to distal regulatory regions.

On the other hand, the TN instances (correctly predicted orthosteric sites in our definition) represent orthosteric ATP-competitive inhibitors spanning multiple kinase families including These orthosteric sites share a common architecture: the ATP-binding cleft, defined by the conserved hinge region, the DFG motif, and the regulatory spine (R-spine). This region is under intense purifying selection due to its essential role in ATP binding and phosphoryl transfer, resulting in strong sequence conservation across the kinome. The diversity of kinase families represented in the TN set underscores the conserved nature of the ATP-binding cleft, which creates a strong and consistent discriminative signal that the model can reliably detect.

The interesting insight comes from comparing the standalone KinSite predictions with those from the merged Dual-Site dataset (PPI-Site + KinSite). The merged dataset produces two significant effects. First, it recovers 145 additional orthosteric sites that were missed in the standalone KinSite dataset. These newly recovered TN instances are almost exclusively Type I and Type II ATP-competitive kinase inhibitors binding to the canonical ATP-binding cleft. The PPI-Site dataset provides structural regularization by introducing structurally diverse orthosteric pockets from non-kinase proteins—particularly nucleotide-binding folds and Rossmann-like domains that share features with kinase ATP sites. In the standalone KinSite dataset, the extreme class imbalance (ρ ≈ 51.1) biases the model toward the majority class, causing some orthosteric sites to be misclassified. PPI-Site introduces additional orthosteric examples that help the model build a more robust representation of orthosteric site geometry, thereby improving discrimination in the kinase context. Second, the merged dataset recovers additional Type III allosteric kinase inhibitors among the TP predictions (Figure 9E-L, middle panels). These newly recovered sites including examples from C-SRC (3F3U), CK1 (3JVS), IGF-1R (3LW0), AKT1 (3O96), MEK1 (3PP1, 3PXF), VEGFR2 (3VHK), and FAK (414F) are structurally and mechanistically similar to the four initially classified sites. They share the same defining features: defined hydrophobic pocket geometry adjacent to the ATP site, DFG-out conformational state, and moderate sequence conservation. Because they share structural features with the initial Type III sites, the model can generalize to these additional examples when provided with structural regularization from PPI-Site. The structural diversity of PPI-Site exposes the model to a broader range of pocket geometries, enabling it to recognize the common features of Type III sites across different kinase families.

However, the merged dataset fails to robustly recover Type IV allosteric kinase inhibitors (Figure 9M-O, bottom panel). Type IV inhibitors bind to a regulatory, allosteric pocket that is completely distal (far removed) from both the standard ATP-binding site and the substrate-binding cleft. Validated PDB structures containing Type IV allosteric kinase inhibitors include BCR-ABL1 myristoyl pocket inhibitors (3K5V, 4AW1), MEK1 allosteric inhibitors (3PP1), and JNK1 distal pocket inhibitors (3O2M). Despite the availability of these structures, the model cannot recover them—not because of dataset composition or class imbalance, but because of fundamental challenges inherent to Type IV sites. The bottom panel of Figure 9 illustrates these distal binding sites, highlighting their spatial separation from the ATP-binding cleft and their location in conformationally flexible regulatory regions. The failure to recover Type IV inhibitors stems from several interconnected factors. First, Type IV binding sites are located in conformationally flexible regulatory regions—myristoyl pockets, PIF pockets, activation loops, and docking interfaces—that undergo large-scale conformational changes during kinase activation and regulatory transitions. Unlike the well-defined ATP-binding cleft or the adjacent B Pocket, Type IV sites lack a stable, conserved three-dimensional structure, making them difficult to capture from a single static conformation. Second, these sites exhibit weak or absent sequence conservation across kinase families, being often lineage-specific or unique to particular kinases. Unlike Type III sites, which have moderate sequence conservation, Type IV sites are characterized by neutral or weak evolutionary constraints. Third, because Type IV sites are diverse, lineage-specific, and structurally plastic, they do not provide recurrent structural or sequence patterns that machine learning models can learn. Each Type IV site is essentially a unique regulatory feature of its specific kinase, making it difficult to generalize from limited training examples. Fourth, true Type IV inhibitors are rare in structural databases, with only a handful of validated PDB entries representing a few kinase families with diverse binding modes and pocket geometries.

These findings align with our previous studies which examined PLM-based prediction of kinase binding sites^31^, where we demonstrated that orthosteric ATP-binding sites occupy minimally frustrated, evolutionarily constrained regions that generate strong sequence signatures, whereas allosteric sites reside in neutrally frustrated, conformationally plastic zones that lack consistent patterns. The present results extend this framework: Type III sites, despite being allosteric, retain enough structural definition and moderate evolutionary constraint to be detectable with structural regularization; Type IV sites, with their extreme conformational plasticity and evolutionary lability, remain beyond the reach of current multimodal representations. The inability to recover Type IV inhibitors regardless of dataset composition underscores that some allosteric sites are fundamentally challenging due to their physical and evolutionary properties, not due to limitations in model architecture or training data.

### A Biological Framework for Separability Regimes

The structural-functional analysis, integrating the conformational ambiguity of PPI-Site (Figure 8) with the evolutionary dichotomy of KinSite (Figure 9), reveals a coherent biological framework underlying the three separability regimes. In the low-separability PPI-Site regime, allosteric and orthosteric sites share the same protein family context, structural architecture, and ligand-binding functionality, differing primarily in their spatial relationship to protein–protein interaction interfaces. This conformational distinction is captured only by pocket geometry, while sequence and text modalities introduce noise rather than complementary information.

In the intermediate KinSite regime, three distinct site categories emerge with different detectability levels. Orthosteric ATP-binding clefts, with their strong sequence conservation and defined geometry, provide the strongest discriminative signal. Type III sites, located in hydrophobic pockets adjacent to the ATP site, retain enough structural definition and moderate evolutionary constraint to be detectable, particularly with structural regularization. Type IV sites, binding to distal regulatory regions, exhibit extreme conformational plasticity and evolutionary lability that preclude robust detection. The extreme class imbalance adds a statistical challenge, yet the underlying evolutionary signals provide consistent discriminative features across architectures.

In the high-separability AlloDiverse regime, the broad structural and functional heterogeneity of allosteric sites spanning diverse protein families creates a strong discriminative signal that multiple modalities can capture. The diversity of allosteric architectures and regulatory mechanisms overwhelms any shared features with orthosteric sites, enabling near-ceiling performance across most evaluated configurations. This progression from PPI-Site to KinSite to AlloDiverse represents a systematic increase in the biological distance between regulatory classes, which in turn determines the utility of different modalities for discrimination. The synergistic recovery in the Dual-Site dataset demonstrates that structural diversity and evolutionary signals are complementary sources of discriminative information. Neither alone is sufficient for robust discrimination across all contexts, but together they enable the model to resolve ambiguous cases. However, the persistent failure to recover Type IV inhibitors highlights that some allosteric sites remain challenging due to their intrinsic structural plasticity and evolutionary lability rather than limitations in dataset composition or model architecture.

## Discussion

The systematic characterization of multimodal protein foundation models for allosteric site prediction reveals three principal findings with significant implications for computational biology and drug discovery. First, dataset composition is the dominant determinant of predictive performance. Our variance decomposition analysis demonstrates that dataset identity accounts for 63.7% of the explainable variance in ROC-AUC, dwarfing the contributions of embedding composition (1.4%) and architectural choice (3.4%). This result challenges the prevailing emphasis on architectural innovation as the primary driver of progress in binding site prediction. The intrinsic characteristics of the training data—structural diversity, conformational heterogeneity, class balance, and the biological relationship between allosteric and orthosteric sites—exert a far greater influence on prediction success than model design choices. This finding does not diminish the importance of model architecture; rather, it reframes the central challenge of allosteric site prediction: the primary obstacle is not architectural sophistication but the availability and composition of training data that adequately capture the biological signals distinguishing regulatory classes. The identification of three distinct separability regimes provides a framework for understanding when multimodal integration is beneficial. In the low-separability PPI-Site regime, allosteric and orthosteric sites share structural and evolutionary contexts and are distinguished primarily by conformational differences. Here, pocket geometry alone is the most informative modality, and additional modalities introduce noise rather than complementary signal. This counterintuitive finding—that adding more information degrades performance—reflects the biological reality that when regulatory classes share evolutionary and functional contexts, global sequence and text signals provide variance rather than discrimination. In the intermediate KinSite regime, evolutionary constraint differentially marks orthosteric ATP-binding sites, making sequence embeddings essential and text annotations necessary for functional context. In the high-separability AlloDiverse regime, allosteric sites span broad structural and functional diversity, enabling multiple modalities to contribute synergistically to near-ceiling performance. This progression from low to high separability reveals that the value of multimodal integration is not absolute but depends critically on the biological distance between the classes being discriminated.

Beyond these empirical observations, our study establishes a general analytical framework for systematically profiling multimodal protein foundation models across distinct biological regimes. Rather than treating multimodal models solely as predictive tools, this approach uses them as scientific instruments to dissect how evolutionary, structural, functional, and dynamical information contribute to discrimination under different biological contexts. By integrating controlled embedding ablations, encoder-level comparisons, and variance decomposition, the framework disentangles the influence of dataset characteristics from that of model design, providing a principled basis for interpreting multimodal representations beyond conventional benchmarking. Although demonstrated here using OneProt, the approach is model-agnostic and readily transferable to other multimodal protein foundation models and biological prediction tasks. The framework’s modular design—separating encoder-level (what modalities shape the latent space) from embedding-level (what features are used for classification)—enables systematic attribution of performance gains to specific biological signals, a critical capability for building interpretable and trustworthy predictive models.

Second, modality contributions are fundamentally context-dependent and reflect the underlying biological organization of binding sites. The structural-functional analysis reveals that discriminative signals vary systematically with protein family context and binding site architecture. In PPI-Site, allosteric and orthosteric sites share Rossmann fold architectures and SAM-binding functionality, differing primarily in their spatial relationship to protein–protein interaction interfaces. Pocket geometry captures this conformational signal, while sequence and text modalities provide little additional discrimination. This observation reflects the biological reality that the same protein, the same ligand-binding pocket, and often the same set of residues can be classified as either allosteric or orthosteric depending on subtle conformational arrangements. The near-random performance across all architectures in this regime—and the fact that pocket-only embeddings slightly outperform multimodal combinations—underscores a fundamental challenge: when regulatory classes are defined by spatial relationships rather than intrinsic molecular differences, the discriminative signal is weak and conformational in nature.

In the kinase regime, a more nuanced picture emerges that reveals a hierarchy of site detectability. Orthosteric ATP-binding clefts, with their strong sequence conservation and defined pocket geometry, provide the most robust discriminative signal. These sites are under intense purifying selection due to their essential catalytic role, generating strong and unambiguous sequence signatures that protein language models can readily detect. Type III allosteric sites, located in hydrophobic pockets adjacent to the ATP site, retain enough structural definition and moderate evolutionary constraint to be detectable, particularly when structural regularization from diverse protein families is available. The correct classification of these sites benefits from the DFG-out conformational signature and the defined hydrophobic pocket geometry that distinguishes them from the surrounding protein surface. In contrast, Type IV allosteric sites, which bind to distal regulatory regions such as myristoyl pockets, PIF pockets, and activation loop docking interfaces, exhibit extreme conformational plasticity and evolutionary lability that preclude robust detection regardless of dataset composition. These sites reside in neutrally frustrated regions that permit sequence drift and conformational variation, enabling lineage-specific regulatory tuning but producing weak or absent sequence signals. The failure to recover Type IV inhibitors even in the merged dataset underscores that some allosteric sites are fundamentally challenging due to their physical properties—structural plasticity and evolutionary lability—rather than limitations in model architecture or training data. This hierarchy—orthosteric > Type III > Type IV in terms of detectability—reflects the progressive loss of structural definition and evolutionary constraint as sites move from the catalytic core to distal regulatory regions.

The largest performance gains in the kinase regime arise from incorporating functional text annotations, which capture protein-specific regulatory mechanisms and biological context that sequence embeddings alone cannot provide. UniProt-derived text annotations encode information about protein family, pathway, subcellular localization, and regulatory mechanisms—knowledge that is essential for distinguishing allosteric sites with similar structural features but distinct biological functions. Sequence embeddings contribute information on evolutionary constraint, whereas text embeddings provide the functional specialization that most effectively distinguishes allosteric from orthosteric kinase binding sites. The extreme class imbalance in KinSite (ρ ≈ 51.1) further amplifies the importance of modality selection, as the model must rely heavily on discriminative signals to recover rare positive examples.

Third, architectural complexity provides diminishing returns once sufficient biological signal is present. Structural and dynamical encoders contribute meaningfully in intermediate-separability regimes, but their impact diminishes in both low-separability settings (where the underlying signal is too weak for any architecture to exploit) and high-separability settings (where multiple architectures converge to near-ceiling performance). The convergence of different architectural configurations in AlloDiverse indicates that the critical factor is the inclusion of structural information rather than the specific structural encoder used to capture it. This pattern was particularly evident in Dual-Site, where the addition of structural encoders consistently improved performance, but no single structural representation strategy dominated. The structure-token encoder, which captures global fold-level organization, provided the most robust performance across regimes, suggesting that broader structural context is more informative than fine-grained local geometry for distinguishing allosteric from orthosteric sites.

A key insight emerging from this study is that combining datasets with complementary biological characteristics yields models that perform better than those trained on either dataset alone. The Dual-Site dataset recovered numerous cases missed in standalone training—including both additional orthosteric sites (Type I and II inhibitors that were previously misclassified) and additional Type III allosteric sites—demonstrating that structural diversity and evolutionary signals are complementary sources of information. The structural diversity of PPI-Site provides a regularization effect, improving the model’s ability to recognize both orthosteric and Type III allosteric sites by exposing it to a broader range of pocket geometries. Conversely, the evolutionary signals from KinSite help resolve ambiguous cases in PPI-Site where structural features alone were insufficient. However, even this synergistic combination could not recover Type IV inhibitors, reinforcing that some allosteric sites remain beyond the reach of current representations due to their intrinsic biophysical properties.

These findings have direct implications for practical applications in drug discovery. For kinase drug discovery, where orthosteric ATP sites are well-characterized but allosteric sites offer opportunities for selective modulation, models should prioritize sequence and functional embeddings that capture strong evolutionary signals and regulatory context. The ability to detect Type III sites with structural regularization suggests that combining diverse training data can improve identification of this therapeutically relevant class. However, the persistent failure to recover Type IV inhibitors highlights the need for dynamics-aware representations that capture conformational ensembles and regulatory plasticity. For protein–protein interaction targets, where allosteric and orthosteric sites may share structural contexts, geometric representations capable of capturing subtle conformational differences are essential. For proteome-wide screening, where allosteric sites may arise from diverse structural and functional contexts, multimodal integration becomes critical. The field should therefore devote increased attention to systematic dataset curation and characterization. The fact that dataset identity explains nearly two-thirds of performance variance underscores that progress in allosteric site prediction will come not from architectural innovation alone but from careful design of training datasets that span relevant biological regimes. Rather than training on a single representative dataset, models should be trained on diverse datasets that encompass different structural contexts, evolutionary constraints, and regulatory mechanisms. Future directions include developing adaptive models that dynamically weight modalities based on biological context, incorporating conformational ensembles to improve discrimination in low-separability regimes, extending the analysis to cryptic and transient binding sites, and integrating physics-based constraints to enhance physical interpretability and generalization. By establishing the dataset-driven nature of separability regimes in allosteric site prediction, this study provides a foundation for the development of more robust, context-aware, and physically grounded multimodal models for allosteric drug discovery.

### Conclusions

In this study, we systematically investigated the determinants of multimodal protein foundation model performance for distinguishing allosteric from orthosteric competitive binding sites across datasets spanning distinct biological regimes. Our analysis reveals that dataset composition—not architectural complexity—is the dominant factor governing predictive performance, accounting for 63.7% of explainable variance in ROC-AUC. We identify three distinct separability regimes: a low-separability regime where current representations are unable to reliably distinguish regulatory classes (PPI-Site); an intermediate regime where multimodal integration substantially improves performance (KinSite, Dual-Site); and a high-separability regime where most architectures converge to near-ceiling performance (AlloDiverse). This progression reveals that the value of multimodal integration is not absolute but depends critically on the biological distance between the classes being discriminated.

The utility of different biological modalities is regime-dependent and reflects the underlying organization of binding sites. In the PPI-Site regime, pocket geometry dominates because allosteric and orthosteric sites share structural and evolutionary contexts, differing primarily in their spatial relationship to protein–protein interaction interfaces. In the KinSite regime, functional text annotations provide the strongest complementary information by encoding protein-specific regulatory mechanisms, while sequence embeddings capture differential evolutionary constraints. The structural-functional analysis reveals a hierarchy of site detectability within the kinase regime: orthosteric ATP-binding clefts provide the strongest discriminative signal; Type III allosteric sites are detectable with structural regularization; and Type IV distal allosteric sites remain beyond current multimodal representations due to structural plasticity, evolutionary lability, and lack of recurrent patterns. This hierarchy reflects the progressive loss of structural definition and evolutionary constraint as sites move from the catalytic core to distal regulatory regions.

The Dual-Site regime demonstrates that structural diversity and evolutionary signals are complementary: structural diversity from PPI-Site regularizes the model, improving recognition of both orthosteric sites and Type III allosteric sites, while evolutionary signals from KinSite resolve ambiguous PPI-Site cases. However, even this synergistic combination cannot recover Type IV inhibitors, underscoring that some allosteric sites are fundamentally challenging due to their intrinsic physical properties. The AlloDiverse regime establishes that comprehensive coverage of allosteric site diversity creates a strong discriminative signal that multiple modalities can capture.

These findings establish a principled framework for understanding when and why multimodal integration benefits allostery prediction, moving beyond conventional benchmarking to provide systematic characterization of learnability regimes. The progression from PPI-Site to KinSite to Dual-Site to AlloDiverse represents a systematic increase in allosteric diversity and structural heterogeneity. Each regime reveals different aspects of the allosteric prediction challenge: PPI-Site highlights the difficulty of conformational discrimination within shared structural contexts; KinSite demonstrates the importance of evolutionary signals in structurally conserved families and reveals the hierarchy of allosteric site detectability; Dual-Site shows the synergistic benefits of combining complementary datasets; and AlloDiverse establishes that comprehensive allosteric diversity is the most effective path to robust prediction.

Future work should focus on developing adaptive models that dynamically weight modalities based on biological context, incorporating conformational ensembles and dynamics-aware representations to improve discrimination in challenging regimes, and integrating physics-based constraints to enhance physical interpretability and generalization. By establishing the dataset-driven nature of separability regimes in allosteric site prediction, this study provides a foundation for the development of more robust, context-aware, and physically grounded multimodal models for allosteric drug discovery.

## Author Contributions

Conceptualization, G.V., A.B.; Methodology, G.V., A.B. Software, A.B. Validation, A.B., G.V.; Formal analysis, A.B., G.V.; Investigation, A.B., G.V.; Resources, A.B.;. Data curation, A.B., G.V.; Writing—original draft preparation, A.B., G.V.; Writing—review and editing, A.B., G.V.; Visualization, A.B., G.V.;. Supervision G.V.; Project administration, G.V.;. Funding acquisition, A.B., G.V.

All authors have read and agreed to their individual contributions. All authors have approved the submission of this manuscript, and it has been submitted exclusively to Journal of Chemical Information and Modeling.

## Conflicts of Interest

The authors declare no conflict of interest. The funders had no role in the design of the study; in the collection, analyses, or interpretation of data; in the writing of the manuscript; or in the decision to publish the results.

## Data Availability Statement

All source code, processed datasets, pretrained model checkpoints, and the computational environment required to reproduce the results of this study are publicly available through the OneProt Models GitHub organization (https://github.com/oneprot-models) and the accompanying Zenodo archive (https://doi.org/10.5281/zenodo.20997998).

The GitHub repository oneprot-embeddings (https://github.com/oneprot-models/oneprot-embeddings) contains the complete software implementation used in this work. The repository provides the OneProt framework for multimodal protein embedding extraction together with utilities for downstream prediction and evaluation. The allosteric-site prediction workflow is organized under src/allostery, where generate_pockets/ contains scripts for reconstructing fixed-size binding pocket datasets from the original PPI-Site, KinSite, and ASD resources; extract_embeddings/ contains workflows for generating multimodal OneProt embeddings from the reconstructed pockets using the sequence-, structure-, graph-, molecular dynamics-, and multimodal encoder variants evaluated in this study; plotting/ contains the scripts used to generate all figures and statistical analyses presented in the manuscript; and utils/ provides shared utilities for dataset handling, preprocessing, evaluation, and reproducibility. Environment specifications, including all required software dependencies and package versions, are provided in the directory (https://github.com/oneprot-models/oneprot-embeddings/tree/main/environment), while a fully configured Apptainer/Singularity runtime environment is distributed through the accompanying Zenodo archive as oneprot_container.sif. Complete documentation, configuration files, and example commands required to reproduce the workflow are provided in the repository README.

The archive also includes checkpoints.zip, containing the pretrained downstream model checkpoints corresponding to all seven OneProt embedding ablations evaluated in this study.

Downstream fine-tuning using the OneProt embeddings is performed via the original unmodified OneProt framework and is executed through the scripts src/saprot_fit_mlp.py and src/saprot_fit_mlp_balanced.py for the case when the data needs to be balanced during fine-tuning. The whole procedure is described in general OneProt README.

The accompanying Zenodo archive contains all processed resources required to reproduce the reported experiments.

The HDF5 files (binding_pockets_allosteric.h5, binding_pockets_orthosteric_competitive.h5, allosteric_pockets_kinsite.h5, competitive_pockets_kinsite.h5, and ASD_binding_pockets.h5) contain reconstructed atom-level binding pocket structures used for OneProt pocket embedding extraction for PPI-site allosteric, PPI-site orthosteric, KinSite allosteric, KinSite orthosteric and ASD derived allosteric pockets, respectively. Unless stated otherwise, pockets were generated by retaining the 100 residues closest to the annotated ligand or reconstructed pocket center while preserving all atoms belonging to the selected residues.

The archive further contains all train, validation, and test partitions used throughout this study. PPI-site_splits.zip and KinSite_splits.zip provide MMseqs2-based data partitions generated using a 30% sequence identity threshold to prevent homologous proteins from appearing across dataset splits. AlloDiverse_splits.zip contains the benchmark splits used in this work, where the positive class is derived from the ASD dataset and the validation and test sets correspond to a random 50:50 partition of the original ASD test set. ASD_original_split.zip contains the original train/test partition from the published ASD benchmark, while PPI-site_KinSite_original.zip contains the processed PPI-Site and KinSite datasets obtained from the original publications prior to sequence clustering and repartitioning.

## Acknowledgments

A.B. is supported by the Helmholtz Association Initiative and Networking Fund in the frame of Helmholtz AI as well as by the Helmholtz Foundation Model Initiative within the project “PROFOUND.” A.B. gratefully acknowledges the Gauss Centre for Supercomputing e.V. (www.gauss-centre.eu) for funding this project by providing computing time on the GCS Supercomputer JUWELS at Jülich Supercomputing Centre (JSC).

This research was funded by the National Institutes of Health under Award 1R01AI181600-01, 5R01AI181600-02 and Subaward 6069-SC24-11 to G.V. G.V acknowledges support from Schmid College of Science and Technology at Chapman University for providing computing resources at the Keck Center for Science and Engineering.

## Supporting Information

Supporting Text S1–S4, Figures S1–S5, and Tables S1–S2 provide additional methods, implementation details, dataset descriptions, and comprehensive performance analyses supporting the results presented in the main manuscript.

**Supporting Text S1–S4** Supporting Text S1 describes the OneProt pre-training architecture, including the structural, pocket, sequence, text, and molecular dynamics encoders, as well as multimodal latent-space alignment. Supporting Text S2 details binding-pocket construction, structural preprocessing, and pocket overlap analyses for the benchmark datasets. Supporting Text S3 describes the downstream allostery classification framework, including hyperparameter optimization, performance metrics, and strategies for handling class imbalance. Supporting Text S4 summarizes implementation details, computational resources, and reproducibility information.

**Figures S1–S5 Figure S1** presents receiver operating characteristic (ROC) curves across OneProt encoder architectures, embedding combinations, and allosteric classification datasets. Figure S2 shows precision–recall (PR) curves across encoder architectures under different class-imbalance regimes. Figure S3 summarizes mean ROC-AUC values for all combinations of datasets, embedding configurations, and OneProt encoder architectures. Figure S4 presents threshold-dependent sensitivity across datasets and embedding combinations. Figure S5 presents threshold-dependent specificity across datasets and embedding combinations.

**Tables S1–S2:** Table S1 reports the complete mean ROC-AUC values (± standard deviation across independent runs) for all combinations of datasets, embedding configurations, and OneProt encoder architectures. Table S2 reports the corresponding mean PR-AUC values (± standard deviation across independent runs) for all evaluated configurations, providing a quantitative basis for assessing the impact of class imbalance and modality selection on allosteric site discrimination.

## Notes

### Competing Interest Statement

The authors have declared no competing interest.

