## Supporting Text S1-S4, Supporting Figures S1-S5 and Supporting Tables S1,S2. for "Integrated Framework for Probing Multimodal Protein Foundation Models with Structure-Functional Interpretability Analysis in Detection of Allosteric Binding Sites": SUPPORTING_INFORMATION_BIORXIV.docx

**Text S1. Pre-training Architectures**

**Text S1.1 Structural Encoders**

Structural representations were generated using two complementary structural encoders integrated within OneProt: a graph-based structural encoder and a token-based structural encoder. These encoders capture complementary aspects of protein three-dimensional organization across multiple spatial scales.

Graph-based structural representations were obtained using the ProNet architecture, a hierarchical graph neural network that learns protein representations at three complementary levels of structural granularity: amino-acid, backbone, and all-atom. Given a protein structure containing $L$amino-acid residues, the protein is represented as a graph $G=(V,E,P)$, where $V$denotes the set of residue nodes, $E$represents the edges describing covalent and spatial relationships between residues, and $P$contains the structural information associated with the protein. Through hierarchical modeling at the amino-acid, backbone, and all-atom levels, ProNet captures complementary structural features spanning local geometric environments and higher-order protein organization.

At each level, node representations are iteratively refined using message passing according to the general update rule:

$$m_{v}^{\left( t \right)}=\text{AGGREGATE}_{u\in\mathcal{N(}v)}\left( \text{MESSAGE}^{\left( t \right)}(h_{v}^{\left( t \right)},h_{u}^{\left( t \right)},e_{uv}) \right)$$

$$h_{v}^{\left( t + 1 \right)}=\text{UPDATE}^{\left( t \right)}(h_{v}^{\left( t \right)},m_{v}^{\left( t \right)})$$

where $\mathcal{N(}v)$denotes the neighborhood of node $v$, $e_{uv}$represents geometric edge features derived from the structural information contained in $P$, and MESSAGE, AGGREGATE, and UPDATE are learnable functions at layer $t$.

Through hierarchical message passing across the amino-acid, backbone, and all-atom representations, ProNet learns structural embeddings that capture fine-grained local interactions together with the higher-order three-dimensional organization of the complete protein fold. Within OneProt, these graph embeddings are projected into the shared multimodal latent space and aligned with sequence, pocket, text, and MD representations during contrastive pretraining.

Token-based structural representations were generated using Foldseek structural tokens processed with an ESM2-35M transformer encoder. Foldseek converts protein structures into sequence-like representations by discretizing local three-dimensional residue environments into a learned structural alphabet. Each structural token encodes the geometric relationship between a residue and its nearest spatial neighbor, capturing local tertiary structural information in a discrete representation.

The resulting structural token sequence:

$$S_{struct}=(t_{1},t_{2},\ldots,t_{L})$$

where each token $t_{i}\mathcal{\in T}$belongs to a learned structural alphabet of size $\mid\mathcal{T}\mid=20$, is processed by an ESM2 transformer encoder to produce contextual structural embeddings:

$$H_{struct}=\text{ESM2}(S_{struct})\in\mathbb{R}^{L\times1280}$$

In contrast to graph-based structural representations, structural-token embeddings model proteins as sequences of discretized local structural environments, enabling transformer self-attention to capture long-range dependencies and global fold organization while preserving information about local tertiary geometry.

**Text S1.2 Pocket Encoder**

Pocket embeddings were obtained using the binding-site encoder integrated within the OneProt framework. Binding pockets are represented as three-dimensional residue graphs and processed using a graph neural network based on the ProNet architecture. Graph-based representations are particularly suitable for binding-site modeling because they preserve spatial residue relationships and local geometric organization while remaining invariant to global coordinate transformations.

For each binding site, a binding-pocket center $\mathbf{c}\in\mathbb{R}^{3}$was defined. For each residue $i$, the residue centroid was computed as the average Cartesian coordinate of all atoms belonging to that residue:

$$\mathbf{r}_{i}=\frac{1}{\mid A_{i}\mid}\sum_{a\in A_{i}} \mathbf{x}_{a}$$

where $A_{i}$denotes the set of atoms comprising residue $i$, and $\mathbf{x}_{a}\in\mathbb{R}^{3}$is the Cartesian coordinate of atom $a$.

Residues were ranked according to their Euclidean distance from the binding-site center:

$$d_{i}=\parallel\mathbf{r}_{i}-\mathbf{c}\parallel_{2}$$

and the $K=100$nearest residues were selected to define the binding pocket.

The resulting pocket was represented as a residue graph:

$$G_{pocket}=(V_{pocket},E_{pocket},P_{pocket})$$

where $V_{pocket}$denotes the set of selected residue nodes, $E_{pocket}$represents the graph connectivity defined by the ProNet encoder, and $P_{pocket}$contains the associated three-dimensional structural information.

For each residue, graph node features included the residue identity, backbone atom coordinates (N, Cα, and C), backbone torsion-angle embeddings, and side-chain torsion-angle embeddings. Backbone torsion angles were encoded using sine and cosine transformations:

$$\mathbf{b}_{i}=[\cos\phi_{i},\cos\psi_{i},\cos\omega_{i},\sin\phi_{i},\sin\psi_{i},\sin\omega_{i}]$$

whereas side-chain torsion angles were represented as:

$$\mathbf{s}_{i}=[\sin\chi_{i,1},\sin\chi_{i,2},\sin\chi_{i,3},\sin\chi_{i,4},\cos\chi_{i,1},\cos\chi_{i,2},\cos\chi_{i,3},\cos\chi_{i,4}]$$

The pocket graph was then processed by the ProNet graph encoder to obtain a latent structural representation:

$$\mathbf{h}_{pocket}=f_{ProNet}(G_{pocket})$$

where $f_{ProNet}$denotes the hierarchical graph neural network operating on the residue graph.

Within OneProt, this latent representation was further transformed by a learnable projection head and L2 normalization:

$$\mathbf{z}_{pocket}=\text{Normalize}(\text{Proj}(\mathbf{h}_{pocket}))$$

where Proj denotes the learnable projection module and Normalize represents L2 normalization. The resulting embedding $\mathbf{z}_{pocket}$constitutes the final pocket representation used for multimodal contrastive alignment.

**Text S1.3 Text Encoder**

Textual representations were derived from UniProt functional annotations and processed using the BiomedBERT encoder integrated within OneProt model. BiomedBERT is a domain-adapted BERT model pretrained on biomedical literature, achieving improved performance on biomedical natural language processing tasks.

UniProt annotations consist of concatenated text fields including protein names and families, organism information, function descriptions, subcellular localization, and pathway and similarity information. Given a textual annotation $T=(w_{1},w_{2},\ldots,w_{M})$of length $M$tokens, the BiomedBERT encoder produces contextualized representations:

$$H_{\text{text}}=\text{BiomedBERT}(T)\in\mathbb{R}^{M\times768}$$

The final text embedding is obtained through mean pooling across token positions:

$$\mathbf{h}_{\text{text}}=\frac{1}{M}\sum_{i=1}^{M} H_{\text{text}}[i]\in\mathbb{R}^{768}$$

The pooled representation is subsequently projected into the shared OneProt latent space through a trainable projection head:

$$\mathbf{z}_{\text{text}}=P_{\text{text}}(\mathbf{h}_{\text{text}})\in\mathbb{R}^{1024}$$

where $P_{\text{text}}$denotes the projection network. This modality captures semantic and functional biological information associated with protein activity, molecular function, and biochemical context. Previous OneProt studies demonstrated that inclusion of textual modalities substantially improves multimodal latent-space organization and downstream transfer performance.

**Text S1.4 MD Encoder**

We used the extended OneProt architecture, which incorporates MD trajectory representations. Because allosteric regulation is fundamentally associated with conformational fluctuations, ensemble redistribution, and long-range dynamical coupling, static experimental structures may incompletely capture the conformational landscapes underlying regulatory behavior. MD trajectories therefore provide complementary information describing protein flexibility and time-dependent structural variability beyond single static conformations.

The MD modality was implemented using the latent MD encoder derived from MDGen, a transformer-based generative framework for MD trajectory modeling. Unlike conventional structure encoders operating on individual conformations, the MDGen encoder processes full MD trajectories as temporally ordered structural ensembles.

Given an input trajectory $X_{MD}=\{X_{1},X_{2},\ldots,X_{T}\}$comprising $T$simulation frames and $L$residues, each frame is represented using SE(3)-invariant roto-translational offsets together with backbone and side-chain torsion-angle representations. The encoder produces latent trajectory representations:

$$H_{MD}=f_{MDGen}(X_{MD})\in\mathbb{R}^{B\times T\times L\times D}$$

where $B$is the batch size, $T$is the number of trajectory frames, $L$is the number of residues, and $D$is the latent embedding dimension (1024).

Architecturally, the latent MD encoder is based on a Scalable Interpolant Transformer (SIT) framework augmented with long-context Hyena operators replacing conventional temporal attention layers, enabling efficient modeling of long-timescale temporal dependencies across molecular trajectories. Spatial features are processed through transformer-based structural layers, while temporal evolution is captured through Hyena-based sequence modeling across trajectory frames.

To obtain fixed-length embeddings compatible with the OneProt multimodal framework, temporal pooling across trajectory frames was first applied:

$$\mathbf{h}_{t,l}=\frac{1}{T}\sum_{t=1}^{T} H_{MD}[t,l]\in\mathbb{R}^{D}$$

followed by spatial pooling across residues:

$$\mathbf{h}_{MD}=\frac{1}{L}\sum_{l=1}^{L} \mathbf{h}_{t,l}\in\mathbb{R}^{1024}$$

The resulting pooled trajectory representations were subsequently projected into the shared OneProt latent space using modality-specific MLP projection heads, producing 1024-dimensional MD embeddings aligned with the remaining OneProt modalities.

MD trajectories used during pretraining were curated from mdCATH, GPCRmd, and ATLAS databases through the MDRepo repository. These datasets provide complementary coverage of protein families, conformational regimes, and simulation timescales, including membrane proteins, globular proteins, and structurally diverse protein folds. To reduce heterogeneity between simulation sources, trajectories were temporally standardized to a common sampling interval of $\Delta t=0.1$ns and converted into torsion-angle and rigid-frame representations compatible with the latent MD encoder.

**Text S1.5 Cross-Modal Representation Alignment and Transfer**

Modality-specific encoders within OneProt are jointly aligned using multimodal contrastive learning based on the InfoNCE objective, conceptually analogous to multimodal representation-learning frameworks such as CLIP and ImageBind. Embeddings derived from different biological modalities are projected into a shared latent space, where alignment is performed pairwise with the sequence modality acting as the central anchor representation.

For a batch of $N$protein samples, let $\mathbf{z}_{\text{seq}}^{\left( i \right)}$, $\mathbf{z}_{\text{pocket}}^{\left( i \right)}$, $\mathbf{z}_{\text{text}}^{\left( i \right)}$, $\mathbf{z}_{\text{struct}}^{\left( i \right)}$, and $\mathbf{z}_{\text{MD}}^{\left( i \right)}$denote the normalized embeddings for sample $i$from the sequence, pocket, text, structural, and MD modalities, respectively.

The contrastive loss for aligning sequence and pocket modalities is defined as:

$$\mathcal{L}_{\text{seq-pocket}}=-\frac{1}{N}\sum_{i=1}^{N} \log\frac{\exp(\mathbf{z}_{\text{seq}}^{\left( i \right)}\cdot\mathbf{z}_{\text{pocket}}^{\left( i \right)}/\tau)}{\sum_{j=1}^{N} \exp(\mathbf{z}_{\text{seq}}^{\left( i \right)}\cdot\mathbf{z}_{\text{pocket}}^{\left( j \right)}/\tau)}$$

where $\tau>0$is a temperature parameter controlling the sharpness of the softmax distribution, and the dot product $\mathbf{z}_{\text{seq}}^{\left( i \right)}\cdot\mathbf{z}_{\text{pocket}}^{\left( j \right)}$measures the cosine similarity between normalized embeddings.

The total multimodal alignment loss is the sum over all modality pairs anchored to the sequence modality:

$$\mathcal{L}_{\text{total}}=\sum_{m\mathcal{\in M}} \mathcal{L}_{\text{seq-}m}$$

where $\mathcal{M=\{}\text{pocket},\text{text},\text{struct},\text{MD}\}$.

This framework enables transfer of structural, functional, and dynamical information across heterogeneous biological modalities. During pretraining, all modality-specific encoders are jointly optimized to minimize $\mathcal{L}_{\text{total}}$, resulting in a shared latent space where embeddings from different modalities corresponding to the same protein are pulled together while those from different proteins are pushed apart.

Previous OneProt studies demonstrated that multimodally aligned representations substantially improve downstream transfer performance compared to their unimodal counterparts, indicating that biological information is propagated between modalities during pretraining. Consequently, modality-specific embeddings within OneProt represent enriched multimodal representations rather than isolated unimodal encodings.

The extent and nature of this cross-modal knowledge transfer depend on the specific encoder configuration participating in multimodal alignment. Previous OneProt studies and subsequent extensions incorporating MD representations demonstrated that different modality combinations produce distinct downstream transfer characteristics and latent-space organization properties. In the present study, these encoder-level ablations therefore provide a framework for probing how structural, functional, and dynamical modalities influence representations associated with allosteric regulation.

Although downstream classifiers operate only on extracted sequence, pocket, and text embeddings, additional modalities such as graph-based structural representations, structural tokens, and MD trajectories may still influence downstream prediction indirectly through their contribution to latent-space organization during multimodal pretraining.

**Text S2. Pocket Construction and Structural Processing**

Binding pocket representations were constructed following the preprocessing and multimodal representation framework used in the OneProt protein foundation model. OneProt aligns heterogeneous biological modalities—including protein sequence, three-dimensional structure, binding pockets, textual annotations, and MD trajectories—within a shared latent embedding space through modality-specific encoders and multimodal contrastive learning.

**Text S2.1 Pocket Construction for PPI-Site Dataset**

For the PPI-Site dataset, binding pockets were reconstructed from the cavity annotations provided in the original resource. Annotated ligands were first identified within the corresponding Protein Data Bank (PDB) structures, and the geometric centroid of the ligand atoms was used to define the pocket center:

$$\mathbf{c}_{\text{pocket}}=\frac{1}{N_{\text{lig}}}\sum_{i=1}^{N_{\text{lig}}} \mathbf{x}_{i}^{\text{lig}}$$

where $\left\{ \mathbf{x}_{i}^{\text{lig}}{\}}_{i=1}^{N_{\text{lig}}} \subset\mathbb{R}^{3} \right.$are the coordinates of ligand atoms.

For the associated protein chain, residue centroids were computed as the average coordinates of all atoms belonging to each standard amino-acid residue:

$$\mathbf{c}_{r}=\frac{1}{N_{r}}\sum_{a\in r} \mathbf{x}_{a}$$

where $N_{r}$is the number of atoms in residue $r$.

Residues were subsequently ranked according to the Euclidean distance between their centroids and the ligand center:

$$d(r,\mathbf{c}_{\text{pocket}})=\parallel\mathbf{c}_{r}-\mathbf{c}_{\text{pocket}}\parallel_{2}$$

and the $K=100$nearest residues were retained to construct fixed-size pocket representations:

$$\mathcal{R}_{\text{pocket}}=\{r_{\left( 1 \right)},r_{\left( 2 \right)},\ldots,r_{\left( K \right)}\}$$

where $r_{\left( 1 \right)},r_{\left( 2 \right)},\ldots$denote residues sorted by increasing distance to the pocket center. All atoms belonging to the selected residues were preserved, yielding atom-level pocket structures together with their associated residue sequences. The complete protein-chain sequence was stored separately for sequence-based representation extraction.

**Text S2.2 Pocket Overlap and Conformational Variability Analysis**

Because the PPI-Site dataset contains structurally resolved pocket annotations spanning both allosteric and orthosteric competitive binding regimes within shared protein contexts, it enables direct comparison of local binding environments across regulatory classes. Equivalent analyses were not performed on the AlloDiverse dataset because ASD pockets are reconstructed from residue-level annotations rather than experimentally resolved cavity definitions, preventing direct structural comparison of matched pocket pairs. Similarly, the KinSite dataset predominantly reflects highly conserved ATP-binding environments under strong class imbalance, limiting the interpretability of residue-overlap analyses across regulatory states.

To assess the extent of structural overlap between allosteric and orthosteric competitive binding environments in PPI-Site, we compared pockets across both classes for proteins sharing the same protein and chain identifiers. Comparisons were performed at both the residue-composition level and the atom-coordinate level.

**Text S2.3 Residue-Level Overlap Analysis**

Residue-level overlap was quantified using the Jaccard similarity coefficient. Given residue sets $A$and $B$corresponding to two pockets, the Jaccard similarity was defined as:

$$J(A,B)=\frac{\mid A\cap B\mid}{\mid A\cup B\mid}$$

where $\mid A\cap B\mid$denotes the number of shared residues and $\mid A\cup B\mid$denotes the total number of unique residues across both pockets.

In addition to Jaccard similarity, we computed the overlap coefficient:

$$O(A,B)=\frac{\mid A\cap B\mid}{\min(\mid A\mid,\mid B\mid)}$$

which normalizes the shared residue count by the size of the smaller pocket and is therefore more sensitive to partial containment relationships between pockets of different sizes.

**Text S2.4 Coordinate-Level Geometric Similarity**

To assess coordinate-level geometric similarity, atom-level nearest-neighbor statistics were computed using KD-tree nearest-neighbor search. Given atomic coordinate sets $X=\{\mathbf{x}_{i}{\}}_{i=1}^{N}$and $Y=\{\mathbf{y}_{i}{\}}_{i=1}^{M}$corresponding to two pockets, the fraction of atoms in pocket $X$located within a distance threshold $\tau$of any atom in pocket $Y$was defined as:

$$f_{X\to Y}=\frac{1}{N}\sum_{i=1}^{N} \mathbf{1}\left( \min_{j}\parallel\mathbf{x}_{i}-\mathbf{y}_{j}\parallel_{2}<\tau\right)$$

where $\mathbf{1}(\cdot)$denotes the indicator function and $\tau=4$Å was used as the spatial proximity threshold.

Continuous geometric similarity was further quantified using the mean nearest-neighbor atom distance:

$$d_{X\to Y}=\frac{1}{N}\sum_{i=1}^{N} \min_{j}\parallel\mathbf{x}_{i}-\mathbf{y}_{j}\parallel_{2}$$

Using this procedure, we identified 100 pocket pairs (<5% of the dataset) exhibiting identical residue composition ($J=1.0$), indicating that the same 100 residues were selected for both allosteric and orthosteric competitive pockets within the same protein structure. These cases arise when proteins are crystallized in distinct ligand-bound regulatory states, such that nearest-residue pocket extraction yields identical residue membership despite differing ligand identities and binding regimes.

Despite complete residue-level overlap, the corresponding pockets frequently exhibited substantial coordinate-level geometric divergence. Mean nearest-neighbor atom distances ranged from <0.1 Å to >18 Å, indicating that structurally similar residue environments can adopt markedly different conformations depending on ligand-binding state. These observations suggest that allosteric and orthosteric regulation may, in some cases, emerge from distinct conformational organizations within partially shared local structural environments rather than from entirely disjoint residue compositions. Given the relatively small number of such cases and their substantial coordinate-level variability, these instances were retained in the dataset.

**Text S2.5 Pocket Construction for ASD Dataset**

For the positive instances from ASD dataset, representative allosteric regions were reconstructed from residue-level annotations using spatial clustering and geometric pocket reconstruction. Residues annotated as allosteric were first mapped onto experimentally resolved protein structures using their residue identifiers. Each residue was represented by the three-dimensional coordinates of its Cα atom.

Spatially contiguous allosteric regions were identified using Density-Based Spatial Clustering of Applications with Noise (DBSCAN) which groups residues according to local spatial density without requiring a predefined number of clusters. DBSCAN was selected because it naturally accommodates disconnected and sparsely annotated allosteric regions frequently encountered in residue-level annotations.

Given a set of residue coordinates $\left\{ \mathbf{x}_{i}{\}}_{i=1}^{N} \subset\mathbb{R}^{3} \right.$, two residues $\mathbf{x}_{i}$and $\mathbf{x}_{j}$are considered neighbors if their Euclidean distance satisfies:

$$\parallel\mathbf{x}_{i}-\mathbf{x}_{j}\parallel_{2}<\varepsilon$$

where $\varepsilon$defines the neighborhood radius. In this study we used $\varepsilon=10$Å and minimum number of samples minPts = 1. The selected radius allows grouping of residues forming spatially contiguous allosteric regions while remaining permissive to structurally sparse annotations frequently observed in experimentally characterized allosteric sites. Setting minPts = 1 ensures that isolated annotated residues are retained as valid singleton clusters rather than discarded as noise, accommodating sparse residue-level annotations frequently encountered in experimentally characterized allosteric regions.

When multiple disconnected clusters were identified within the same protein structure, the largest cluster was retained as the representative allosteric region. The pocket center $\mathbf{c}$was subsequently defined as the centroid of the cluster coordinates:

$$\mathbf{c}=\frac{1}{N}\sum_{i=1}^{N} \mathbf{x}_{i}$$

where $N$denotes the number of residues in the selected cluster. Residues were then ranked according to the Euclidean distance between their centroids and the reconstructed pocket center, and the $K=100$nearest residues were selected to define the final fixed-size pocket representation following the same procedure described for the PPI-Site dataset.

**Text S3. Downstream Allostery Classification Framework**

**Text S3.1 Hyperparameter Optimization of Downstream Classifiers**

For downstream allostery classification, embeddings were first extracted from the frozen OneProt backbone using the selected combinations of pocket, sequence, and text embeddings. These embeddings were subsequently used to train Multi-Layer Perceptron (MLP) classifiers for discrimination between allosteric and orthosteric competitive binding sites.

To optimize classifier performance, we performed a systematic hyperparameter sweep over both architectural and training parameters. The evaluated search space comprised learning rates (0.001, 0.01), batch sizes (32, 64), maximum epochs (50), hidden-layer configurations ([256], [512, 256]), dropout rates (0.10, 0.25), layer normalization (enabled, disabled), activation functions (ReLU, GELU), and residual connections (enabled, disabled).

For each dataset, encoder architecture, embedding combination, and random seed, all hyperparameter combinations were evaluated. Model selection was performed using validation-set ROC-AUC, and the best-performing configuration was subsequently used for test-set evaluation. Early stopping was employed to reduce overfitting, with training terminated when validation performance ceased to improve.

This strategy enabled efficient exploration of a diverse range of classifier architectures while maintaining the OneProt backbone frozen. Consequently, downstream performance primarily reflects information encoded within the pretrained multimodal representations rather than extensive task-specific adaptation of the foundation model. The use of precomputed embeddings substantially reduced computational cost relative to end-to-end fine-tuning, allowing systematic evaluation across multiple datasets, encoder configurations, embedding combinations, and random seeds.

**Text S3.2 Performance Metrics**

Performance was evaluated using the following metrics.

ROC-AUC (Area Under the Receiver Operating Characteristic Curve) measures the model's ability to rank positive instances higher than negative instances, integrated over all possible classification thresholds:

$$\text{ROC-AUC}=\int_{0}^{1} \text{TPR}(t)\text{ }d\text{FPR}(t)$$

where $\text{TPR}(t)=\text{TP}(t)/(\text{TP}(t)+\text{FN}(t))$is the true positive rate at threshold $t$, and $\text{FPR}(t)=\text{FP}(t)/(\text{FP}(t)+\text{TN}(t))$is the false positive rate.

PR-AUC (Area Under the Precision-Recall Curve) is particularly informative for imbalanced datasets, measuring the trade-off between precision $\text{Precision}(t)=\text{TP}(t)/(\text{TP}(t)+\text{FP}(t))$and recall $\text{Recall}(t)=\text{TP}(t)/(\text{TP}(t)+\text{FN}(t))$:

$$\text{PR-AUC}=\int_{0}^{1} \text{Precision}(r)\text{ }dr$$

TPR (True Positive Rate) and TNR (True Negative Rate) are class-specific accuracy measures:

$$\text{TPR}=\text{TP}/(\text{TP}+\text{FN})$$

which is the proportion of allosteric sites correctly identified, and

$$\text{TNR}=\text{TN}/(\text{TN}+\text{FP})$$

which is the proportion of orthosteric sites correctly identified.

In addition to conventional ranking metrics, class-specific rates were analyzed to better characterize model behavior under strongly imbalanced allostery classification regimes.

**Text S3.3 Class Imbalance**

For the AlloDiverse dataset, positive samples substantially outnumbered negative samples ($N_{\text{allo}}=15,904$, $N_{\text{ortho}}=5,203$). To mitigate training bias introduced by class imbalance, positive instances were randomly subsampled during each training iteration to maintain balanced mini-batches while leaving validation and test distributions unchanged. Specifically, for each mini-batch, we sampled $B/2$positive and $B/2$negative instances uniformly at random from the training set, where $B$is the batch size. This balanced sampling strategy ensured that the classifier received equal exposure to both classes during training, preventing the model from simply learning the prior class distribution.

For the KinSite dataset with its severe class imbalance ($\rho\approx51.1$), this balanced sampling strategy was particularly important to ensure that the rare allosteric examples were adequately represented during training. The validation and test sets were left imbalanced to reflect real-world usage scenarios where allosteric sites are rare.

**Text S4. Implementation Details**

All experiments were implemented using Python 3.12 with PyTorch 2.7.0 as the deep learning framework. The OneProt multimodal backbone was obtained from the official repository and used with frozen pretrained weights. All MLP classifiers were implemented using PyTorch interface.

Computational resources consisted of NVIDIA A100 GPUs with 40GB memory for model inference and training. The total computational cost for all experiments, including the ten independent runs per configuration, was approximately 2,000 GPU-hours.

Code, dataset splits, and pretrained model weights will be made available upon publication to ensure reproducibility of all results.

**Supporting Figures**

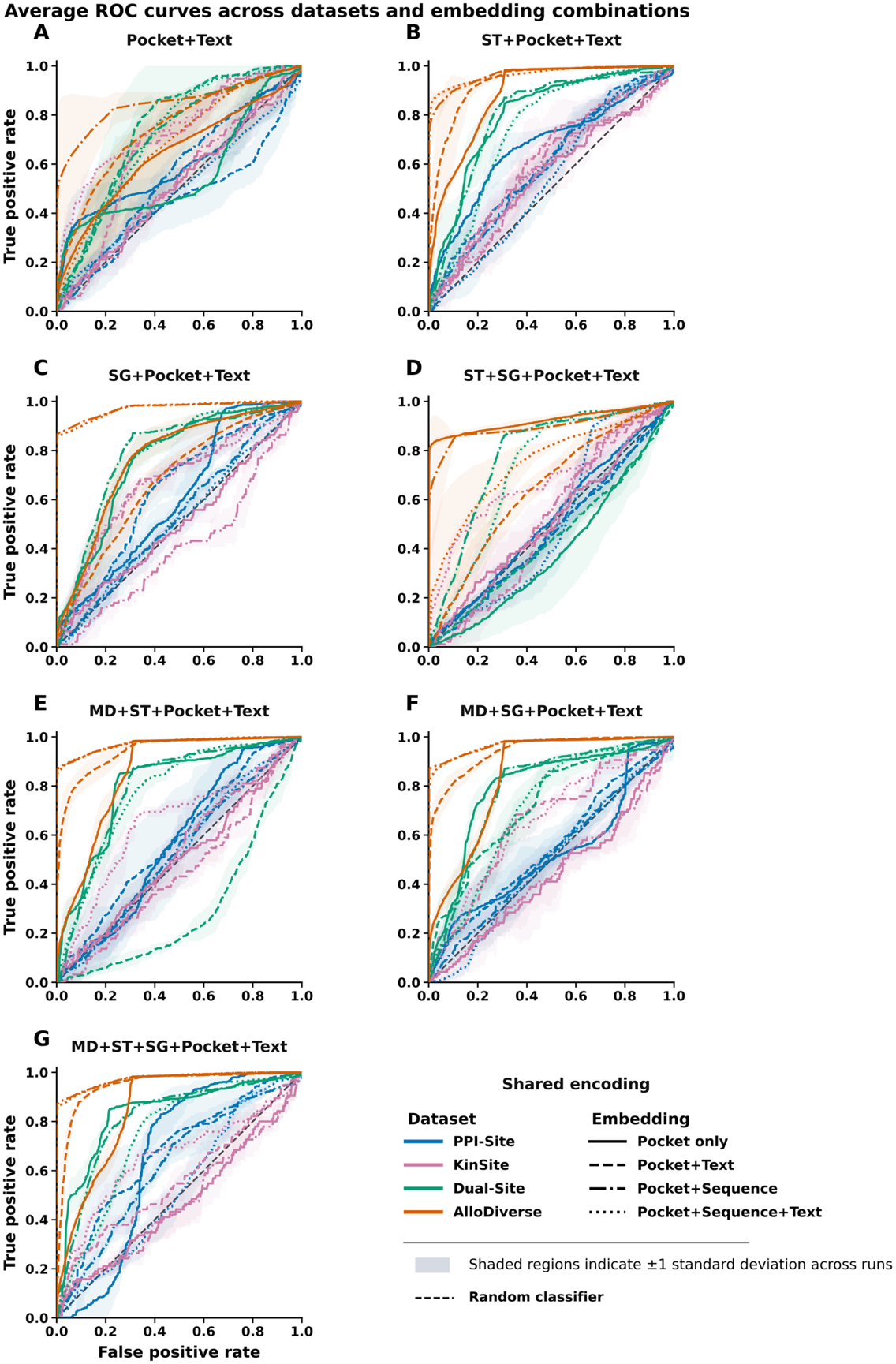

**Figure S1. Dataset-dependent ROC-AUC performance across OneProt encoder architectures and embedding combinations.** Receiver operating characteristic (ROC) curves illustrating the performance of seven OneProt encoder architectures (individual panels) evaluated across four allosteric discrimination datasets representing distinct separability regimes. Colors indicate datasets: PPI-Site (blue), KinSite (purple), Dual-Site (green), and AlloDiverse (red). Line styles denote the embedding combinations used for downstream classification: Pocket+Text (solid), Pocket+Sequence (dashed), and Pocket+Sequence+Text (dotted). Curves represent averages across independent runs, and shaded regions indicate ±1 standard deviation. Across all architectures, a consistent progression from low- to high-separability regimes is observed. PPI-Site remains close to random performance regardless of architecture or embedding composition. KinSite exhibits moderate separability and substantial variability. Dual-Site shows markedly improved discrimination, particularly for architectures incorporating structural and molecular-dynamics information. AlloDiverse consistently achieves near-ceiling ROC-AUC values across architectures. These results confirm that dataset composition exerts a substantially greater influence on allosteric discrimination performance than architectural variation alone.

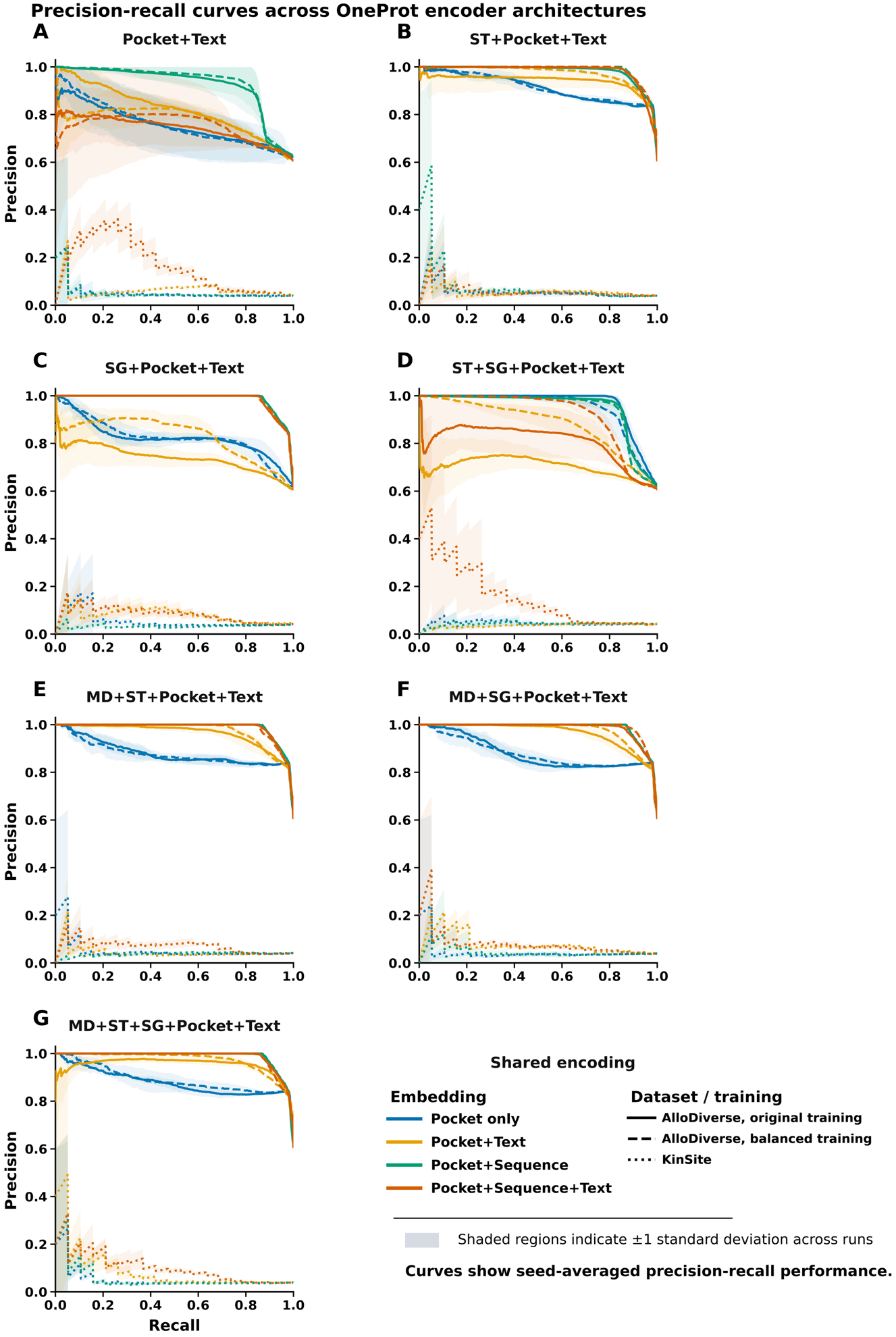
 **Figure S5. Precision–recall performance across OneProt encoder architectures under different class-imbalance regimes.** Precision–recall (PR) curves for seven OneProt encoder architectures (individual panels) evaluated using three embedding combinations: Pocket+Text (blue), Pocket+Sequence (green), and Pocket+Sequence+Text (red). The x-axis within each panel corresponds to the seven OneProt encoder architectures: Pocket+Text (baseline), ST+Pocket+Text (structure-token), SG+Pocket+Text (structure-graph), MD+ST+Pocket+Text (MD + structure-token), MD+SG+Pocket+Text (MD + structure-graph), ST+SG+Pocket+Text (both structural encoders), and MD+ST+SG+Pocket+Text (full multimodal). Line style denotes dataset or training condition: AlloDiverse original training (solid lines), AlloDiverse balanced training (dashed lines), and KinSite (dotted lines). Curves represent averages across independent runs, and shaded regions indicate ±1 standard deviation. The figure highlights how model behavior changes under different class-imbalance and separability conditions. In the highly separable AlloDiverse regime, most architectures achieve near-perfect precision across a broad range of recall values, particularly when sequence-containing embeddings are included. In contrast, the strongly imbalanced KinSite dataset exhibits substantially lower precision–recall performance, with pronounced sensitivity to embedding composition and architecture. Sequence-containing embeddings (green and red) generally maintain higher precision at increasing recall levels, whereas Pocket+Text (blue) shows more rapid performance degradation. Architectures incorporating structural and molecular-dynamics modalities frequently preserve higher precision over a wider recall range, suggesting that structural and dynamical information contributes complementary signal for recovering rare allosteric sites. These results underscore the importance of evaluating allostery prediction using metrics sensitive to positive-class recovery and further support the existence of distinct dataset-dependent learnability regimes

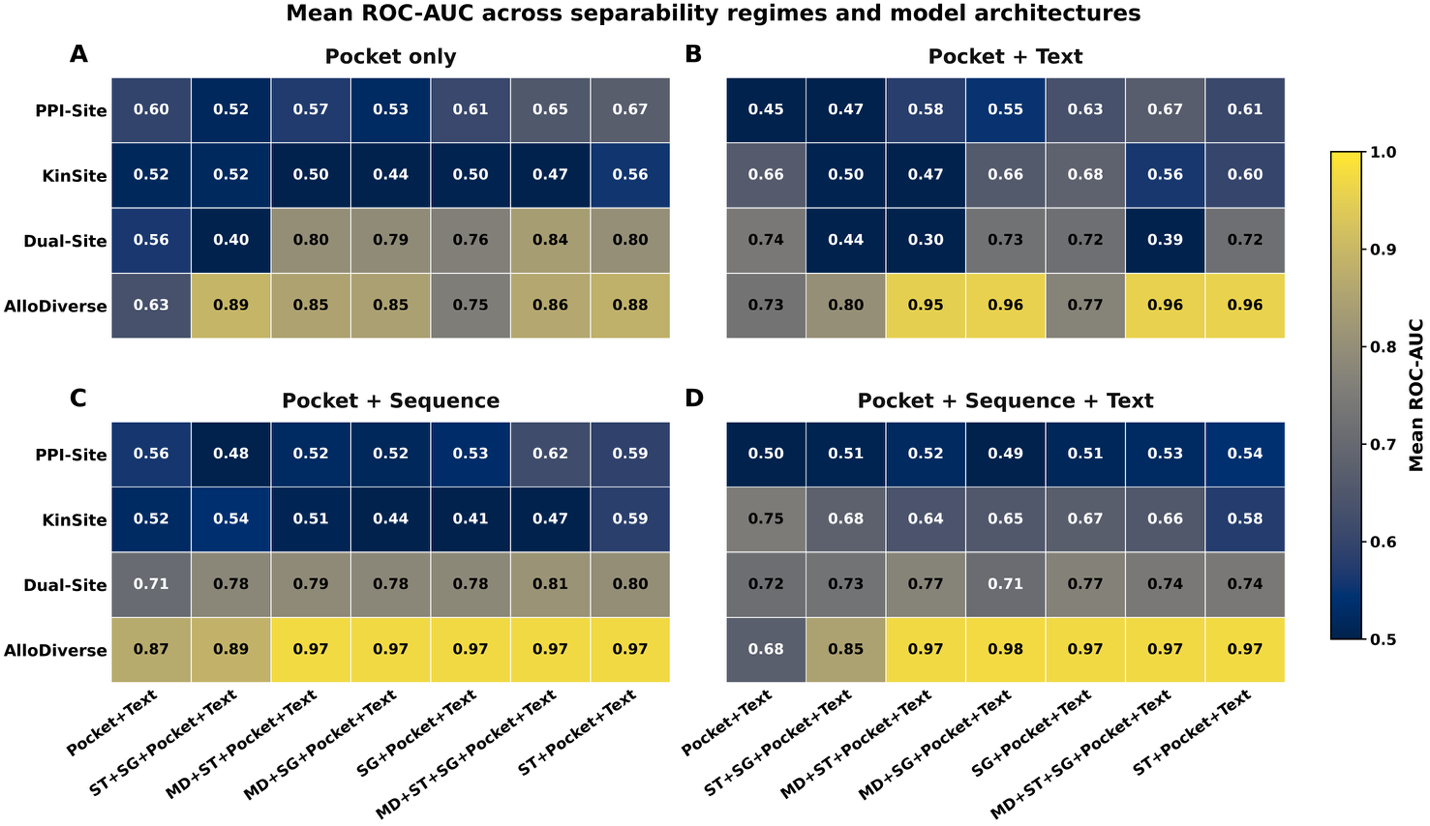

**Figure S3. Mean ROC-AUC across separability regimes, embedding combinations, and OneProt encoder architectures.** Heatmaps summarize the mean ROC-AUC values averaged across independent runs for all combinations of dataset regime (rows) and encoder architecture (columns). Panels correspond to the four downstream embedding combinations: (A) Pocket only, (B) Pocket+Text, (C) Pocket+Sequence, and (D) Pocket+Sequence+Text. Rows represent the four datasets arranged according to increasing separability: PPI-Site, KinSite, Dual-Site, and AlloDiverse. Columns correspond to the seven evaluated OneProt encoder architectures. Cell colors indicate mean ROC-AUC, with numerical values shown in each cell. Performance is primarily determined by dataset regime rather than architecture, with a clear progression from low performance in PPI-Site to near-ceiling performance in AlloDiverse.

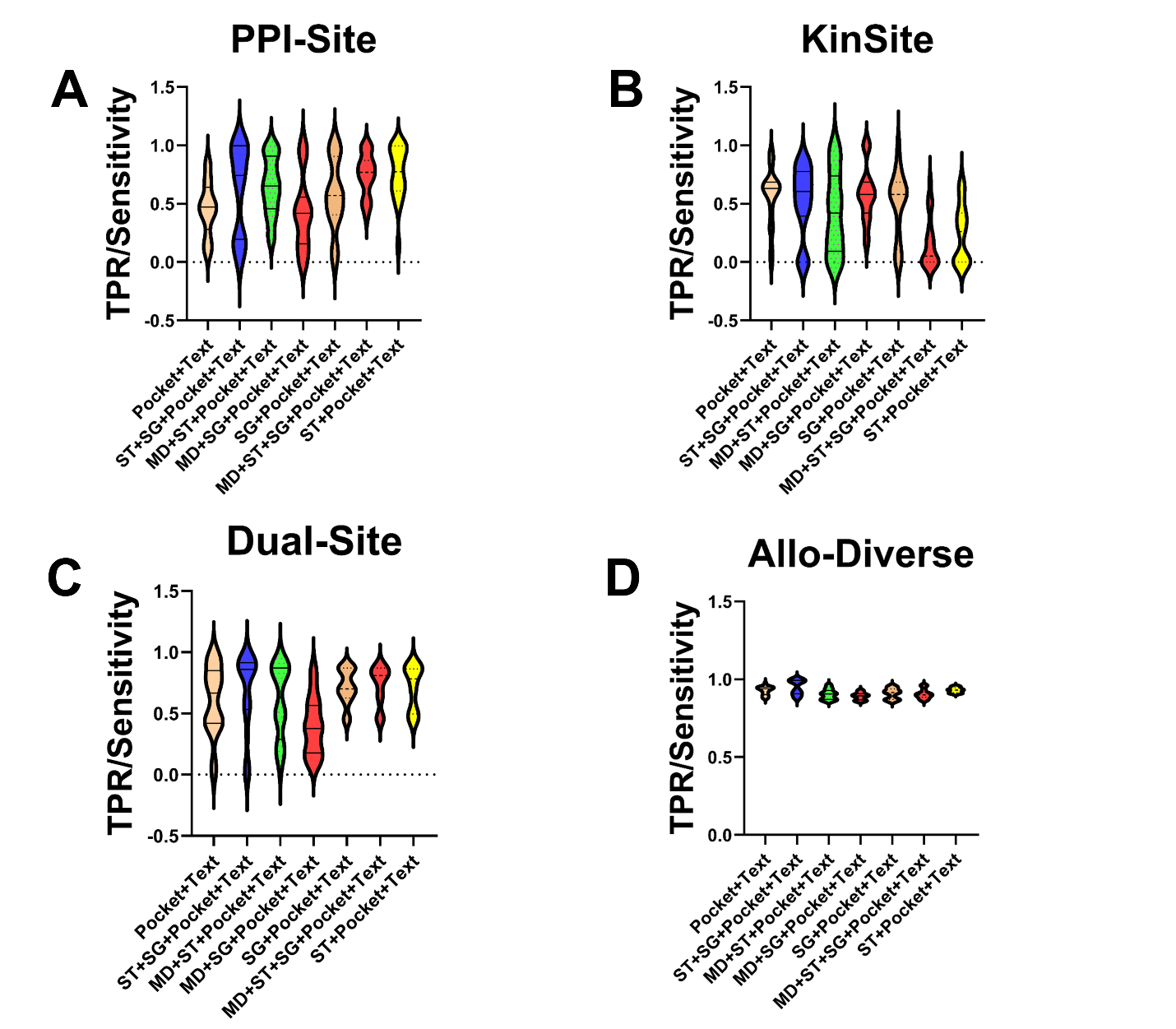

**Figure S4. TPR/Sensitivity (true positive rate) across allosteric binding site discrimination regimes.** Violin plots show the distributions of sensitivity (true positive rate for allosteric sites) across the four datasets representing distinct separability regimes: PPI-Site (A), KinSite (B), Dual-Site (C) , and AlloDiverse (D). The x-axis within each panel corresponds to the seven OneProt encoder architectures evaluated in this study: Pocket+Text (baseline), ST+Pocket+Text (structure-token), SG+Pocket+Text (structure-graph), MD+ST+Pocket+Text (MD + structure-token), MD+SG+Pocket+Text (MD + structure-graph), ST+SG+Pocket+Text (both structural encoders), and MD+ST+SG+Pocket+Text (full multimodal). Colored points within each violin indicate the embedding combination used for downstream classification. Boxes indicate the median and interquartile range across random seeds, while individual points represent single runs. The dashed horizontal line denotes random-expectation level (0.5). The figure reveals a progressive transition from low-separability (PPI-Site) to high-separability (AlloDiverse) regimes. In PPI-Site, sensitivity exhibits substantial variability and remains close to random performance across all architectures and embedding combinations, indicating limited discriminative signal. KinSite displays uniformly low sensitivity across configurations, reflecting the extreme difficulty of recovering allosteric sites under severe class imbalance (ρ ≈ 51.1). Dual-Site shows improved sensitivity, with sequence-containing embeddings (green and red) consistently outperforming pocket-only and pocket+text combinations across most architectures. In AlloDiverse, sensitivity approaches ceiling performance across all architectures and embedding combinations, indicating that most models readily identify allosteric sites.

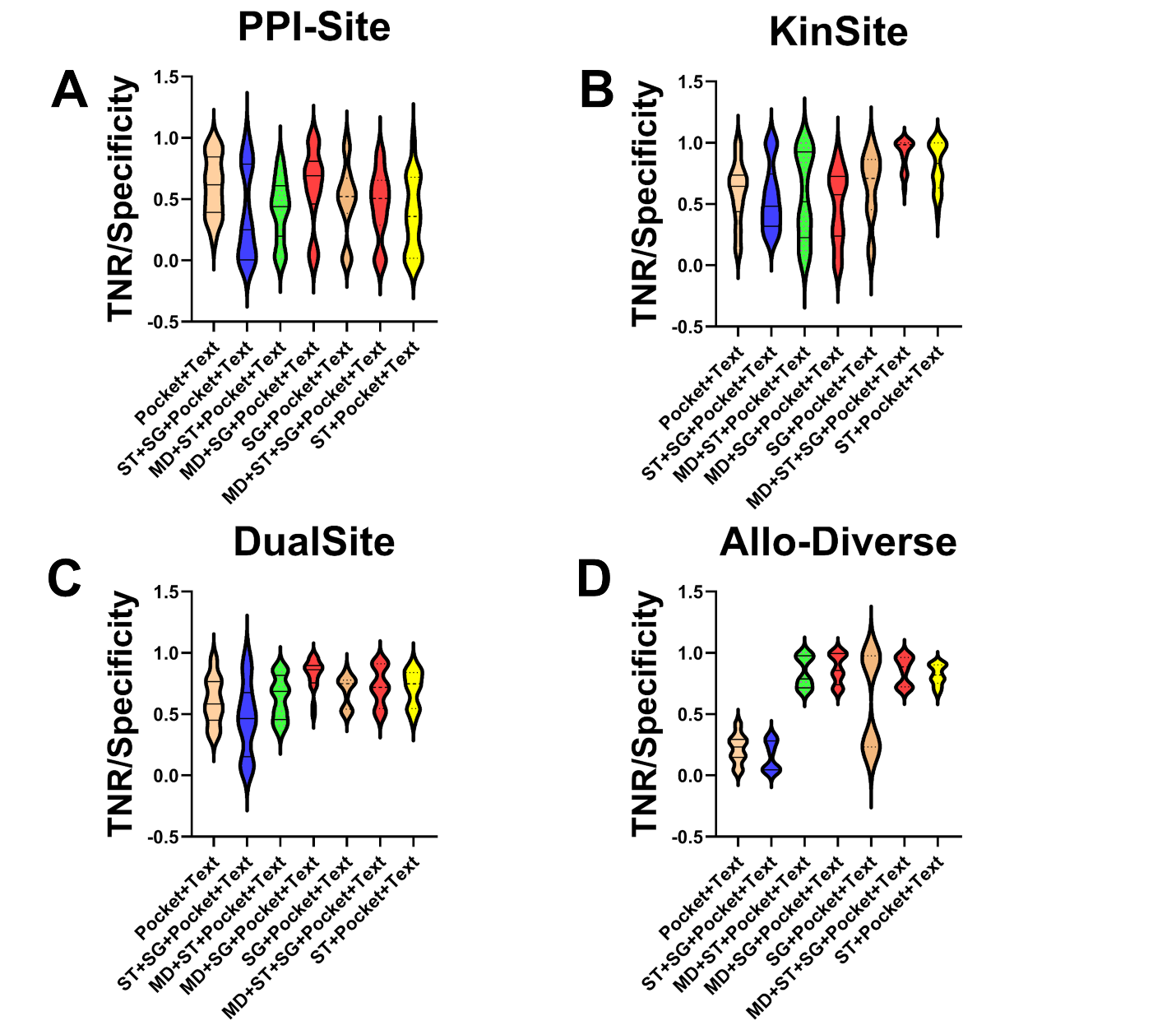

**Figure S5. TNR/Specificity (true negative rate) across allosteric binding site discrimination regimes.** Violin plots show the distributions of specificity (true negative rate for orthosteric competitive sites) across the four datasets representing distinct separability regimes: PPI-Site (A), KinSite (B), Dual-Site (C), and AlloDiverse (D). The x-axis within each panel corresponds to the seven OneProt encoder architectures evaluated in this study: Pocket+Text (baseline), ST+Pocket+Text (structure-token), SG+Pocket+Text (structure-graph), MD+ST+Pocket+Text (MD + structure-token), MD+SG+Pocket+Text (MD + structure-graph), ST+SG+Pocket+Text (both structural encoders), and MD+ST+SG+Pocket+Text (full multimodal). Colored points within each violin indicate the embedding combination used for downstream classification. Boxes indicate the median and interquartile range across random seeds, while individual points represent single runs. The dashed horizontal line denotes random-expectation level (0.5). The figure reveals a progressive transition from low-separability (PPI-Site) to high-separability (AlloDiverse) regimes. In PPI-Site, specificity exhibits substantial variability and remains close to random performance across all architectures and embedding combinations, indicating limited discriminative signal. KinSite displays higher specificity compared to sensitivity (Figure S2), reflecting the relative ease of identifying orthosteric ATP-binding sites under extreme class imbalance (ρ ≈ 51.1). Dual-Site shows improved specificity, with sequence-containing embeddings (green and red) consistently outperforming Pocket+Text across most architectures. In AlloDiverse, specificity remains more variable than sensitivity, indicating that models readily identify allosteric sites but differ in their ability to reject orthosteric sites.
