## Supporting Text S1-S4, Supporting Figures S1-S5 and Supporting Tables S1,S2. for "Integrated Framework for Probing Multimodal Protein Foundation Models with Structure-Functional Interpretability Analysis in Detection of Allosteric Binding Sites": SUPPORTING_INFORMATION_BIORXIV.pdf

### Text S1. Pre-training Architectures

#### Text S1.1 Structural Encoders

Structural representations were generated using two complementary structural encoders integrated within OneProt: a graph-based structural encoder and a token-based structural encoder. These encoders capture complementary aspects of protein three-dimensional organization across multiple spatial scales.

Graph-based structural representations were obtained using the ProNet architecture, a hierarchical graph neural network that learns protein representations at three complementary levels of structural granularity: amino-acid, backbone, and all-atom. Given a protein structure containing $L$amino-acid residues, the protein is represented as a graph $G = (V, E, P)$, where $V$denotes the set of residue nodes, $E$represents the edges describing covalent and spatial relationships between residues, and $P$contains the structural information associated with the protein. Through hierarchical modeling at the amino-acid, backbone, and all-atom levels, ProNet captures complementary structural features spanning local geometric environments and higher-order protein organization.

At each level, node representations are iteratively refined using message passing according to the general update rule:

$$m_v^{(t)} = \text{AGGREGATE}_{u \in \mathcal{N}(v)} \left( \text{MESSAGE}^{(t)} (h_v^{(t)}, h_u^{(t)}, e_{uv}) \right)$$

$$h_v^{(t+1)} = \text{UPDATE}^{(t)} (h_v^{(t)}, m_v^{(t)})$$

where $\mathcal{N}(v)$denotes the neighborhood of node $v$, $e_{uv}$represents geometric edge features derived from the structural information contained in $P$, and MESSAGE, AGGREGATE, and UPDATE are learnable functions at layer $t$.

Through hierarchical message passing across the amino-acid, backbone, and all-atom representations, ProNet learns structural embeddings that capture fine-grained local interactions together with the higher-order three-dimensional organization of the complete protein fold. Within OneProt, these graph embeddings are projected into the shared multimodal latent space and aligned with sequence, pocket, text, and MD  representations during contrastive pretraining.

Token-based structural representations were generated using Foldseek structural tokens processed with an ESM2-35M transformer encoder. Foldseek converts protein structures into sequence-like representations by discretizing local three-dimensional residue environments into a learned structural alphabet. Each structural token encodes the geometric relationship between a residue and its nearest spatial neighbor, capturing local tertiary structural information in a discrete representation.

The resulting structural token sequence:

$$S_{struct} = (t_1, t_2, \dots, t_L)$$

where each token $t_i \in \mathcal{T}$belongs to a learned structural alphabet of size $| \mathcal{T} | = 20$, is processed by an ESM2 transformer encoder to produce contextual structural embeddings:

For a batch of $N$protein samples, let $\mathbf{z}_{\text{seq}}^{(i)}$, $\mathbf{z}_{\text{pocket}}^{(i)}$, $\mathbf{z}_{\text{text}}^{(i)}$, $\mathbf{z}_{\text{struct}}^{(i)}$, and $\mathbf{z}_{\text{MD}}^{(i)}$denote the normalized embeddings for sample $i$from the sequence, pocket, text, structural, and MD modalities, respectively.

The contrastive loss for aligning sequence and pocket modalities is defined as:

$$\mathcal{L}_{\text{seq-pocket}} = -\frac{1}{N}\sum_{i=1}^{N} \log \frac{\exp\,(\mathbf{z}_{\text{seq}}^{(i)} \cdot \mathbf{z}_{\text{pocket}}^{(i)}/\tau)}{\sum_{j=1}^{N} \exp\,(\,\mathbf{z}_{\text{seq}}^{(i)} \cdot \mathbf{z}_{\text{pocket}}^{(j)}/\tau)}$$

where $\tau > 0$is a temperature parameter controlling the sharpness of the softmax distribution, and the dot product $\mathbf{z}_{\text{seq}}^{(i)} \cdot \mathbf{z}_{\text{pocket}}^{(j)}$measures the cosine similarity between normalized embeddings.

The total multimodal alignment loss is the sum over all modality pairs anchored to the sequence modality:

$$\mathcal{L}_{\text{total}} = \sum_{m \in \mathcal{M}} \mathcal{L}_{\text{seq-}m}$$

where $\mathcal{M} = \{\text{pocket,text,struct,MD}\}$.

$$\mathbf{c}_{\text{pocket}} = \frac{1}{N_{\text{lig}}} \sum_{i=1}^{N_{\text{lig}}} \mathbf{x}_i^{\text{lig}}$$

where $\left\{ \mathbf{x}_i^{\text{lig}} \right\}_{i=1}^{N_{\text{lig}}} \subset \mathbb{R}^3$ are the coordinates of ligand atoms.

For the associated protein chain, residue centroids were computed as the average coordinates of all atoms belonging to each standard amino-acid residue:

$$\mathbf{c}_r = \frac{1}{N_r} \sum_{a \in r} \mathbf{x}_a$$

where $N_r$ is the number of atoms in residue $r$.

Residues were subsequently ranked according to the Euclidean distance between their centroids and the ligand center:

$$d(r, \mathbf{c}_{\text{pocket}}) = \| \mathbf{c}_r - \mathbf{c}_{\text{pocket}} \|_2$$

and the $K = 100$nearest residues were retained to construct fixed-size pocket representations:

$$\mathcal{R}_{\text{pocket}} = \{r_{(1)}, r_{(2)}, \dots, r_{(K)}\}$$

where $r_{(1)}, r_{(2)}, \dots$denote residues sorted by increasing distance to the pocket center. All atoms belonging to the selected residues were preserved, yielding atom-level pocket structures together with their associated residue sequences. The complete protein-chain sequence was stored separately for sequence-based representation extraction.

**Text S2.2 Pocket Overlap and Conformational Variability Analysis**

Because the PPI-Site dataset contains structurally resolved pocket annotations spanning both allosteric and orthosteric competitive binding regimes within shared protein contexts, it enables direct comparison of local binding environments across regulatory classes. Equivalent analyses were not performed on the AlloDiverse dataset because ASD pockets are reconstructed from residue-level annotations rather than experimentally resolved cavity definitions, preventing direct structural comparison of matched pocket pairs. Similarly, the KinSite dataset predominantly reflects highly conserved ATP-binding environments under strong class imbalance, limiting the interpretability of residue-overlap analyses across regulatory states.

To assess the extent of structural overlap between allosteric and orthosteric competitive binding environments in PPI-Site, we compared pockets across both classes for proteins sharing the same protein and chain identifiers. Comparisons were performed at both the residue-composition level and the atom-coordinate level.

**Text S2.3 Residue-Level Overlap Analysis**

Residue-level overlap was quantified using the Jaccard similarity coefficient. Given residue sets $A$and $B$corresponding to two pockets, the Jaccard similarity was defined as:

$$J(A,B) = \frac{| A \cap B |}{| A \cup B |}$$

where $| A \cap B |$denotes the number of shared residues and $| A \cup B |$denotes the total number of unique residues across both pockets.

In addition to Jaccard similarity, we computed the overlap coefficient:

Given a set of residue coordinates $\{\mathbf{x}_i\}_{i=1}^{N} \subset \mathbb{R}^3$ , two residues $\mathbf{x}_i$and $\mathbf{x}_j$are considered neighbors if their Euclidean distance satisfies:

$$\| \mathbf{x}_i - \mathbf{x}_j \|_2 < \varepsilon$$

Code, dataset splits, and pretrained model weights will be made available upon publication to ensure reproducibility of all results.

### Supporting Figures

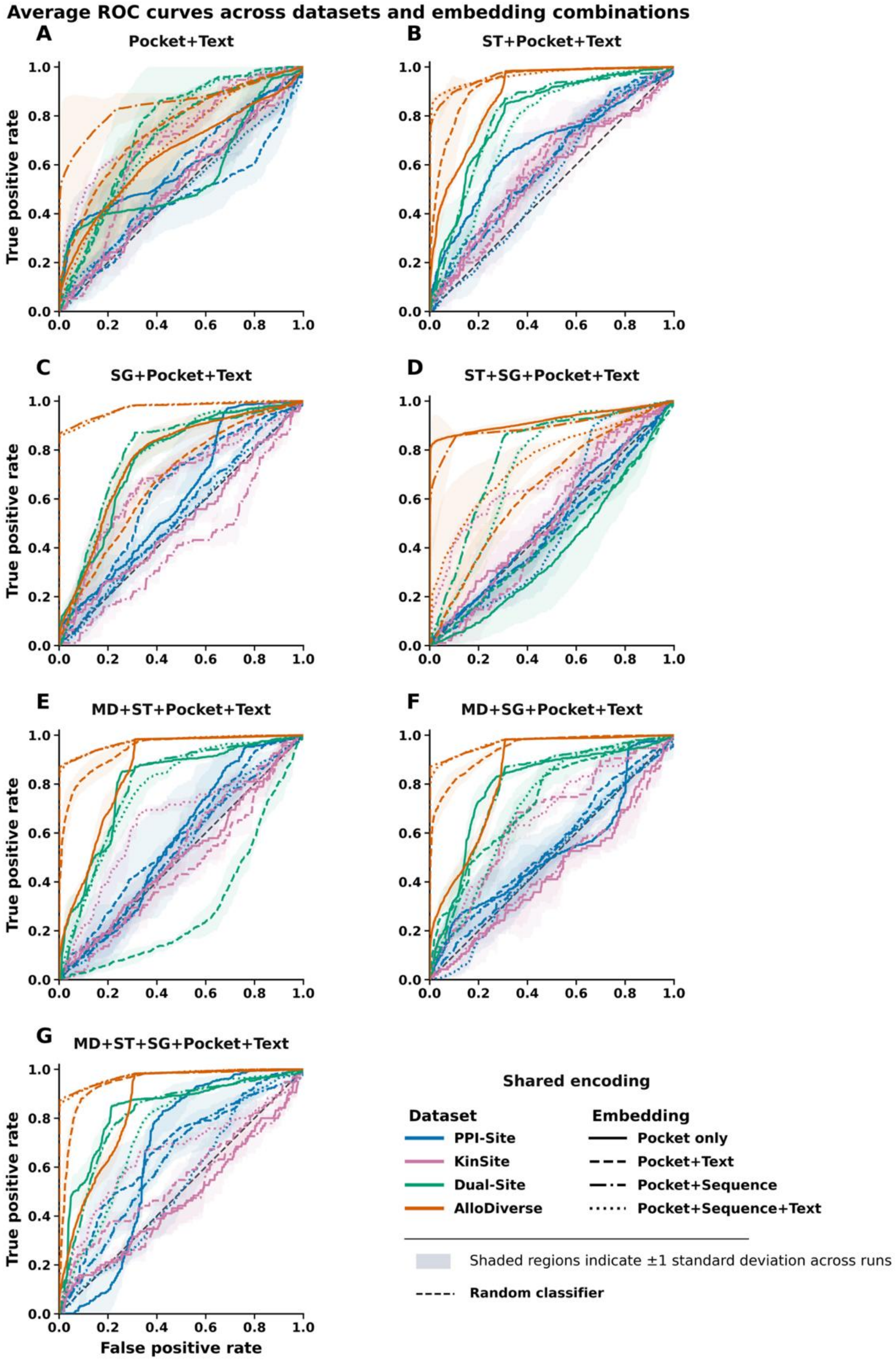

**Figure S1. Dataset-dependent ROC-AUC performance across OneProt encoder architectures and embedding combinations.** Receiver operating characteristic (ROC) curves illustrating the performance of seven OneProt encoder architectures (individual panels) evaluated across four allosteric discrimination datasets representing distinct separability regimes. Colors indicate datasets: PPI-Site (blue), KinSite (purple), Dual-Site (green), and AlloDiverse (red). Line styles denote the embedding combinations used for downstream classification: Pocket+Text (solid), Pocket+Sequence (dashed), and Pocket+Sequence+Text (dotted). Curves represent averages across independent runs, and shaded regions indicate ±1 standard deviation. Across all architectures, a consistent progression from low- to high-separability regimes is observed. PPI-Site remains close to random performance regardless of architecture or embedding composition. KinSite exhibits moderate separability and substantial variability. Dual-Site shows markedly improved discrimination, particularly for architectures incorporating structural and molecular-dynamics information. AlloDiverse consistently achieves near-ceiling ROC-AUC values across architectures. These results confirm that dataset composition exerts a substantially greater influence on allosteric discrimination performance than architectural variation alone.

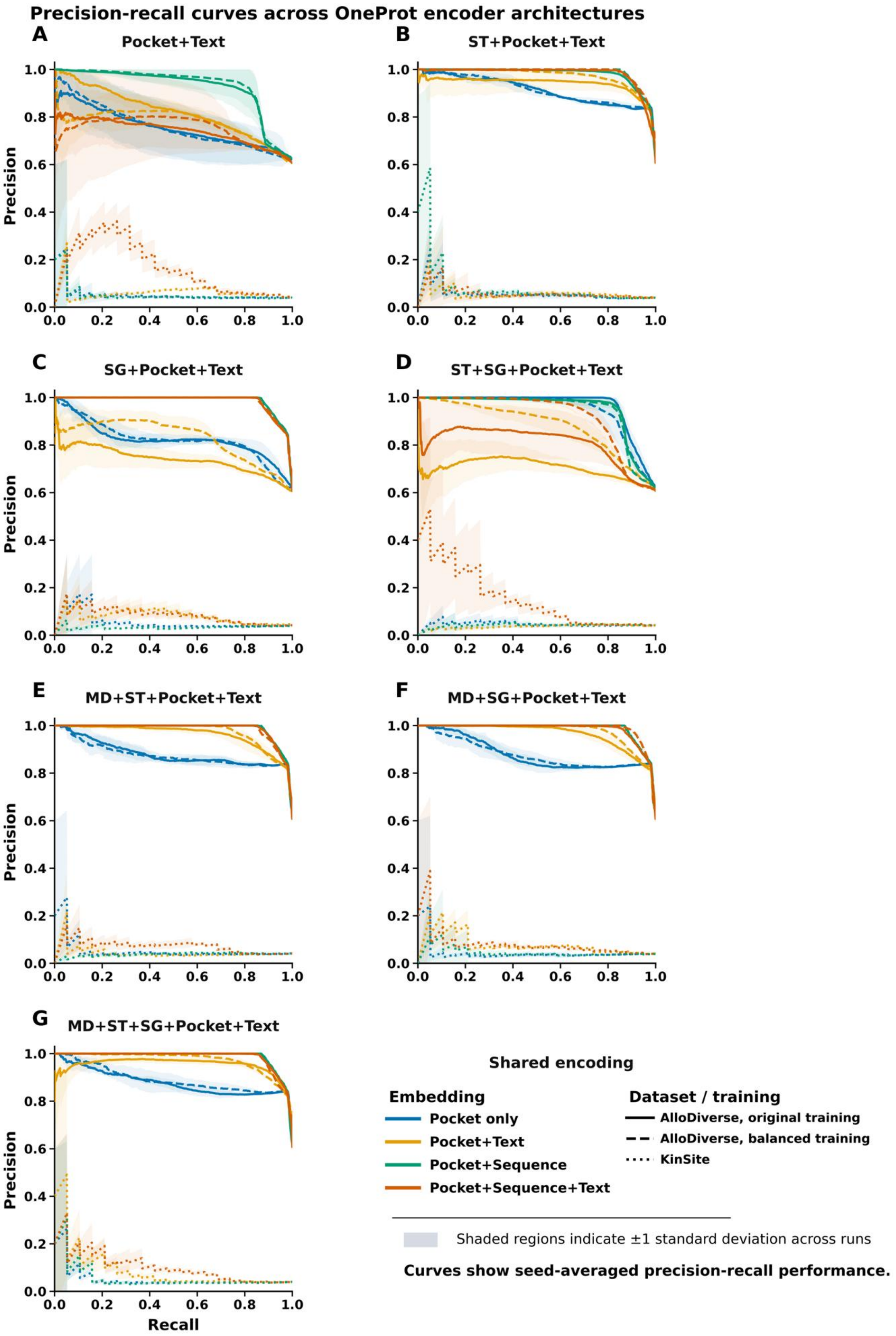

Precision-recall curves across OneProt encoder architectures
A
Pocket+Text
B
ST+Pocket+Text
C
SG+Pocket+Text
D
ST+SG+Pocket+Text
E
MD+ST+Pocket+Text
F
MD+SG+Pocket+Text
G
MD+ST+SG+Pocket+Text
Precision
Recall
Shared encoding
Embedding
Pocket only
Pocket+Text
Pocket+Sequence
Pocket+Sequence+Text
Dataset / training
AlloDiverse, original training
AlloDiverse, balanced training
KinSite
Shaded regions indicate ±1 standard deviation across runs
Curves show seed-averaged precision-recall performance.

**Figure S5. Precision–recall performance across OneProt encoder architectures under different class-imbalance regimes.** Precision–recall (PR) curves for seven OneProt encoder architectures (individual panels) evaluated using three embedding combinations: Pocket+Text (blue), Pocket+Sequence (green), and Pocket+Sequence+Text (red). The x-axis within each panel corresponds to the seven OneProt encoder architectures: Pocket+Text (baseline), ST+Pocket+Text (structure-token), SG+Pocket+Text (structure-graph), MD+ST+Pocket+Text (MD + structure-token), MD+SG+Pocket+Text (MD + structure-graph), ST+SG+Pocket+Text (both structural encoders), and MD+ST+SG+Pocket+Text (full multimodal). Line style denotes dataset or training condition: AlloDiverse original training (solid lines), AlloDiverse balanced training (dashed lines), and KinSite (dotted lines). Curves represent averages across independent runs, and shaded regions indicate ±1 standard deviation. The figure highlights how model behavior changes under different class-imbalance and separability conditions. In the highly separable AlloDiverse regime, most architectures achieve near-perfect precision across a broad range of recall values, particularly when sequence-containing embeddings are included. In contrast, the strongly imbalanced KinSite dataset exhibits substantially lower precision–recall performance, with pronounced sensitivity to embedding composition and architecture. Sequence-containing embeddings (green and red) generally maintain higher precision at increasing recall levels, whereas Pocket+Text (blue) shows more rapid performance degradation. Architectures incorporating structural and molecular-dynamics modalities frequently preserve higher precision over a wider recall range, suggesting that structural and dynamical information contributes complementary signal for recovering rare allosteric sites. These results underscore the importance of evaluating allostery prediction using metrics sensitive to positive-class recovery and further support the existence of distinct dataset-dependent learnability regimes

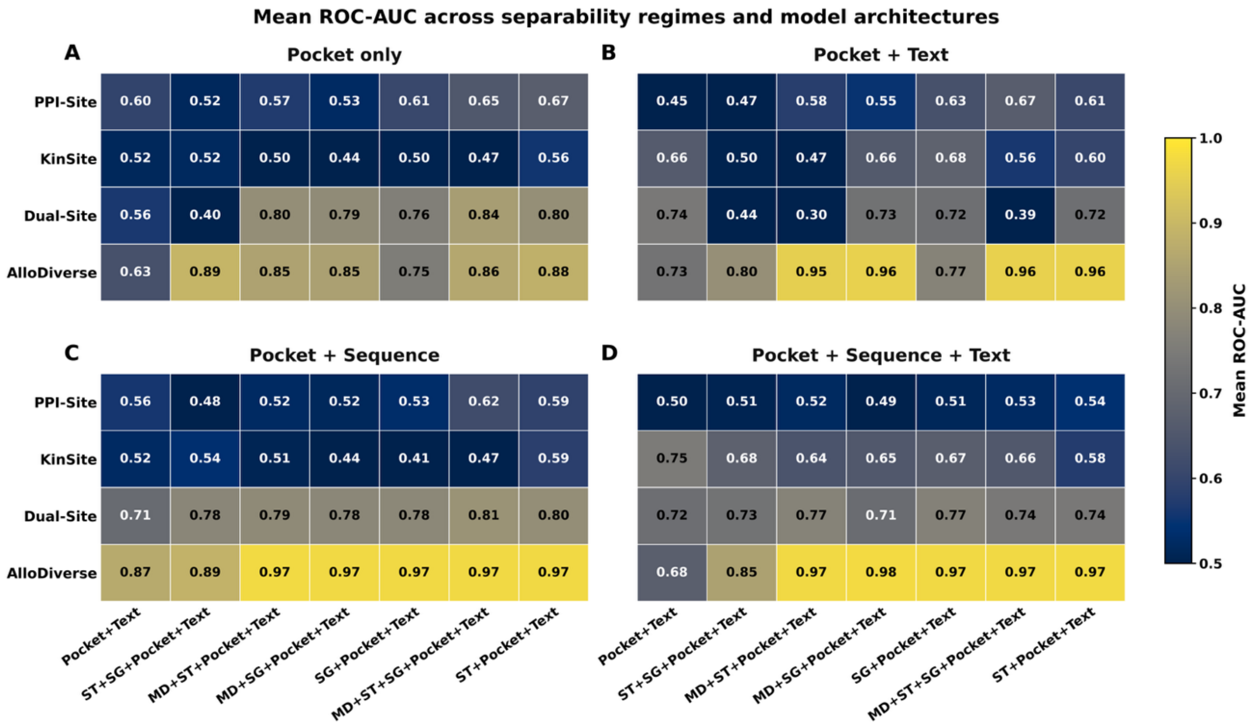


**Figure S3. Mean ROC-AUC across separability regimes, embedding combinations, and OneProt encoder architectures.** Heatmaps summarize the mean ROC-AUC values averaged across independent runs for all combinations of dataset regime (rows) and encoder architecture (columns). Panels correspond to the four downstream embedding combinations: (A) Pocket only, (B) Pocket+Text, (C) Pocket+Sequence, and (D) Pocket+Sequence+Text. Rows represent the four datasets arranged according to increasing separability: PPI-Site, KinSite, Dual-Site, and AlloDiverse. Columns correspond to the seven evaluated OneProt encoder architectures. Cell colors indicate mean ROC-AUC, with numerical values shown in each cell. Performance is primarily determined by dataset regime rather than architecture, with a clear progression from low performance in PPI-Site to near-ceiling performance in AlloDiverse.

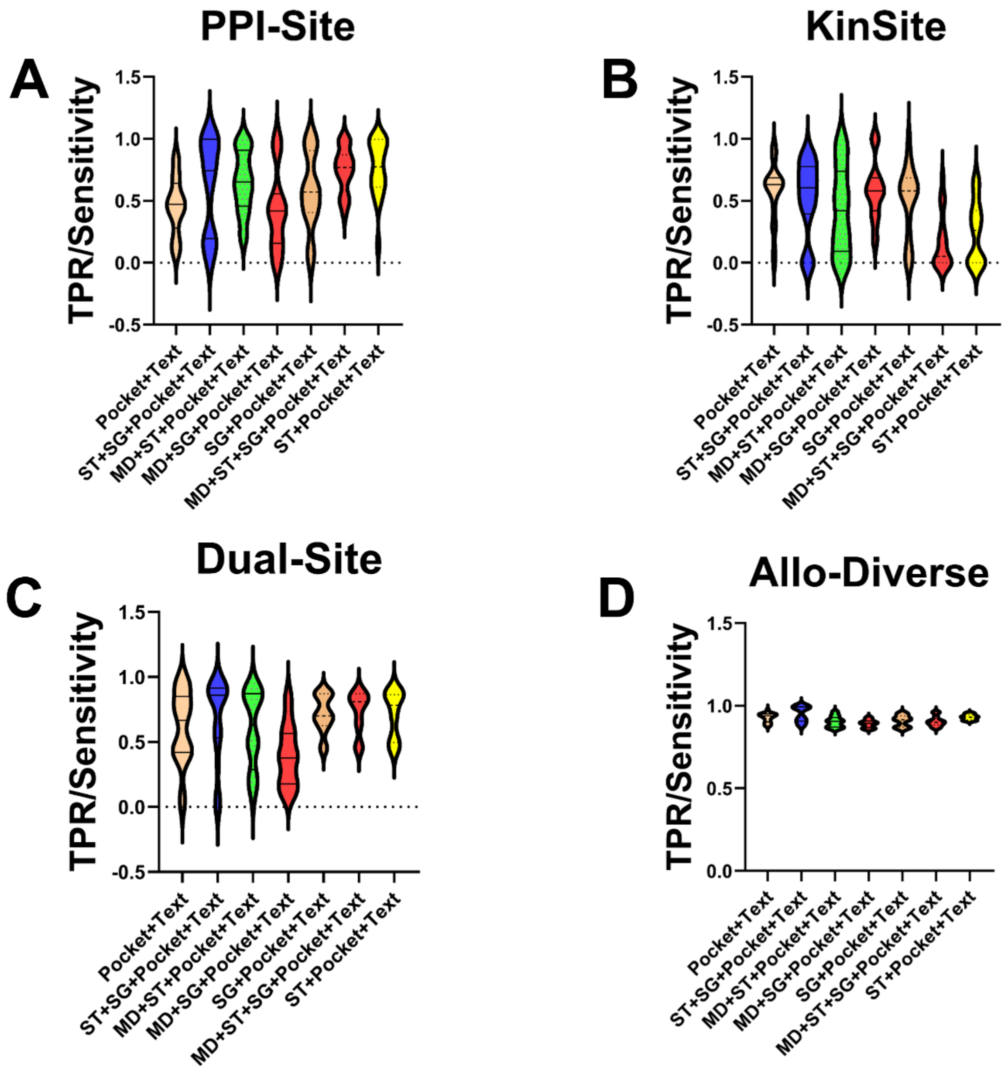


**Figure S4. TPR/Sensitivity (true positive rate) across allosteric binding site discrimination regimes.** Violin plots show the distributions of sensitivity (true positive rate for allosteric sites) across the four datasets representing distinct separability regimes: PPI-Site (A), KinSite (B), Dual-Site (C) , and AlloDiverse (D). The x-axis within each panel corresponds to the seven OneProt encoder architectures evaluated in this study: Pocket+Text (baseline), ST+Pocket+Text (structure-token), SG+Pocket+Text (structure-graph), MD+ST+Pocket+Text (MD + structure-token), MD+SG+Pocket+Text (MD + structure-graph), ST+SG+Pocket+Text (both structural encoders), and MD+ST+SG+Pocket+Text (full multimodal). Colored points within each violin indicate the embedding combination used for downstream classification. Boxes indicate the median and interquartile range across random seeds, while individual points represent single runs. The dashed horizontal line denotes random-expectation level (0.5). The figure reveals a progressive transition from low-separability (PPI-Site) to high-separability (AlloDiverse) regimes. In PPI-Site, sensitivity exhibits substantial variability and remains close to random performance across all architectures and embedding combinations, indicating limited discriminative signal. KinSite displays uniformly low sensitivity across configurations, reflecting the extreme difficulty of recovering allosteric sites under severe class imbalance ($\rho \approx 51.1$). Dual-Site shows improved sensitivity, with sequence-containing embeddings (green and red) consistently outperforming pocket-only and pocket+text combinations across most architectures. In AlloDiverse, sensitivity approaches ceiling performance across all architectures and embedding combinations, indicating that most models readily identify allosteric sites.

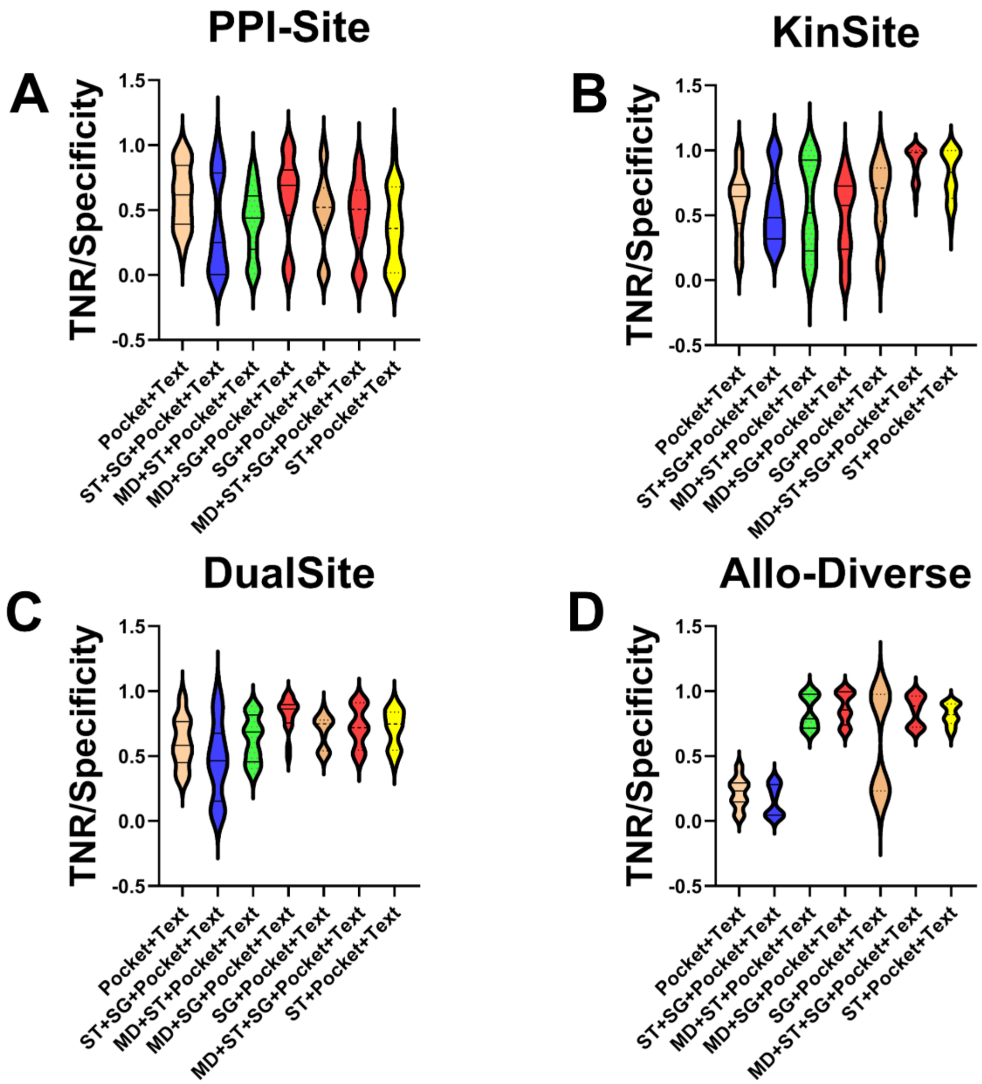


**Figure S5. TNR/Specificity (true negative rate) across allosteric binding site discrimination regimes.** Violin plots show the distributions of specificity (true negative rate for orthosteric competitive sites) across the four datasets representing distinct separability regimes: PPI-Site (A), KinSite (B), Dual-Site (C), and AlloDiverse (D). The x-axis within each panel corresponds to the seven OneProt encoder architectures evaluated in this study: Pocket+Text (baseline), ST+Pocket+Text (structure-token), SG+Pocket+Text (structure-graph), MD+ST+Pocket+Text

(MD + structure-token), MD+SG+Pocket+Text (MD + structure-graph), ST+SG+Pocket+Text (both structural encoders), and MD+ST+SG+Pocket+Text (full multimodal). Colored points within each violin indicate the embedding combination used for downstream classification. Boxes indicate the median and interquartile range across random seeds, while individual points represent single runs. The dashed horizontal line denotes random-expectation level (0.5). The figure reveals a progressive transition from low-separability (PPI-Site) to high-separability (AlloDiverse) regimes. In PPI-Site, specificity exhibits substantial variability and remains close to random performance across all architectures and embedding combinations, indicating limited discriminative signal. KinSite displays higher specificity compared to sensitivity (Figure S2), reflecting the relative ease of identifying orthosteric ATP-binding sites under extreme class imbalance ($\rho \approx 51.1$). Dual-Site shows improved specificity, with sequence-containing embeddings (green and red) consistently outperforming Pocket+Text across most architectures. In AlloDiverse, specificity remains more variable than sensitivity, indicating that models readily identify allosteric sites but differ in their ability to reject orthosteric sites.
