## Supplementary figures and images for "Integrated Framework for Probing Multimodal Protein Foundation Models with Structure-Functional Interpretability Analysis in Detection of Allosteric Binding Sites"

### FigureS1_JCIM_SUBMISSION.tif

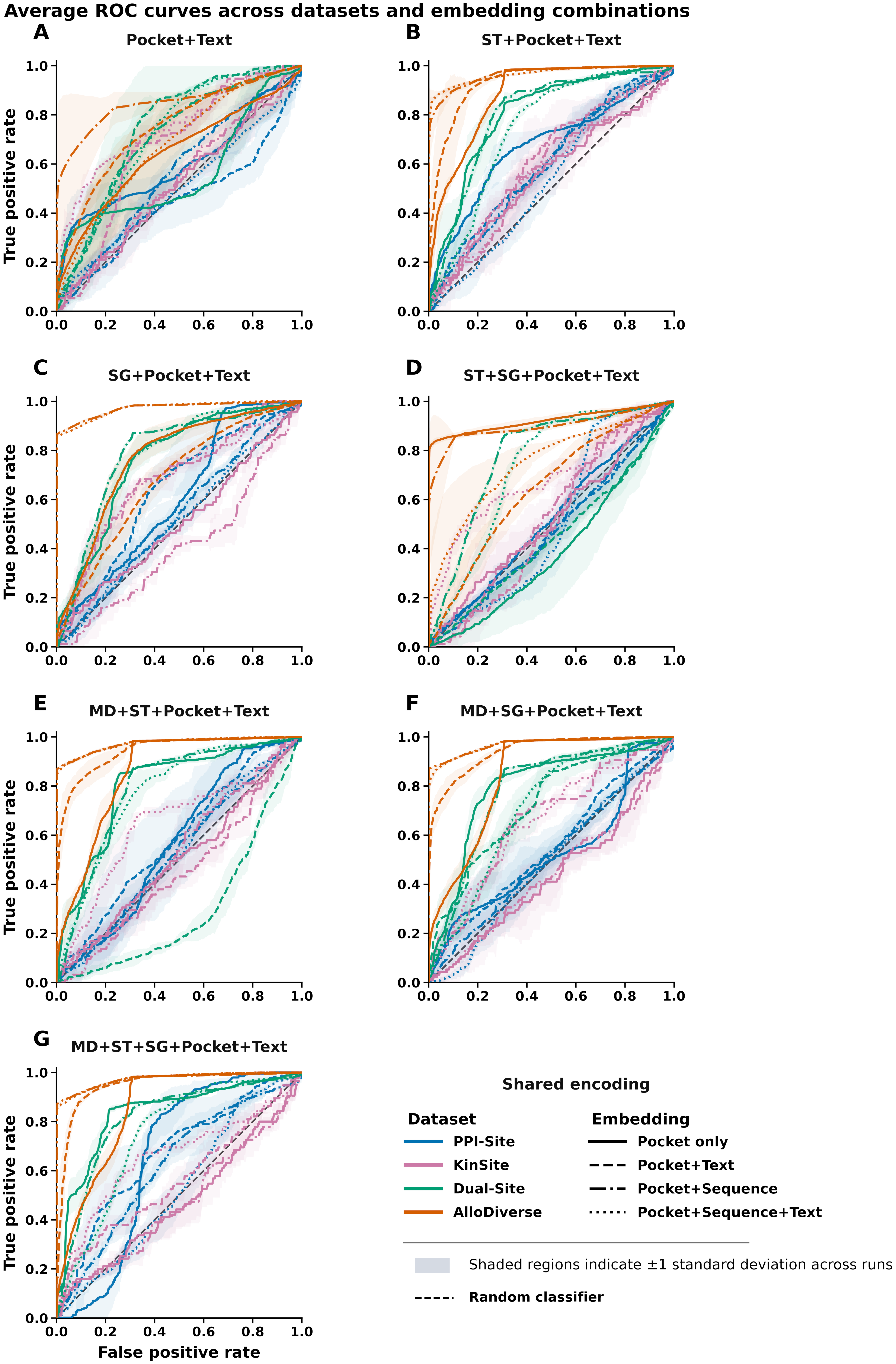

### FigureS2_JCIM_SUBMISSION.tif

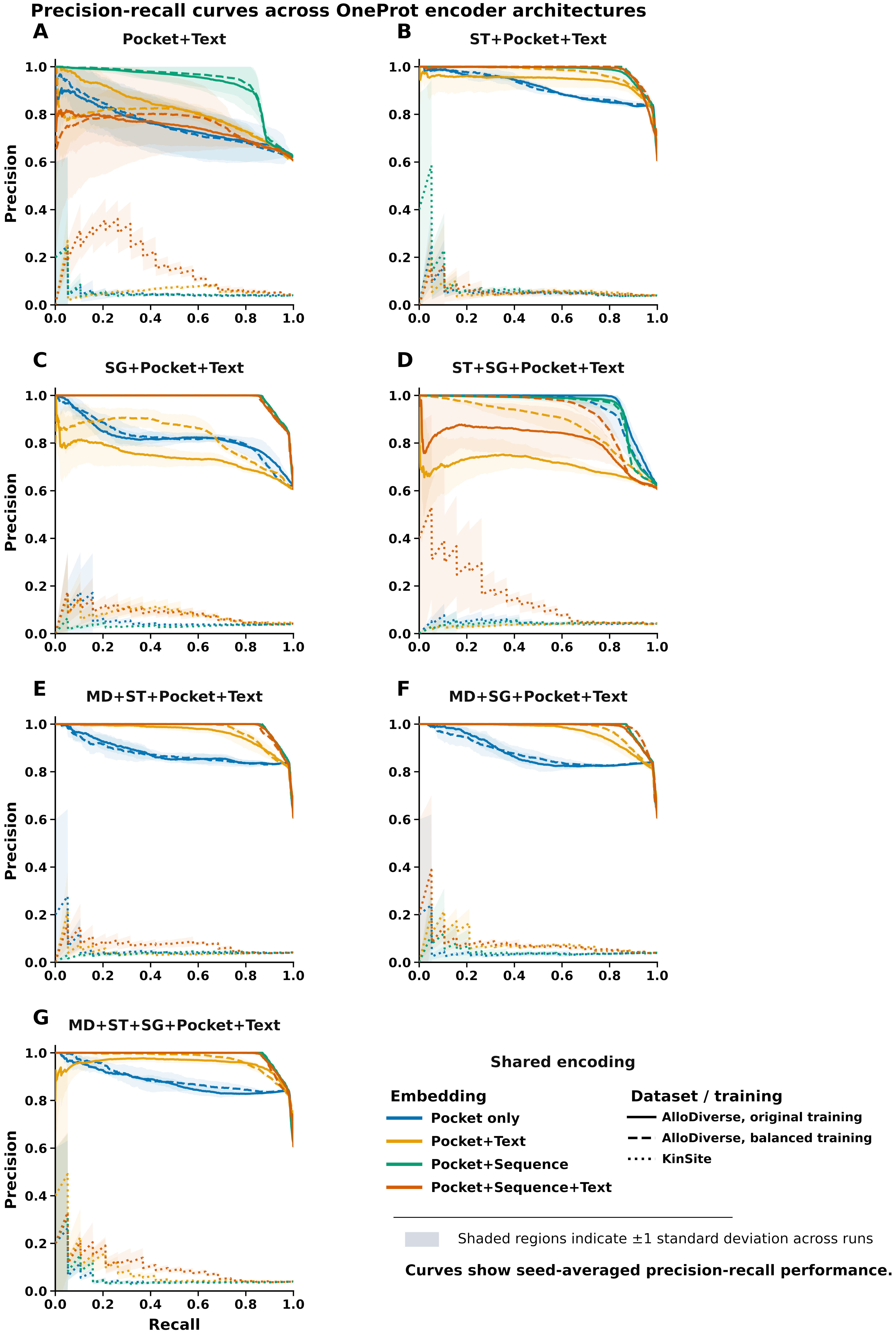

### FigureS3_JCIM_SUBMISSION.tif

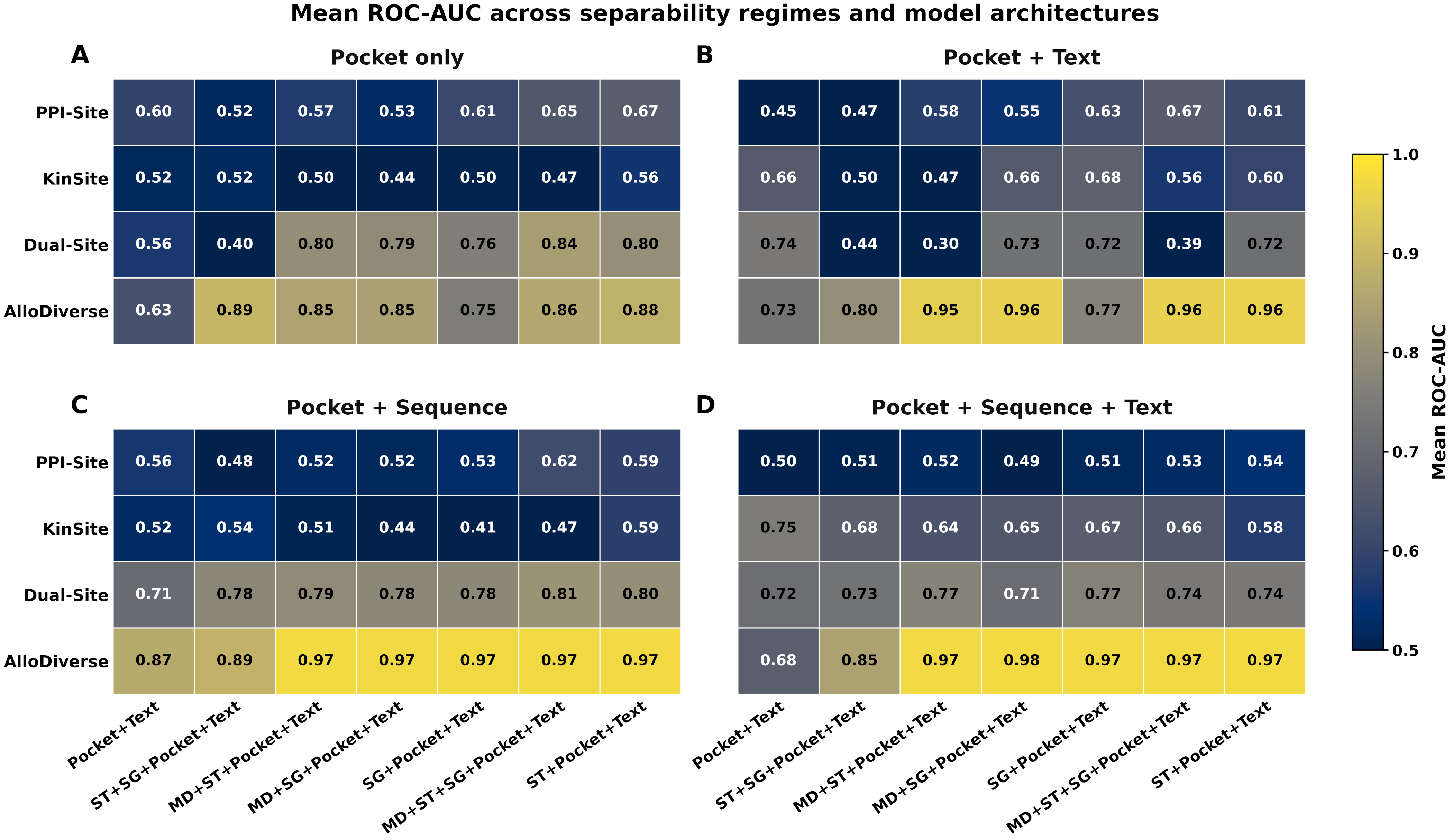

### FigureS4_JCIM_SUBMISSION.tif

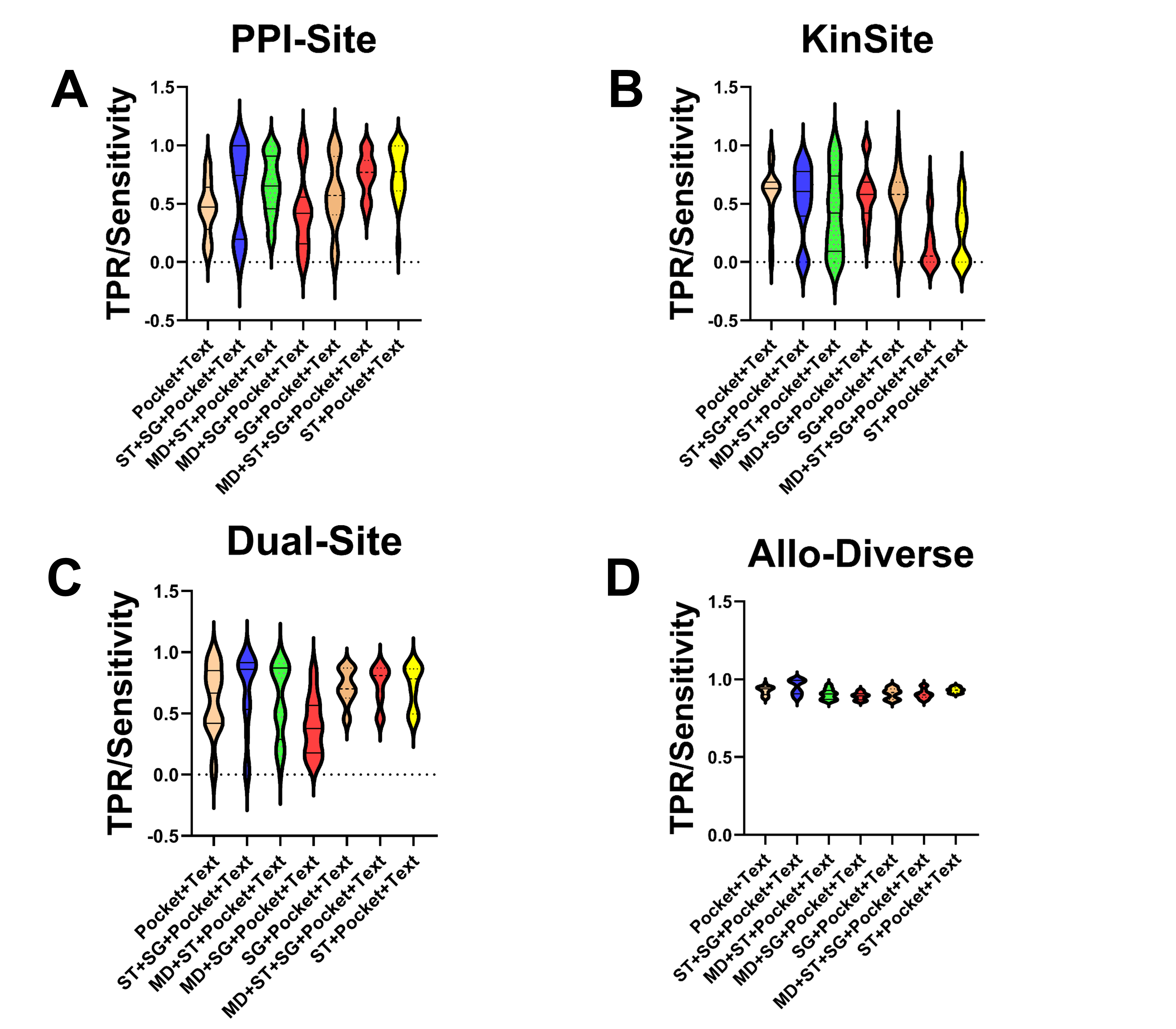

### FigureS5_JCIM_SUBMISSION.tif

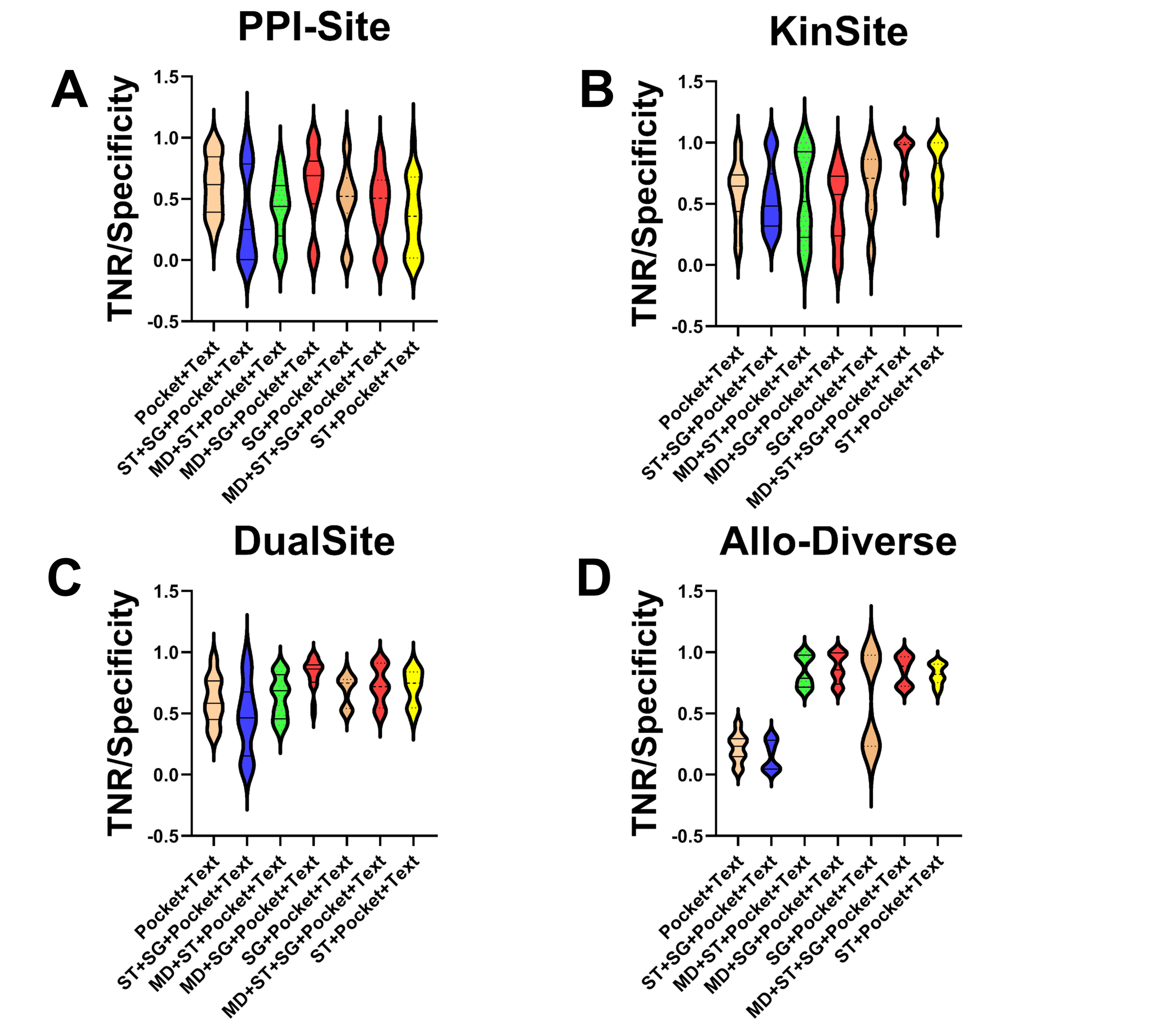
